# Circadian- and sex-dependent increases in intravenous cocaine self-administration in *Npas2* mutant mice

**DOI:** 10.1101/788786

**Authors:** Lauren M. DePoy, Darius D. Becker-Krail, Wei Zong, Kaitlyn Petersen, Neha M. Shah, Jessica H. Brandon, Alyssa M. Miguelino, George C. Tseng, Ryan W. Logan, Colleen A. McClung

**Affiliations:** Department of Psychiatry, Translational Neuroscience Program, University of Pittsburgh School of Medicine, 15219; Center for Neuroscience, University of Pittsburgh, 15261; Department of Biostatistics, University of Pittsburgh, 15261; Center for Systems Neurogenetics of Addiction, The Jackson Laboratory, 04609

**Keywords:** circadian, sex-differences, cocaine, self-administration, substance use, *Npas2*

## Abstract

Substance use disorder is associated with disruptions in circadian rhythms. The circadian transcription factor neuronal PAS domain protein 2 (NPAS2) is enriched in reward-related brain regions and regulates reward, but its role in substance use is unclear. To examine the role of NPAS2 in drug taking, we measured intravenous cocaine self-administration (acquisition, dose-response, progressive ratio, extinction, cue-induced reinstatement) in wild-type (WT) and *Npas2* mutant mice at different times of day. In the light (inactive) phase, cocaine reinforcement was increased in all *Npas2* mutants, while self-administration and motivation were affected sex-dependently. These sex differences were amplified during the dark (active) phase with *Npas2* mutation increasing self-administration, reinforcement, motivation, extinction responding and reinstatement in females, but only reinforcement in males. To determine whether circulating hormones are driving these sex differences, we ovariectomized WT and *Npas2* mutant females and confirmed that unlike sham controls, ovariectomized mutant mice showed no increase in self-administration. To identify whether striatal brain regions are activated in *Npas2* mutant females, we measured cocaine-induced ΔFosB expression. Relative to WT, ΔFosB expression was increased in D1+ neurons in the nucleus accumbens core and dorsolateral striatum in *Npas2* mutant females after dark phase self-administration. We also identified potential target genes that may underlie the behavioral responses to cocaine in *Npas2* mutant females. These results suggest NPAS2 regulates reward and activity in specific striatal regions in a sex and time of day specific manner. Striatal activation could be augmented by circulating sex hormones, leading to an increased effect of *Npas2* mutation in females.

**Significance Statement:** Circadian disruptions are a common symptom of substance use disorders and chronic exposure to drugs of abuse alters circadian rhythms, which may contribute to subsequent substance use. Diurnal rhythms are commonly found in behavioral responses to drugs of abuse with drug sensitivity and motivation peaking during the dark (active) phase in nocturnal rodents. Emerging evidence links disrupted circadian genes to substance use vulnerability and drug-induced alterations to these genes may augment drug-seeking. The circadian transcription factor NPAS2 is enriched in reward-related brain regions and regulates reward, but its role in substance use is unclear. To examine the role of NPAS2 in drug taking, we measured intravenous cocaine self-administration in wild-type and *Npas2* mutant mice at different times of day.

## Introduction

Circadian disruptions are a common symptom of many psychiatric disorders (Kowatch et al., 1992; Morgan & Malison, 2007; Matuskey et al., 2011), including substance use disorder (SUD)(Spanagel et al., 2005; McClung, 2007; Logan et al., 2014; Lauren M DePoy et al., 2017). Diurnal rhythms in the behavioral responses to drugs of abuse are common with drug sensitivity and drug use peaking during the active compared to inactive phase, or dark and light respectively in nocturnal rodents (Baird & Gauvin, 2000; Abarca et al., 2002; Roberts et al., 2002; Sleipness et al., 2005). Chronic exposure to drugs of abuse alters circadian rhythms, which may contribute to subsequent substance use (SU)(Roberts et al., 2002; Logan et al., 2014; Lauren M DePoy et al., 2017). Emerging evidence from rodents and humans links disrupted circadian genes to SU vulnerability (Dong et al., 2011; Forbes et al., 2012; Ozburn et al., 2012, 2013; Blomeyer et al., 2013; Gamsby et al., 2013; Bi et al., 2014; Baranger et al., 2016) and drug-induced alterations to these genes may augment drug-seeking.

Almost every cell in the brain and body expresses a molecular clock, comprised of several interlocking transcriptional-translational feedback loops (Reppert & Weaver, 2002; Ko & Takahashi, 2006; Takahashi, 2016). The molecular clock is regulated by Circadian Locomotor Output Cycles Kaput (CLOCK) or its homologue, neuronal PAS domain protein 2 (NPAS2), which dimerize with Brain and Muscle ARNT-like 1 (BMAL1) to control transcription of many genes. After translation, these proteins enter the nucleus and inhibit the transcriptional activity of (CLOCK/NPAS2)/BMAL1, closing the negative feedback loop (Ko & Takahashi, 2006). Our laboratory has previously demonstrated that mutations in *Clock* lead to increased cocaine and alcohol intake in mice (McClung et al., 2005; Ozburn et al., 2012, 2013) and a loss of diurnal rhythmicity in cocaine self-administration (Roberts et al., 2002). We have also shown that while *Clock* mutation increases cocaine preference, *Npas2* mutation attenuates preference (Ozburn et al., 2015a). NPAS2 is similar to CLOCK in structure and function, yet relative to CLOCK, NPAS2 expression and promotor binding is highly rhythmic in the striatum (Garcia et al., 2000a; Ozburn et al., 2015a), including the nucleus accumbens (NAc), a major neural substrate of reward (Wise & Rompre, 1989). In fact, in the NAc rhythms in *Npas2* are more highly affected by cocaine than *Clock* (Falcon et al., 2013) and *Npas2* knockdown increases glutamatergic transmission onto dopamine *Drd1*-expressing neurons (Parekh et al., 2019). Together, these findings suggest NPAS2 might play uniquely important role in the regulation of reward within the striatum.

This study aimed to determine whether NPAS2 regulates intravenous cocaine self-administration, a translational model of drug taking, the reinforcing and motivational properties of drugs and relapse-like behavior. We investigated whether *Npas2* mutation modulates the diurnal rhythm in drug taking by measuring self-administration during both the light and dark phase. We also examined whether *Npas2* mutation differentially impacts male and female mice since sex differences are prominent in circadian rhythms (Hatcher et al., 2018) and SUD (Becker & Hu, 2008). We find that NPAS2 regulates cocaine self-administration differentially across sex and time of day (TOD). Ovarian hormones and cocaine-induced expression of striatal ΔFosB are associated with increased self-administration in *Npas2* mutant females during the dark phase. Furthermore, we identify potential target genes that might underlie the behavioral responses to cocaine in *Npas2* mutant females.

## Methods and Materials

### Subjects

Male and female *Npas2* mutant mice or wild-type (WT) littermates, maintained on a C57BL/6J background, were used. These mice were originally described by Garcia et al., 2000. This mutation removes the bHLH domain of NPAS2, leaving the majority of the protein intact, but incapable of binding to BMAL1 (Garcia et al., 2000b). Adult mice were maintained on a 12:12 light-dark cycle with lights on (Zeitgeber Time (ZT0)) at 0700 or 1900. Behavioral testing occurred during the light phase from ZT2-7, unless specifically indicated as a dark phase experiment (ZT14-19). Food and water were provided *ad libitum* unless otherwise indicated. Procedures were approved by the University of Pittsburgh IACUC.

### Drug

Cocaine hydrochloride was provided by the National Institute on Drug Abuse. Animals were injected with 2.5, 5 or 15 mg/kg (intraperitoneal, i.p.; volume 10 ml/kg) in conditioned place preference and locomotor sensitization and 0-1 mg/kg/infusion for cocaine self-administration.

### Surgery

#### Jugular Catheterization

Mice were anesthetized with a 100 mg/kg ketamine / 1 mg/kg xylazine mixture. Surgery was performed under white light, during the second half of the inactive phase, regardless of light housing conditions for the mice. As previously described (Ozburn et al., 2012; Lauren M. DePoy et al., 2017), the dorsal and ventral sides were shaved and disinfected. The right jugular vein was exposed by blunt dissection and a sterile polyurethane catheter was placed and secured to the vein. The catheter is exteriorized posterior to the scapulae via a dacron mesh mount (Instech). The dorsal and ventral wounds were sutured and mice were pair housed for the duration, unless fighting or uneven sample sizes necessitated single housing. Mice recover for 6-7 days before intravenous self-administration training begins. Catheters were maintained by infusing catheters daily with 0.05 mL gentamicin (0.33 mg/mL) and heparinized saline (30 USP/mL) containing baytril (0.5 mg/kg). Catheter patency was tested approximately once per week using 0.05 mL brevital (3 mg/mL), mice that failed to lose muscle tone were excluded.

#### Ovariectomy

Ovariectomy (OVX) was performed as previously described (Heger et al., 2003). Female mice of at least 10 weeks of age were anesthetized with isoflurane. The ventral side was shaved and disinfected. The ovaries were located and either left intact or ligated and removed in sham and ovariectomized groups respectively. The abdominal wall was secured with absorbable sutures and the skin stapled. Mice were allowed to recover for at least 10 days before food training began. Due to previous isoflurane treatment mice were also anesthetized with isoflurane during the jugular catheterization. Following surgery, mice were moved from a 12:12 light cycle to a reverse light cycle (1900 on) to acclimate before behavioral testing. Mice were allowed to recover for at least 10 days, ensuring reduced levels of circulating sex hormones before food training began. Hormones should be entirely ablated before cocaine self-administration.

### Behavioral testing

#### Conditioned place preference (CPP)

As previously described (Ozburn et al., 2015a) female mice 8 weeks and older were first habituated to a testing room for 30 minutes. On day 1, a pre-conditioning test was conducted, wherein mice were placed in the center of a 3-chamber box. The outer two chambers were distinct with visual and tactile differences. Time in each chamber was recorded over the 20-minute session and any mice spending ≥50% (600 seconds) in one zone were excluded. On the subsequent 4 days mice were injected with either saline or cocaine and restricted to one side of the chamber. Saline was injected on days 2 and 4, and cocaine was injected on days 3 and 5 (2.5 or 5 mg/kg, 10 mL/kg, NIDA drug consortium). Here, a biased design was used, since ≥ 50% of mice showed a chamber bias during the pre-test, wherein the preferred chamber (>10% preference) was paired with saline or chambers were assigned pseudo-randomly if no side preference was found.

#### Locomotor sensitization

One cohort of animals were used to examine locomotor sensitization at least one month following CPP for cocaine. All testing was done in clear plexiglass test chambers (Kinder Scientific Smart Cage Rack System; field dimensions: 9.5” x 18.0”) equipped with infrared photobeams measuring horizontal locomotor activity. Before beginning each session, mice were allowed to acclimate to the test room for one hour. Briefly, the protocol began with 1 day of habituation to the test chamber and 2 days of 10 ml/kg saline injections (i.p.). Mice were then given 5 consecutive days of 15 mg/kg cocaine injections (i.p.). Following a 7-day withdrawal period, mice were given 2 consecutive challenge days of cocaine at the same dose. For all sessions, 60 minutes of locomotor activity was measured as distance traveled (cm), both in total and across 5-minute bins. Chambers were cleaned with 70% ethanol between animals.

#### Food self-administration

Mice were restricted to 85% of their free-feeding weight. Mice were trained to respond for chocolate flavored food pellets (20 mg, grain based precision pellets, Bio-Serv) in Med-Associates operant conditioning chambers. Responding on one lever was reinforced using a fixed ratio 1 (FR1) schedule. A cue light was illuminated over the active lever for the duration of the experiment. Responses on the inactive lever had no programmed consequences but were recorded. Sessions ended at 60 minutes or when the maximum of 30 pellets were acquired. Mice were trained for at least 5 sessions or until they acquired ≥25 pellets for 3 consecutive sessions.

#### Intravenous cocaine self-administration

After recovery from jugular catheterization, mice were trained to respond on an FR1 schedule for cocaine (0.5mg/kg/infusion, 30 µl over 1.7 seconds) on the previously inactive lever from food training (Ozburn et al., 2012). Cocaine was delivered through an armored tether connected to a swivel and syringe pump. Mice were tested 6 days/week with the last day being reserved for patency testing. Drug delivery culminated in extinction of the house light, a compound cue (auditory tone and stimulus light), and a 10-sec timeout during which no additional cocaine reinforcers can be delivered. Sessions ended when mice self-administered 60 infusions or at 60 min (first 8 sessions) or 120 min (subsequent sessions, 6 minimum). Mice were considered to have acquired the cocaine-reinforced response when mice self-administered ≥15 reinforcers across 3 sessions (with ≥ 2:1 active/inactive lever press ratio). Drug intake (infusions), discrimination of the active versus inactive and time to reach criteria were measured.

After acquisition, mice were tested on an FR1 schedule with 2 consecutive sessions of descending unit doses of cocaine (1.0, 0.5, 0.25, 0.125, 0.063, and 0 mg/kg). This dose-response analysis measured the reinforcing properties of cocaine. Next, mice self-administered cocaine at the baseline 0.5 mg/kg unit dose for two days before progressive ratio testing. The dose-response analysis is used to measure the reinforcing properties of cocaine, which will be described throughout as reinforcement. In order to measure motivation to take cocaine in the same mice, 3 counter-balanced unit doses (1.0, 0.5, and 0.25 mg/kg) were presented for 2 consecutive sessions under a progressive ratio schedule, wherein each successive infusion requires more lever press responses (1, 2, 4, 6, 9, 12, 16, 20…240). Sessions end after 4 hours or 1.5 hours without acquiring a reinforcer. This test is used to measure motivation, which will be used to describe changes to the breakpoint ratio, or the last ratio obtained for a dose of cocaine. Mice were again returned to their baseline self-administration dose (0.5 mg/kg/infusion) for two days before extinction training. Here, all lever presses had no programmed consequences and mice were trained for 10 days or until ≤30% peak active responding at 0.5 mg/kg/infusion was reached. Following extinction, mice were tested for sensitivity to cue-induced reinstatement, a model of relapse where presentations of previously cocaine-associated cues invigorate responding on the previously cocaine-reinforced lever. Mice received one non-contingent cue presentation, followed by cues contingent upon responding on the previously active lever.

Mice used for ΔFosB quantification self-administered cocaine (0.5 mg/kg/infusion) for 14 days as above. Here, in an attempt to normalize cocaine intake between WT and *Npas2* mutant mice, infusions were limited to 25, the approximate average number of infusions acquired by WT females during self-administration. After 25 infusions were obtained the session ended and mice were removed from the chambers.

### Dual RNAscope and immunohistochemistry

RNAscope *in situ hybridization* (ISH) for *Drd1* was followed by immunohistochemistry (IHC) for ΔFosB. The percent of dopamine D1 receptor+ and D1 receptor-(D1+,D1-) neurons expressing ΔFosB were measured since these populations differentially regulate reward (Lobo et al., 2010; Yawata et al., 2012; Smith et al., 2013). Mice were trained to self-administer cocaine and brains were flash frozen 24 hr after the last self-administration session (Fig.6a). Coronal sections (14 µm) were cut on a cryostat (Leica) and serial sections between 1.54 and 0.86 mm from bregma were immediately mounted onto Super Frost Plus slides. Each resulting slide contained 4 sections ranging along the anterior-posterior extent of the striatum. Slides with frozen sections were processed using an RNAscope fluorescent multiplex kit x (Advanced Cell Diagnostics #320850). Tissue sections were fixed at 4°C in 10% normal buffered formalin for 30 minutes and slides were rinsed three times 1X phospho-buffered saline (PBS) then dehydrated in a series of ethanol solutions at room temperature (RT)(50, 70, 100, 100% ethanol). We then hybridized the tissue sections with a *Drd1* probe (Advanced Cell Diagnostics 406491) and incubated slides at 40°C for 2 hr in the HybEZ oven. The signal was then amplified through incubation at 40°C with sequential amplifiers (AMP1 30 min, AMP2 15 min, AMP3 30 min) with AMP4 AltA used to detect *Drd1* in green (Alexa 488). Slides were washed twice in 1x wash buffer between incubations and then rinsed in PBS before immunohistochemical staining for ΔFosB (3 x 10 minutes).

Mounted sections were blocked for 1 hr at RT in 10% normal donkey serum and 0.3% 100x triton. Tissue was then incubated in a rabbit anti-FosB primary antibody (1:200, Cell Signaling Technology 2251) in block overnight at 4°C. After rinsing in PBS, sections were incubated in donkey anti-rabbit Alexa Fluor 555 secondary antibody (1:500, Thermo Fisher Scientific A-31572) in block at RT for 1.5 hr, rinsed again, counterstained with DAPI and cover-slipped with prolong gold. Throughout, sections were incubated by filling the hydrophobic barrier with approximately 150 µl of each solution. Although a pan-FosB antibody was used, full-length FosB degrades within 18-24 hr (Perrotti et al., 2008), therefore, all immunoreactivity should reflect ΔFosB.

### Imaging and cell counting

Immunofluorescence images [DAPI (blue), D1 (488 channel, green) and ΔFosB (546 channel, red)] were captured on an Olympus IX83 confocal microscope using a 20x objective at x2 magnification. Approximately 300 x 300 µm images were taken from sections along the anterior-posterior extent of the NAc core and shell, and dorsomedial (DMS) and dorsolateral (DLS) striatum. 5-6 2.5 µm z-stacks were used with an 8 µs/pixel dwell time and 1024 x 1024 pixel size. Consistent laser power, voltage and offset were used between images. Both hemispheres of 2-4 brain sections/region were imaged. Total D1+ and D1-cells as well as D1+ and D1-ΔFosB-expressing cells (see Fig.6c) were counted using ImageJ. The percentage of D1+ and D1-ΔFosB expressing cells was calculated for all conditions and brain regions. Image capture and quantification were performed by one blinded experimenter.

### RNA-sequencing

Brains were extracted and flash frozen from cocaine naïve, adult *Npas2* mutant or WT female mice in the dark phase at ∼ZT16, matching time of death after self-administration when ΔFosB expression was measured (Fig.6). Bilateral 1mm punches were taken centered over the NAc, DMS or DLS. Individual animals were each used as a sample (n=4-6 per condition). Tissue was homogenized and total RNA was isolated using the RNeasy Plus Micro Kit (Qiagen). RNA quantity and quality were assessed using fluorometry (Qubit RNA Broad Range Assay Kit and Fluorometer; Invitrogen) and chromatography (Bioanalyzer and RNA 6000 Nano Kit; Agilent) respectively. Libraries were prepared using TruSeq mRNA kit for stranded mRNA. Total RNA input was enriched for mRNA and fragments. Random primers initiate first strand and second strand Cdna synthesis. Adenylation of 3’ ends was followed by adapter ligation and library amplification with indexing. High Output, 400 M clustered flowcells (75 cycles) were sequenced with the NextSeq500 platform (Illumina) in 75bp single read mode. 30 M reads per sample was targeted. Data were filtered for non- and low-expressed genes. Differential expression (DE) between genotypes, Ingenuity pathway analysis (IPA) of enriched differentially expressed genes (DEGs) and rank-rank hypergeometric overlap (RRHO) between brain regions were performed.

### Statistical Analyses

GraphPad Prism 7 and IBM SPSS statistics were used. Data are expressed as means+SEM with *p*=0.05 considered significant and 0.05<*p*<0.1 considered trending. ANOVAs were performed with significant interactions followed by Bonferroni post-hoc tests corrected for multiple comparisons. Unpaired one-tailed t-tests were used to analyze factors such as total infusions (Fig.7) since we are expecting to replicate and then reverse the increase in self-administration seen in female *Npas2* mutants.

SPSS was used for analysis when additional factors were used: TOD and/or sex. Since light phase and dark phase animals cannot be tested concurrently due to light constraints, as well as the *a priori* hypothesis that mice would self-administer more cocaine during the dark phase, each phase was measured separately. Self-administration was analyzed with 3-way repeated measure (RM) ANOVAs across session (session x sex x genotype), while criteria, total infusions and ΔFosB were analyzed with 2-way ANOVAs (sex x genotype). Data are also represented for males and females separately to highlight any relevant sex differences, although these ANOVAs were not performed (and statistics are not reported) unless significant 3-way interactions were detected. Values ≥2 standard deviations from the mean were considered outliers and excluded.

For sequencing, non-expressed genes were first removed and then the 25% low-expressed genes were filtered out. 14,270 annotated genes remaining after filtering were used as a background data set. Differential expression (DE) analysis comparing WT and mutant mice was conducted by limma-voom in each brain region separately. Genes were considered DE with a loose p-value criteria of p<0.05 without multiple comparison correction. We used rank-rank hypergeometric overlap (RRHO) as a threshold-free method to evaluate the overlap of DE patterns across pairs of brain regions (Cahill et al., 2018). RRHO identifies overlap between two ranked lists of differential gene expression. The genes are ranked by the -log10(p value) multiplied by the effect size direction. To rank the overlapping genes from RRHO by their DE significance, we performed Adaptively-Weighted Fisher’s (AW-Fisher) (Li & Tseng, 2011; Huo et al., 2020) analysis to combine the DE results for DLS and NAc. This provides a meta-analyzed p-value for each gene with increased statistical power and generates weight indicators to reflect the consistency of DE signals in the two regions [*i*.*e*. (1,1) DE in both regions, (1,0) DE in DLS but not NAc, and (0,1) DE in NAc but not DLS]. Overlapping genes between DLS and NAc were then ranked by their AW-Fisher meta-analyzed p-values. Fisher’s exact test was used to test enrichment significance of DEGs in gene sets downloaded from http://ge-lab.org/gskb/. Pathways whose size are smaller than 3 or greater than 500 were not considered.

## Results

### *Npas2* mutant mice have altered behavioral responses to cocaine

To expand upon evidence that NPAS2 regulates the behavioral effects of cocaine, we examined the role of NPAS2 in a translational model of drug taking, intravenous cocaine self-administration. We measured behavior in male and female *Npas2* mutant mice during the light or dark phase, since NPAS2 regulates circadian rhythms (Ko & Takahashi, 2006; Takahashi, 2016).

Animals were first trained to respond for food and discriminate between the two levers over time [day x lever interactions: light (F_(4,440)_=435.04,p<0.0001), dark (F_(4,252)_=114,45,p<0.0001)]. A 4-way ANOVA revealed that response rates varied by genotype and TOD (session x genotype x TOD interaction: F_(4,173)_=4.19,p=0.002) and throughout, light and dark phase behavior were analyzed individually in order to identify TOD-specific effects. During the light phase, response rates tended to vary by genotype and sex (sex x genotype interaction: F_(1,110)_=2.92,p=0.09)(Fig.1abc). On the other hand, only genotype differences were found during the dark phase (main effect of genotype: F_(1,63)_=4.12,p=0.046)(Fig.1def) where mutants responded more on an active lever for food compared to WT mice.

**Figure 1.**
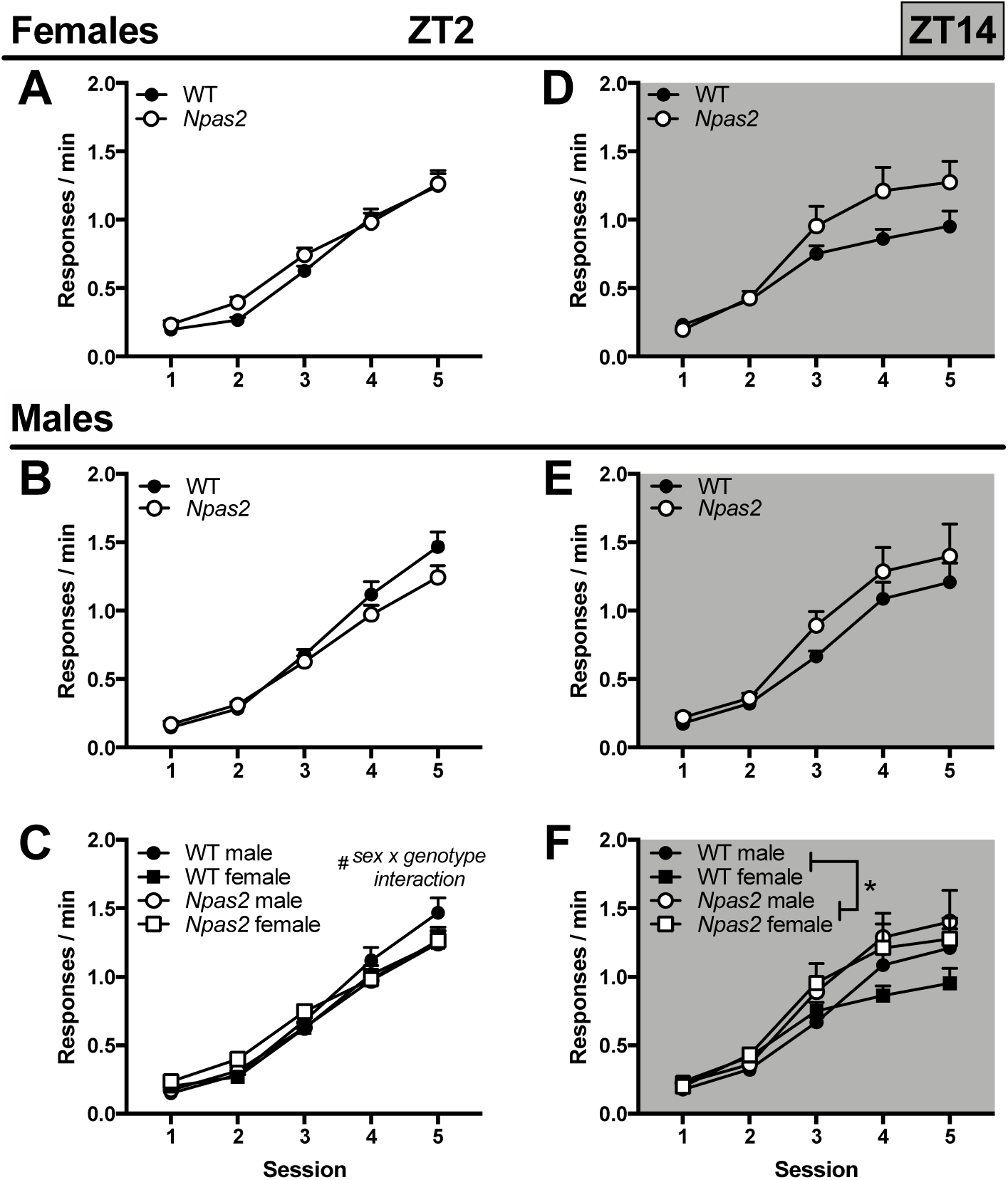
No major differences were found in food self-administration. *Npas2* mutant mice were trained to self-administer food pellets. (A) In the light phase, females are unaffected by *Npas2* mutation in the light phase, while (B) *Npas2* mutant males show a slight decrease in food responding compared to WT mice. (C) This was confirmed with a sex by genotype interaction in a 3-way ANOVA. During the dark phase, *Npas2* mutants responded more for food overall. (D) While this seems to be driven primarily by female and not (E) male mutants, (F) no sex difference was found. Means+SEMs, ^#^p<0.1; *p<0.05, *n*=16-18 dark phase; 25-36 light phase.

After recovery from jugular catheterization, mice were trained to self-administer cocaine. As expected, self-administration was higher during the dark than light phase in all mice, as shown by a main effect of TOD (F_(1,95)_=10.94,p<0.001). Overall, *Npas2* mutation differentially affects males and females (4-way ANOVA; sex x genotype: F_(1,95)_=4.18,p=0.044). Subsequent 3-way ANOVAs were used to investigate the effects of *Npas2* mutation across TOD. During the light phase, male and female (Fig.2ab) *Npas2* mutant mice self-administered more cocaine than WT mice (main effect of genotype: F_(1,56)_=15.98,p<0.001)(Fig.2c). On the other hand, cocaine intake varied by sex and genotype in the dark phase (sex x genotype interaction: F_(1,63)_=4.65,p=0.037)(Fig.2def). To further investigate these effects, we quantified sessions required to reach criteria and total drug intake (infusions). While all *Npas2* mutant mice acquired self-administration faster and took more infusions than WT mice in the light phase [main effect of genotype: criteria (F_(1,53)_=4.74,p=0.034), infusion (F_(1,56)_=16.1,p=0.0002)](Fig.3ab), sex differences emerged during the dark phase where only females mutants had a higher propensity to self-administer cocaine compared to WT females [sex x genotype interaction: criteria (F_(1,39)_=3.92,p=0.055), infusion (F_(1,39)_=3.12,p=0.085)](Fig.3cd). This finding demonstrates that female *Npas2* mutants indeed have increased self-administration during the dark phase, while males are unaffected.

**Figure 2.**
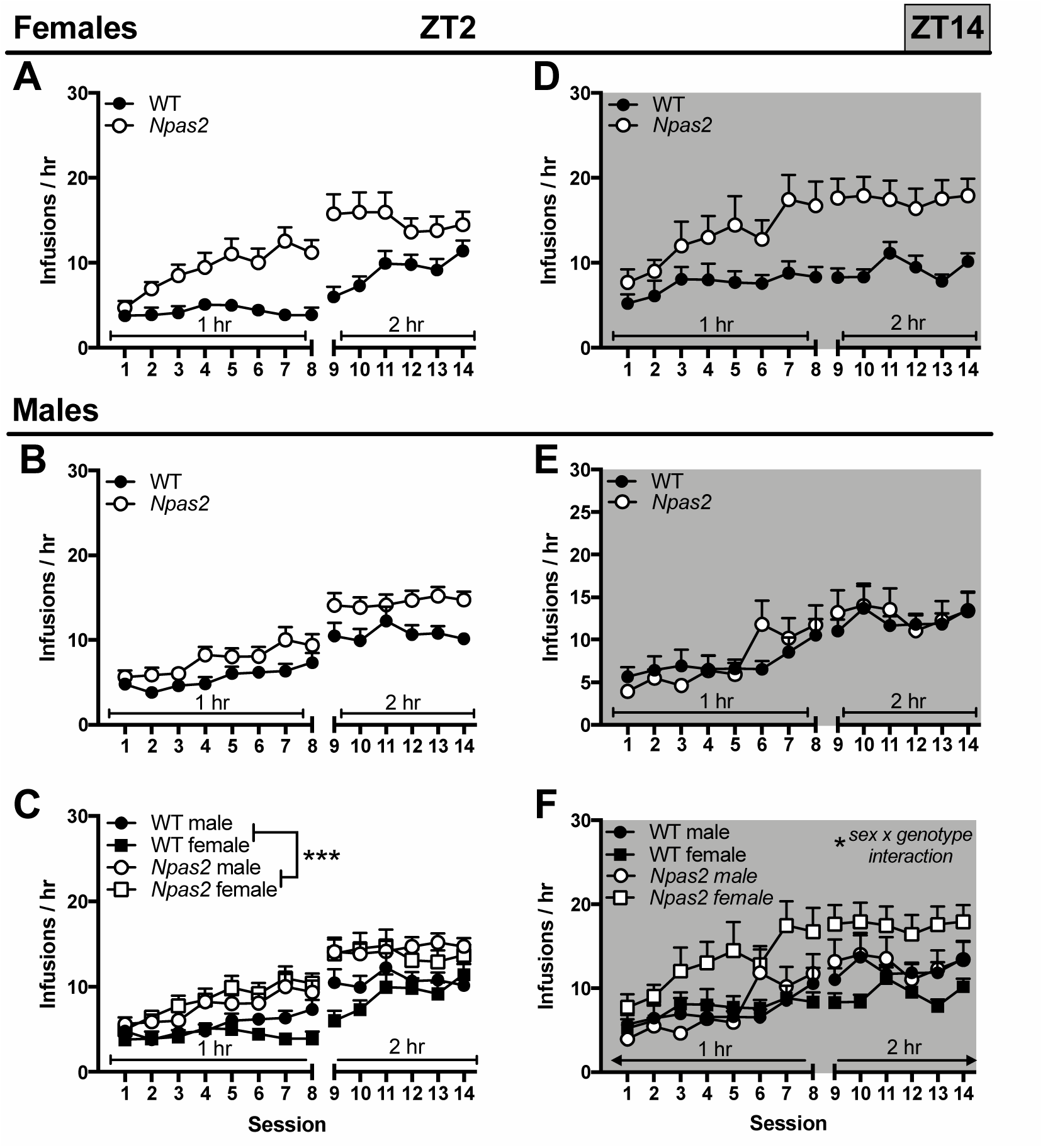
*Npas2* mutant mice self-administer more cocaine, particularly females in the dark phase. *Npas2* mutant mice were trained to self-administer cocaine (0.5mg/kg/infusion). (A) At ZT2, during the light phase, increased cocaine intake is seen in (A) female and (B) male *Npas2* mutant mice. (C) Although this effect seems to be particularly pronounced in female mice, overall no sex difference was found and all *Npas2* mutant mice self-administered more cocaine than WT mice. (D) In the dark, or active, phase (ZT14) cocaine intake appears to be elevated in female *Npas2* mutant mice compared to WT, (E) but not male mutants, (F) which is confirmed by a sex by genotype interaction. Means+SEMs, ^#^p<0.1; *p<0.05; **p<0.01; ***p<0.001, *n*=10-19.

**Figure 3.**
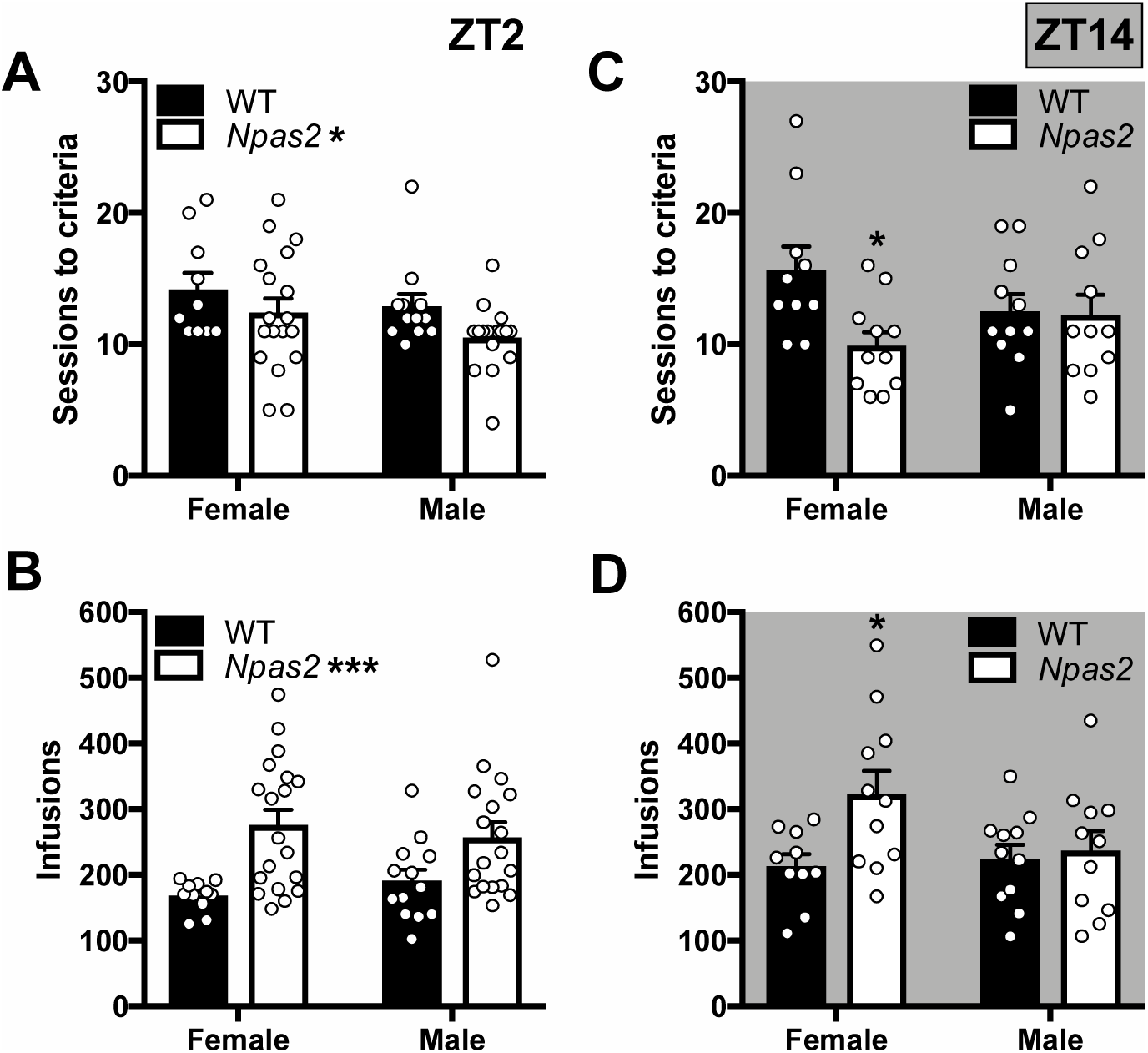
*Npas2* mutants have a greater propensity to self-administer cocaine. (A) An *Npas2* mutation increased the propensity to self-administer cocaine, decreasing the days required to reach acquisition criteria, during the light phase. (B) Total cocaine intake was also increased in mutant mice during the light phase. (C,D) In the dark phase, these effects were selective to females. Means+SEMs, *p<0.05; ***p<0.001, *n*=10-19.

In addition to analyzing cocaine intake directly using infusions, we also analyzed active and inactive lever pressing in order to determine whether *Npas2* mutant mice are taking more cocaine due to overall hyperactivity, which would lead to increases in both active and inactive lever pressing. A 5-way ANOVA revealed that all mice, both WT and mutant, discriminate between the active and inactive lever and respond differentially across TOD (session x lever x sex x TOD interaction: F_(13,1196)_=1.99,p=0.019). Subsequently, 4-way ANOVAs were used to investigate the effects of *Npas2* mutation in the light and dark phase separately. During the light phase, active versus inactive lever pressing varied by genotype (day x lever x genotype interaction: F_(13,637)_=3.75,p<0.0001), while only overall lever pressing, regardless of status, varied by sex (sex x genotype interaction: F_(1,49)_=12.16,p=0.001), suggesting discriminative lever pressing doesn’t vary by sex. Therefore, we excluded sex as a factor and found that similarly to cocaine intake, mutants pressed the active lever more than WT mice (session x genotype interaction: active lever pressing F_(13,663)_=3.48,p<0.0001)(Fig.4c). On the other hand, inactive lever pressing was only slightly increased in *Npas2* mutants compared to WT mie (trending main effect of genotype: inactive lever pressing F_(1,51)_=3.84,p=0.06). Importantly, *Npas2* mutants showed similarly significant discrimination between the active and inactive lever (session x lever interaction: F_(13,468)_=35.02,p<0.0001) as WT mice (session x lever interaction: F_(13,858)_=66.81,p<0.0001).

**Figure 4.**
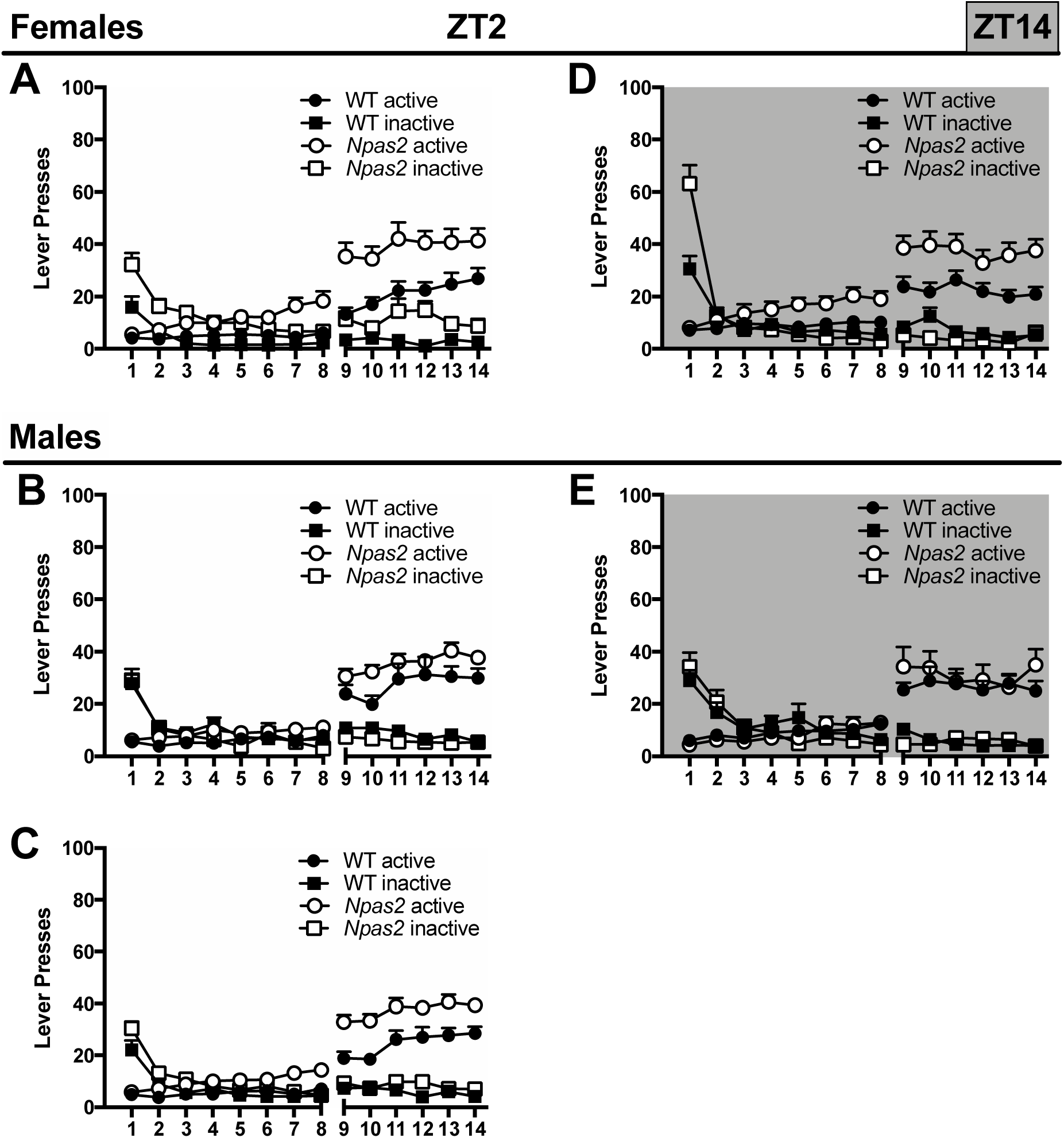
*Npas2* mutant mice respond more on an active lever for cocaine, particularly females in the dark phase. (A) At ZT2, during the light phase, increased active lever pressing for cocaine is seen in (A) female and (B) male *Npas2* mutant mice. (C) Although this effect seems to be greater in female mice, no sex difference was found. All *Npas2* mutant mice pressed the active lever more than WT mice (no posthocs shown), while only a trending increase in inactive lever pressing was found. Both WT and *Npas2* mutants press the active lever more than the inactive lever. (D) In the dark, or active, phase (ZT14) active lever pressing is elevated in female *Npas2* mutant mice (no posthocs shown), (E) but not male mutants. Both WT and *Npas2* mutants press the active more than the inactive lever. Means+SEMs, *n*=10-19.

During the dark phase, a 4-way interaction further emphasizes that *Npas2* mutation differentially affects males and females (session x lever x sex x genotype interaction: F_(13,559)_=2.59,p=0.002). Specifically, female mutants pressed the active lever more than WT mice (session x genotype interaction: active lever pressing F_(13,260)_=2.40,p=0.0046)(Fig.4d), while male mutants show no differences (Fig.4e). Although inactive lever pressing is also changed in *Npas2* mutants females when compared to WT mice (session x genotype interaction: inactive lever pressing F_(13,260)_=8.87,p<0.0001), responding is only increased on the first day of training, suggesting this increase is caused by perseveration on the inactive lever, which was previously reinforced with food and not hyperactivity. As with the light phase, all mice discriminated between the active and inactive levers [session x lever interaction: WT mice (F_(13,338)_=22.4,p<0.0001), *Npas2* mutants (F_(13,260)_=20.37,p<0.0001)].

To further explore the effect of *Npas2* mutation on reward, we measured CPP and locomotor sensitization. We previously found that *Npas2* mutant mice have reduced CPP compared to WT (Ozburn et al., 2015a), but only males were tested. Here we conditioned male and female mice to 2.5 or 5 mg/kg cocaine and found that sex plays a pivotal role in CPP [sex x genotype interaction: 2.5 mg/kg (F_(1,58)_=4.4,p=0.04), 5 mg/kg (F_(1,57)_=7.01,p=0.01)]. Although, we were able to replicate our previous finding that cocaine preference is reduced in male Npas2 mutants compared to WT controls (Ozburn et al., 2015b), we interestingly found that female mutants showed no change in preference (Fig.5a). We then confirmed that the lack of change in preference in mutant females is not due to TOD differences by measuring cocaine CPP during the dark phase. Again, cocaine preference was still unchanged in female *Npas2* mutants (Fig.5a).

**Figure 5.**
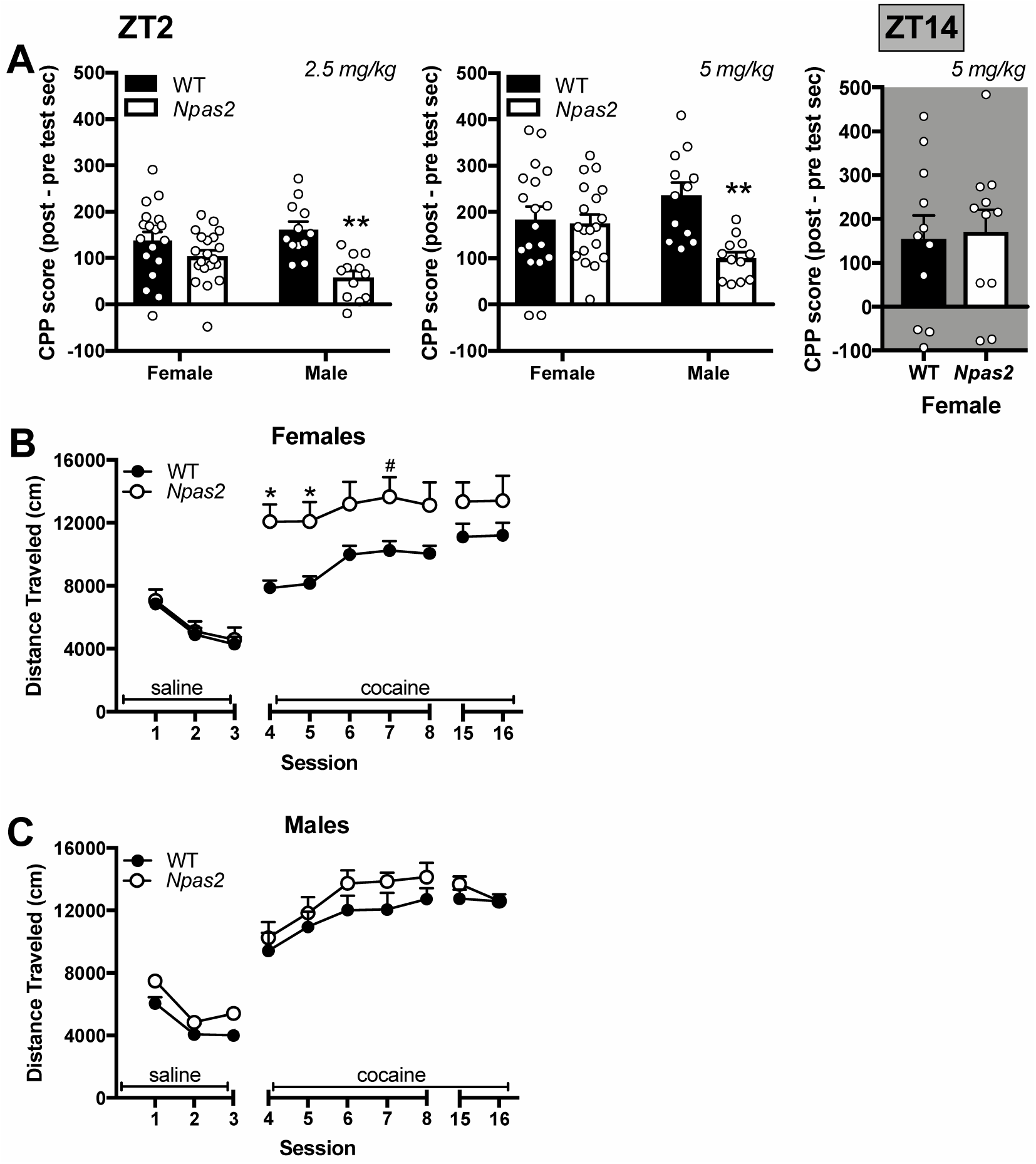
Sex differences in cocaine sensitization and place preference in *Npas2* mutant mice. (A) Male *Npas2* mutant mice showed a decrease in conditioned place preference to cocaine (2.5 and 5 mg/kg), whereas females showed no change. (Right) Cocaine conditioned place preference is also unchanged in the dark phase in female *Npas2* mutants. (B) One cohort of mice were then tested for locomotor sensitization. All mice showed sensitization to cocaine, but female *Npas2* mutant mice also showed elevated locomotor-activating effects of cocaine. (C) No effect of genotype was seen in male mice. Means+SEMs, ^#^p<0.1; *p<0.05; **p<0.01, *n*=12-20.

**Figure 6.**
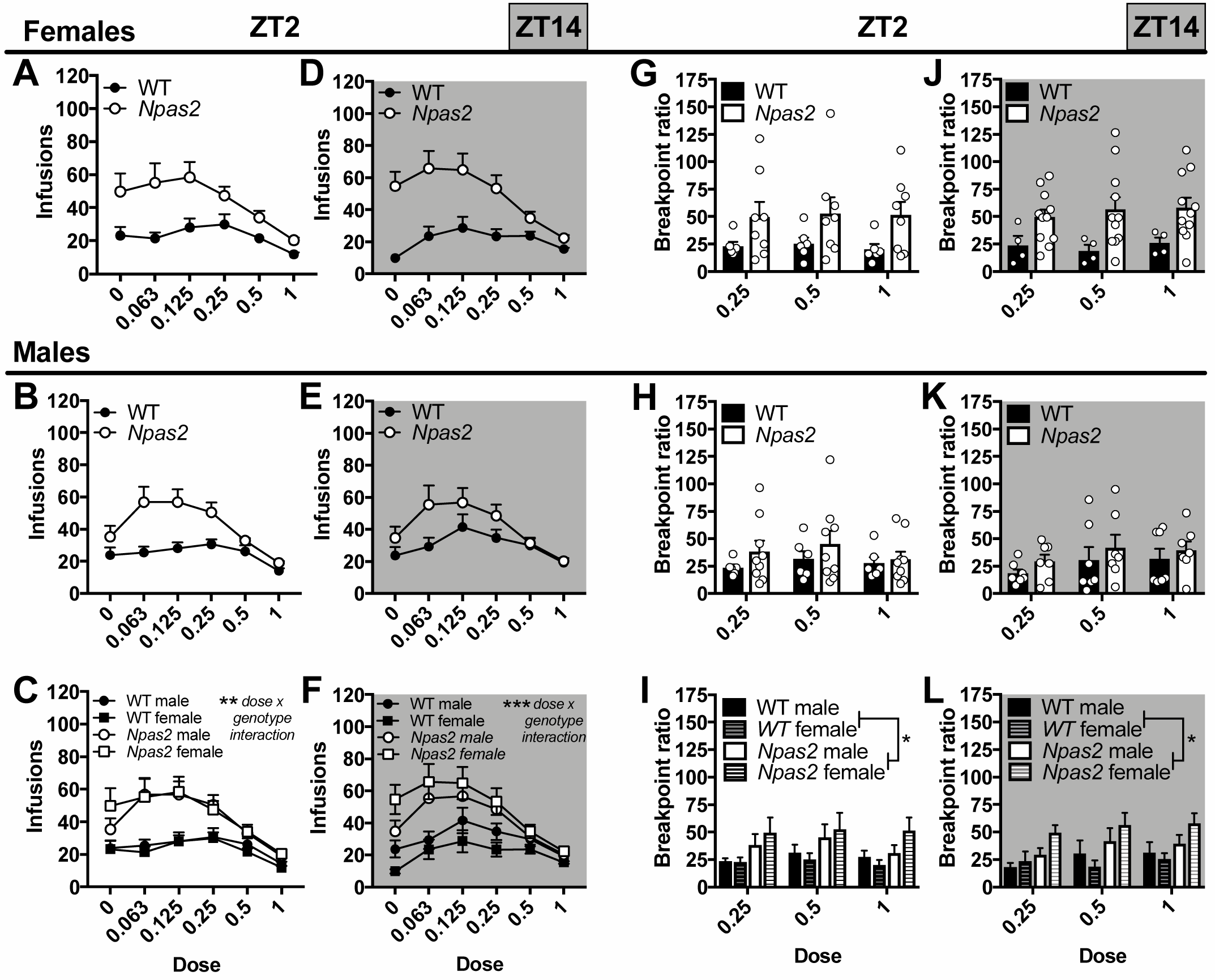
The reinforcing and motivational properties of cocaine were increased in *Npas2* mutant mice. During a dose-response analysis (0-1 mg/kg/infusion) at ZT 2 (light phase), *Npas2* mutant mice self-administered more infusions of cocaine across dose in both (A) female and (B) male *Npas2* mutant mice. (C) This significant increase in cocaine intake across sex suggests an increase in the reinforcing properties of cocaine. At ZT4 the reinforcing properties of cocaine were also increased in (D) female and (E) male mutant mice. Here, effects appear to be greater in female mutants, but (F) no sex effect was found. During progressive ratio testing, (G) female and (H) male *Npas2* mutant mice again worked harder for each infusion of cocaine. (I) Although a significant increase in breakpoint ratio was found across sex, this effect seems to be driven primarily by female mutant mice. Similar results are found during the dark phase, wherein breakpoint ratio was increased in (J) female and (K) male *Npas2* mutants. (L) Again, female mutants appear to be particularly affected, but no significant effect of sex was found. Means+SEMs, *p<0.05; **p<0.01; ***p<0.001, *n*=4-11.

During locomotor sensitization, we found that all mice sensitized to cocaine across sessions (main effect of session: F_(9,288)_=136.43,p<0.001), but the locomotor-activating effects of cocaine were dependent on sex and genotype (session x sex x genotype interaction: F_(9,288)_=2.54,p=0.008). When analyzed independently, female *Npas2* mutant mice showed an increase in the locomotor activating effects of cocaine compared to controls (session x genotype interaction: F_(9,126)_=2.06,p=0.038)(Fig.5b), while male mice showed a sensitization effect over multiple days (main effect of session: F_(9,162)_=99.76,p<0.001)(Fig.5c). These results further support the critical role of sex in how NPAS2 regulates the behavioral effects of cocaine. Importantly, hyperactivity does not appear to contribute to the differences observed in *Npas2* mutants as locomotor activity is not increased in male or female *Npas2* mutants following saline injections when compared to WT mice. These findings mirror our previous results demonstrating that locomotor response to novelty is not increased in male mutants (Ozburn et al., 2015a).

### Increased reinforcing and motivational properties of cocaine in *Npas2* mutant mice

Following acquisition, we measured the reinforcing properties of cocaine using a dose-response analysis. Throughout, *Npas2* mutant mice took more infusions compared to controls, indicating an overall increase in the efficacy of cocaine (dose x genotype interaction: F(5,345)=10.01,p<0.0001), which did not differ regardless of TOD or sex. Similar effects were seen during light phase (dose x genotype interaction: F_(5,180)_=3.98,p=0.002)(Fig.6c) and dark phase self-administration (dose x genotype interaction: F_(5,165)_7.51,p<0.001)(Fig.6f). Next, a progressive ratio schedule was used to measure motivation (breakpoint ratio), but a 4-way ANOVA only revealed a main effect of genotype (F(1,50)=7.90,p=0.007). This was confirmed across TOD, where *Npas2* mutant mice worked harder for each infusion of cocaine than WT controls, as shown by a higher breakpoint ratio during the light phase (main effect of genotype: F_(1,25)_=4.92,p=0.036)(Fig.6i) and dark phase (main effect of genotype: F_(1,25)_=5.14,p=0.032)(Fig.6l).

### Increased extinction responding, and cue-induced reinstatement in female *Npas2* mutant mice in the dark phase

Mice then underwent extinction and cue-induced reinstatement to model relapse-like drug seeking in the absence of cocaine. All mice extinguished responding (main effect of session: F(9,432)=28.68,p<0.0001) and we found a trending 4-factor interaction (session x sex x genotype x TOD interaction: F(9,432)=1.85,p=0.058) indicating divergent effects of genotype on males and females in the light and dark phase. We found a trend for *Npas2* mutation to increase extinction responding during the light phase (main effect of genotype: F_(1,23)_=3.06,p=0.09)(Fig.7c), with small increases in responding in female (Fig.7a) and male mutants (Fig.7b). However, during the dark phase, a 3-way interaction was found (session x sex x genotype interaction: F_(9,225)_=2.65,p=0.006)(Fig.7f) indicating divergent effects of genotype on male and female mice. Specifically, female *Npas2* mutants showed an increase in early extinction responding in the dark phase (session x genotype interaction: F_(9,108)_=2.71,p=0.007)(Fig.7d) compared to controls, while male mutants showed no differences (Fig.7e).

**Figure 7.**
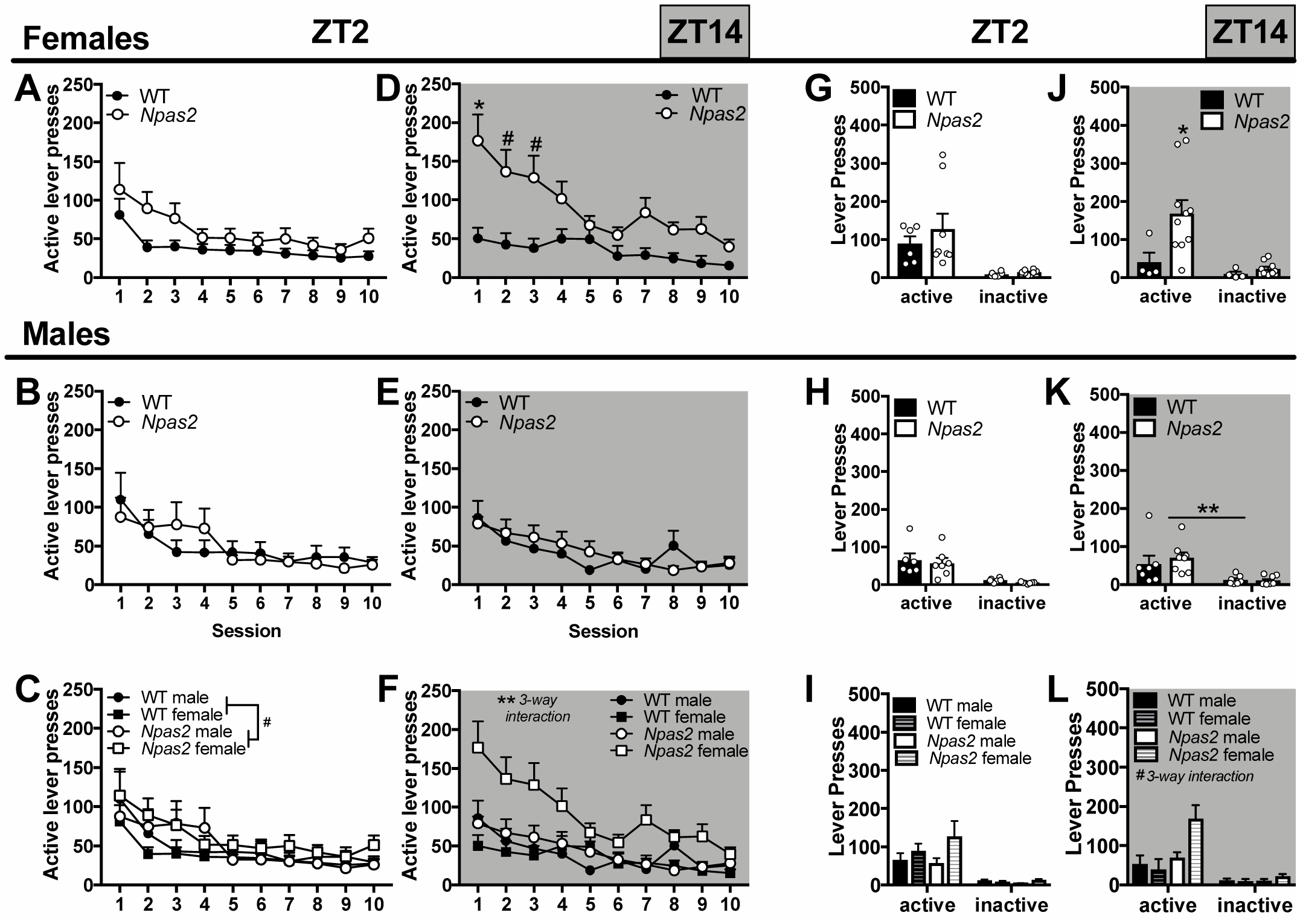
Increased extinction responding and cue-induced reinstatement in female *Npas2* mutant mice during the dark phase. Following progressive ratio, responding on the cocaine-associated lever was extinguished over the course of at least 10 days. Following extinction, responding on the active lever was reinstated with the presence of previously cocaine-associated cues. We found a very slight increase in extinction responding during the light phase in (A) female and (B) male mutant mice, as indicated by a (C) trending effect of genotype. On the other hand, during the dark phase, extinction responding was only increased in (D) female *Npas2* mutants, while (E) male mutants were unaffected. (F) This sex difference was confirmed by an interaction between session, sex and mutation. A similar pattern was detected for cue-induced reinstatement, wherein (G-I) no effects were found during the light phase, but (J) female and (K) male mice were differentially affected by *Npas2* mutation in the dark phase. (L) A trending interaction was found and female *Npas2* mutants responded significantly more during cue-induced reinstatement, suggesting increased drug seeking, but no differences were observed in males. Means+SEMs, ^#^p<0.1; *p<0.05; **p<0.01, *n*=4-11.

After extinction, contingent, previously cocaine-associated cues were presented, which reinstated responding in all mice (main effect of lever: F_(1,48)_=53.38,p<0.0001). Although no TOD differences were found in a 4-way ANOVA, *Npas2* mutation differentially affected males and females (sex x genotype interaction: F_(1485)_=4.49,p=0.039. In subsequent analyses, *Npas2* mutation did not affect cue-induced reinstatement during the light phase (Fs<1) in female or male mice (Fig.7ghi). Similar to extinction, drug seeking during reinstatement depended on sex and genotype during the dark phase (trending lever x sex x genotype interaction: F_(1,25)_=2.92,p=0.10)(Fig.7l). Follow-up ANOVAs indicate that female *Npas2* mutants showed an increase in drug seeking (lever x genotype interaction: F_(1,12)_=4.58,p=0.05)(Fig.7j) compared to WT mice, while male mutants showed no differences (Fig.7k). Together, these findings indicate that in the absence of cocaine, initially or with cocaine-associated cues present, female *Npas2* mutants intensify their drug seeking, specifically in the dark phase.

### Circulating sex hormones contribute to increased cocaine self-administration in female *Npas2* mutant mice

In order to determine whether ovarian hormones contribute to sex differences in self-administration in *Npas2* mutant mice, a separate cohort of females underwent an OVX or sham surgery before dark phase cocaine self-administration. As expected, we found that self-administration varied based on genotype and OVX (trending session x genotype x OVX interaction: F_(13,429)_=1.62,p=0.077). While sham mutant females showed moderately increased cocaine self-administration compared to sham WT females (main effect of genotype: F_(1,18)_=4.09,p=0.058)(Fig.8a), no effect was found in OVX WT and mutant mice (Fs<1)(Fig.8b). Furthermore, total drug intake was slightly increased in mutant sham compared to WT sham females (t_(18)_=1.63,p=0.059)(Fig.8c), but not mutant OVX compared to WT OVX females (t<1)(Fig.8d). These findings suggest that sex hormones mediate the greater effects of *Npas2* mutation seen in female mice.

**Figure 8.**
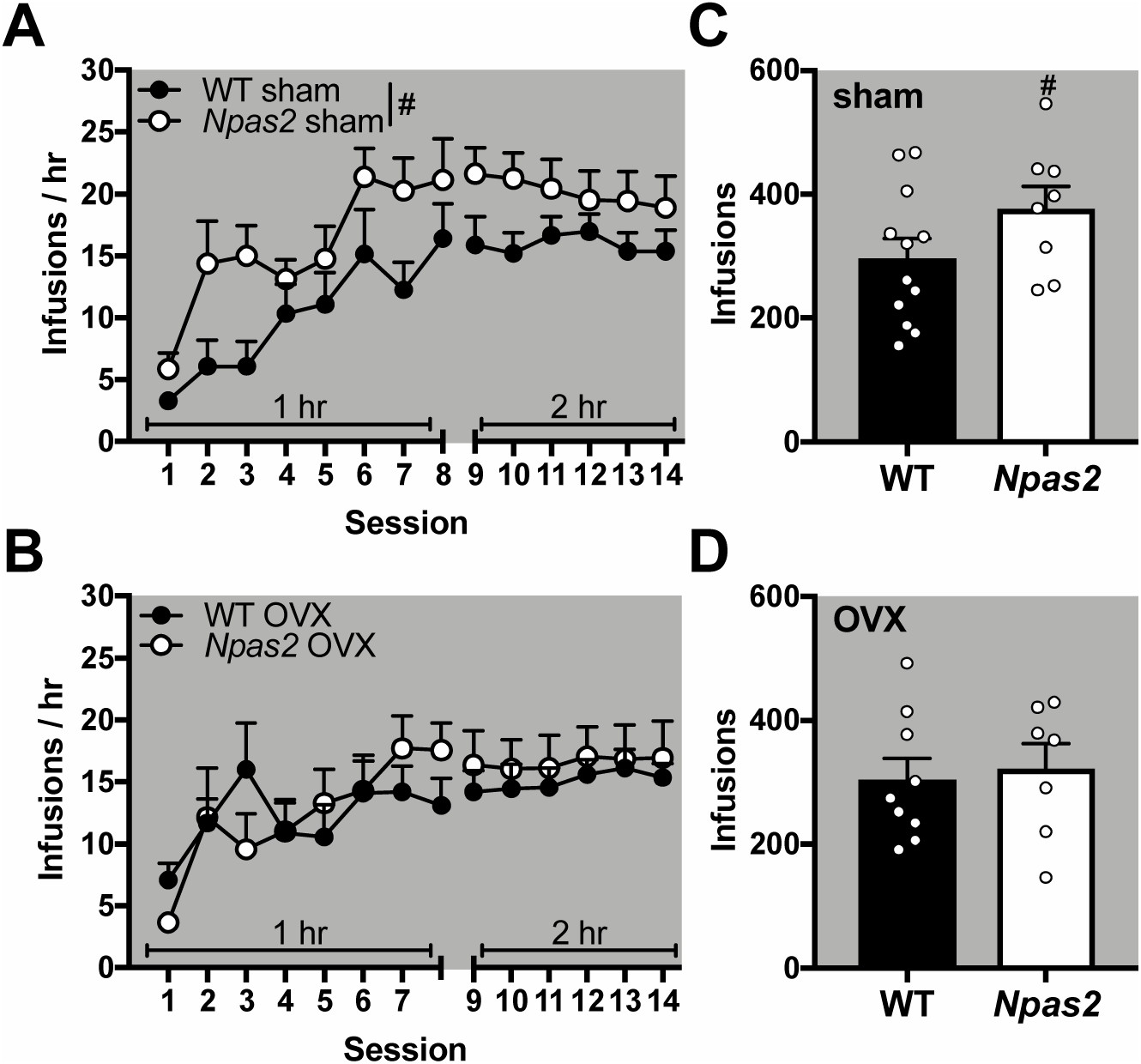
Ovariectomy reversed increased cocaine self-administration in *Npas2* mutant females in the dark phase. A separate cohort of female mice underwent sham surgery or ovariectomy (OVX) before being trained to self-administer intravenous cocaine. (A) Sham surgery treated mice recapitulated our original finding that *Npas2* mutant females take more cocaine than their WT counterparts during acquisition. (B) On the other hand, OVX seemed to reverse this effect with no differences seen between mutant and WT mice. These results were confirmed by total drug intake (infusions), which are (C) moderately increased in sham treated *Npas2* mutant mice, but not (D) OVX mice. Means+SEMs, ^#^p<0.1; *p<0.05, *n*=7-12.

### Increased ΔFosB expression in D1+ neurons in *Npas2* mutant females following dark phase cocaine self-administration

In order to determine which striatal regions might mediate increased self-administration in *Npas2* mutant females, we measured cocaine-induced expression of ΔFosB, a stable, long-lasting variant of FosB (Robison et al., 2013). Female mice self-administered cocaine during the light or dark phase. Mice were limited to 25 infusions in order to normalize acquisition [main effect of genotype: light (F_(1,9)_=2.73,p=0.133), dark (F<1); genotype x session interaction: light (F<1), dark (F_(13,117)_=2.23,p=0.012, no significant post-hocs)] between WT and *Npas2* mutant mice (Fig.9a). Tissue was harvested 24 hr after the last self-administration session.

We quantified the percentage of D1+ and D1-cells expressing ΔFosB in the NAc core, NAc shell, DLS and DMS (Fig.9b). No genotype differences were found in ΔFosB expression after light phase self-administration, but dark phase *Npas2* mutant females had slightly increased ΔFosB expression in the NAc shell (main effect of genotype: F_(1,9)_=4.16,p=0.072) compare to WT females. In both the NAc core and DLS, this increase in ΔFosB was specific to D1+ cells [cell x genotype: NAc core (F_(1,8)_=3.97,p=0.082), DLS (F_(1,10)_=5.64,p=0.039)]. No effects were seen in the DMS. Throughout, ΔFosB expression was higher in D1+ compared to D1-cells [main effect of cell-type: NAc (F_(1,18)_=30.47,p<0.0001), DS (F_(1,19)_=27.66,p<0.0001)].

### Most differentially expressed genes in *Npas2* mutant females are in the DLS and many are ΔFosB targets

We next aimed to identify possible mechanisms that could be driving increased dark phase cocaine self-administration in female *Npas2* mutant mice. Since drug taking is increased early in self-administration (Fig.2d), we believe predispositions exist in female mutants that drive this increase. Given regional differences in cocaine-induced striatal activation in female mutants, we identified DEGs in the NAc, DLS and DMS of cocaine-naïve WT and *Npas2* mutant females in the dark phase (Table.1-3). Using cut-offs of p<0.05 (uncorrected) and fold change (FC)>1.3, we found 343 DEGs in the NAc, 362 in the DMS and 922 in the DLS (Fig.9d). Due to the leniency of this p-value, some false positives are expected and fewer DEGs were found at more stringent cut-offs (Fig. 9d). Striatal regions that are similarly activated after self-administration in *Npas2* mutants show parallel changes in gene expression. The NAc and DLS, where ΔFosB expression is increased in mutant D1+ neurons, show a high level of overlap in DEGs (Fig.9e). However, the NAc and DLS show very little overlap with the DMS, where ΔFosB expression is not increased (Fig.9efg).

**Table 1.**
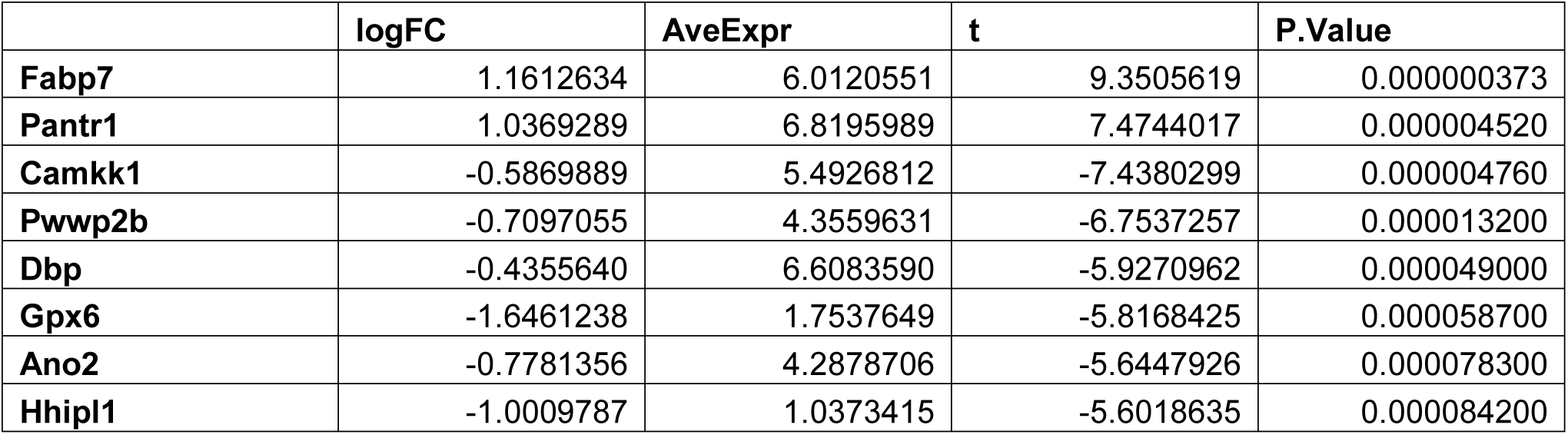
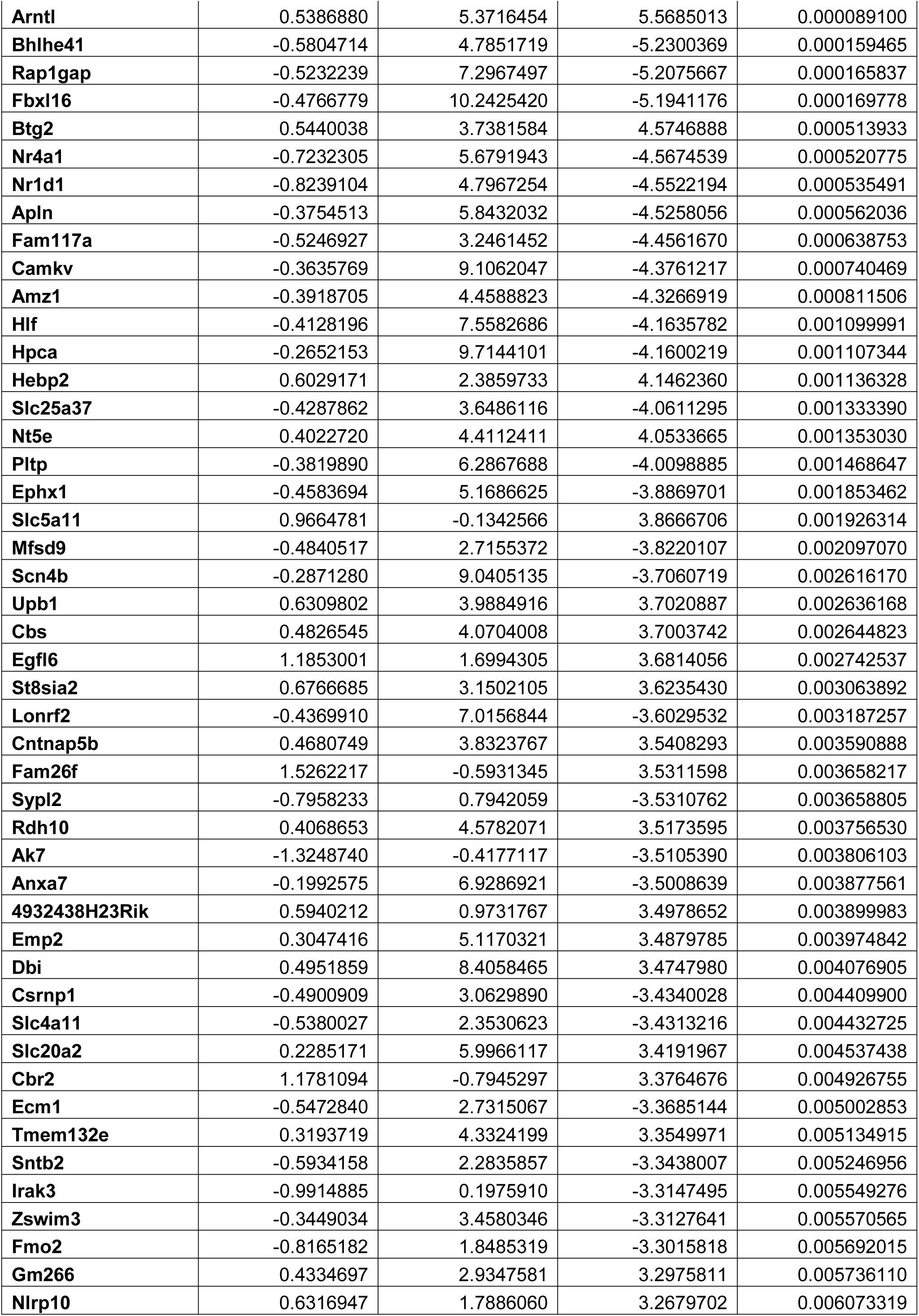
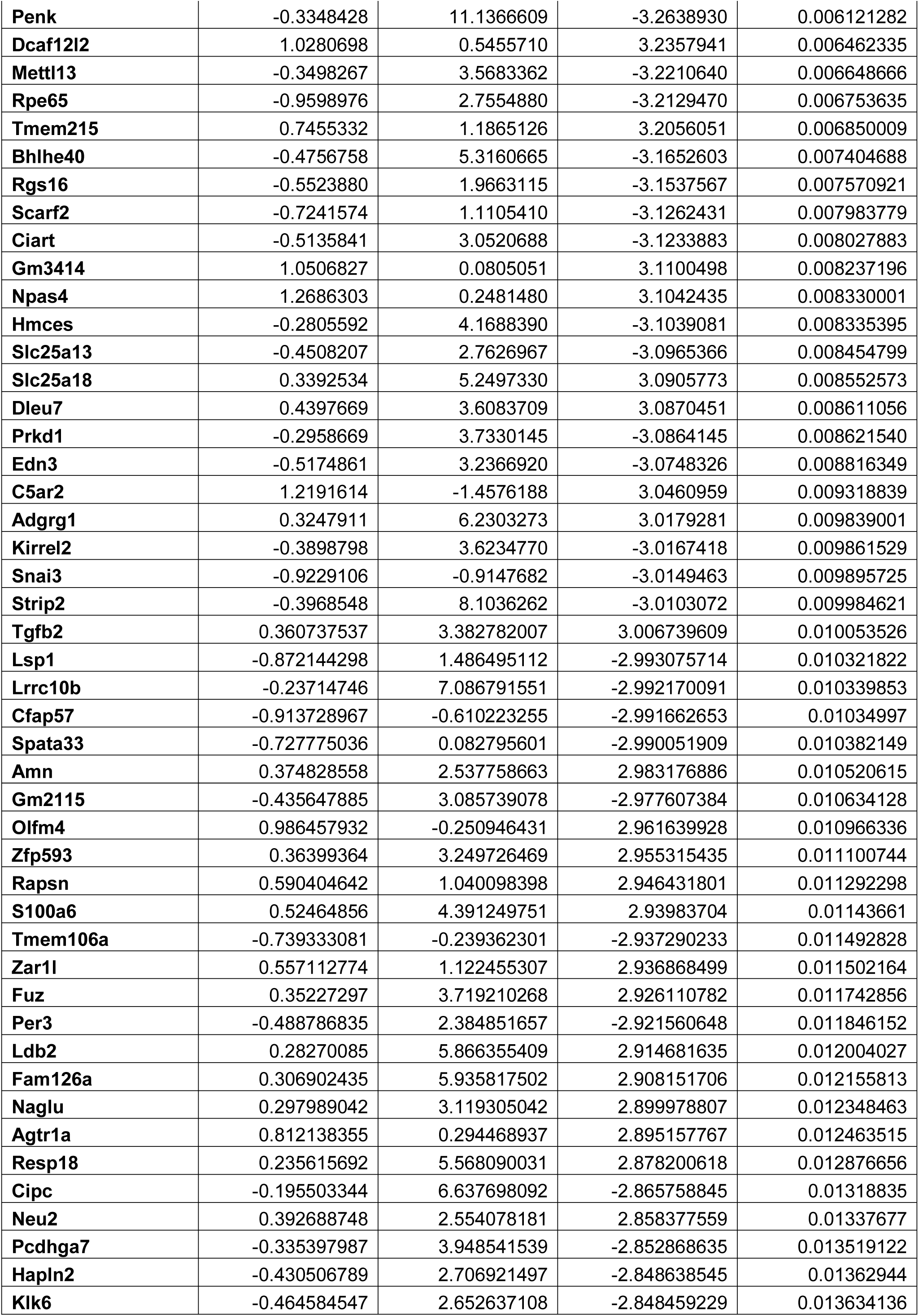
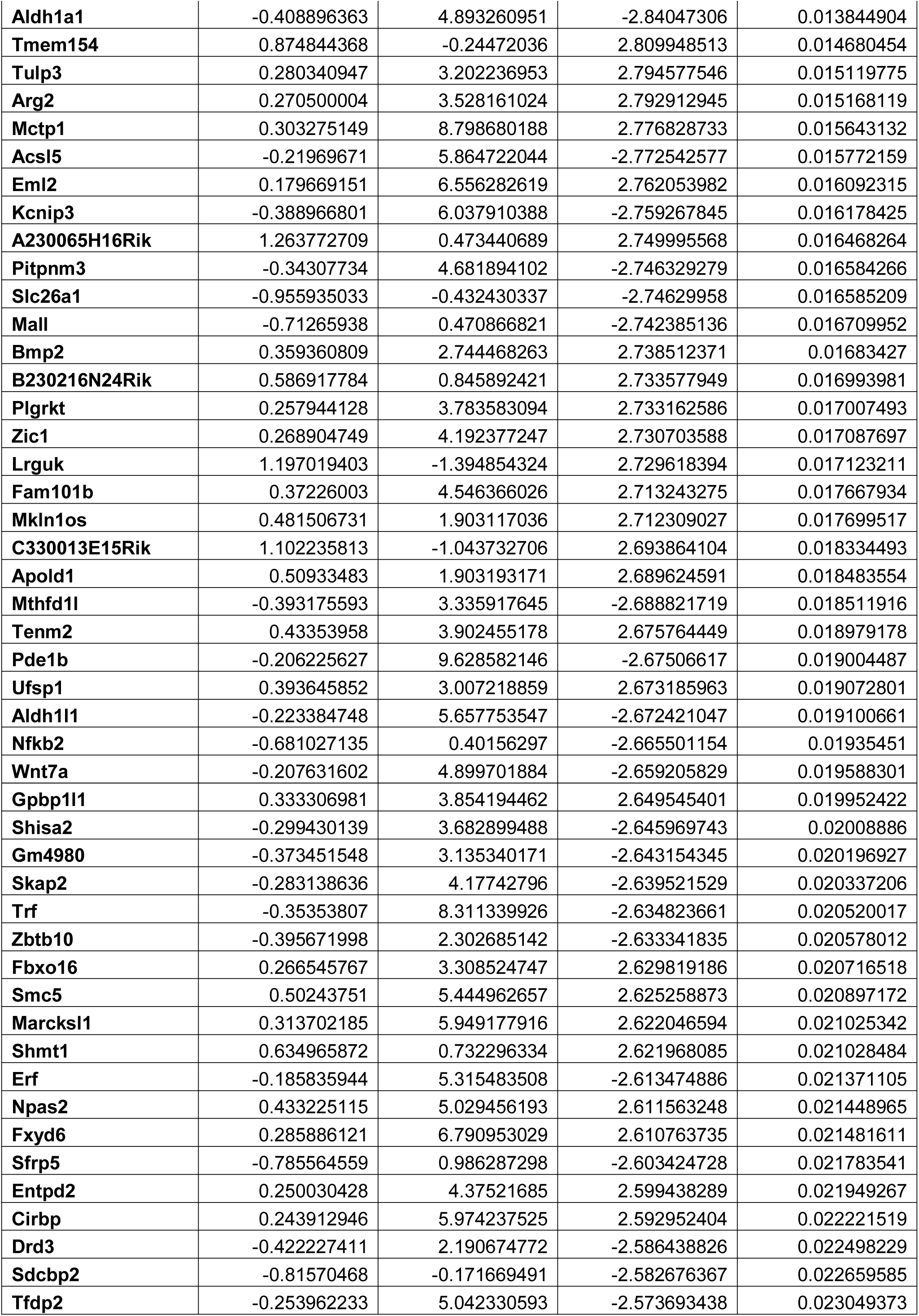
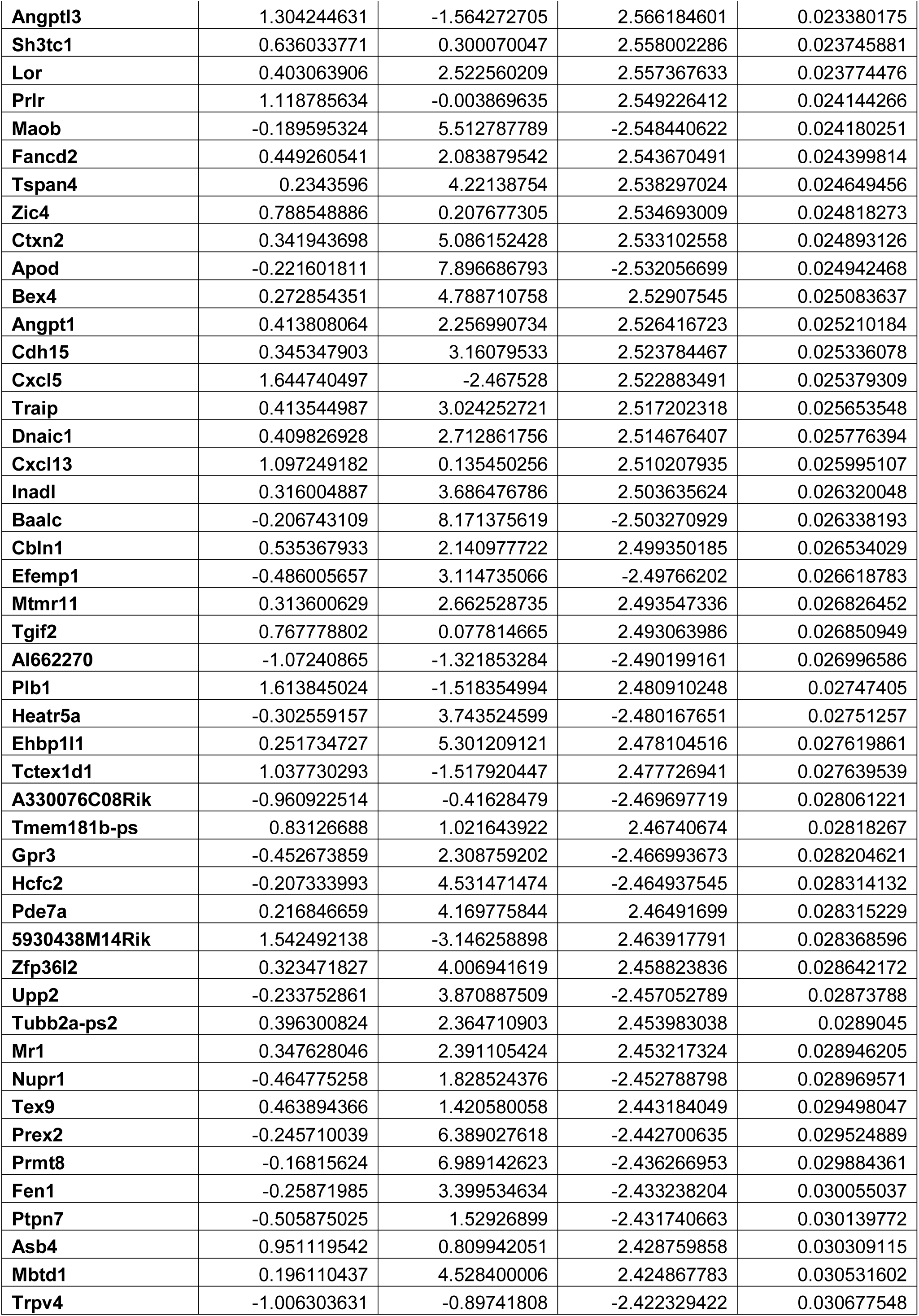
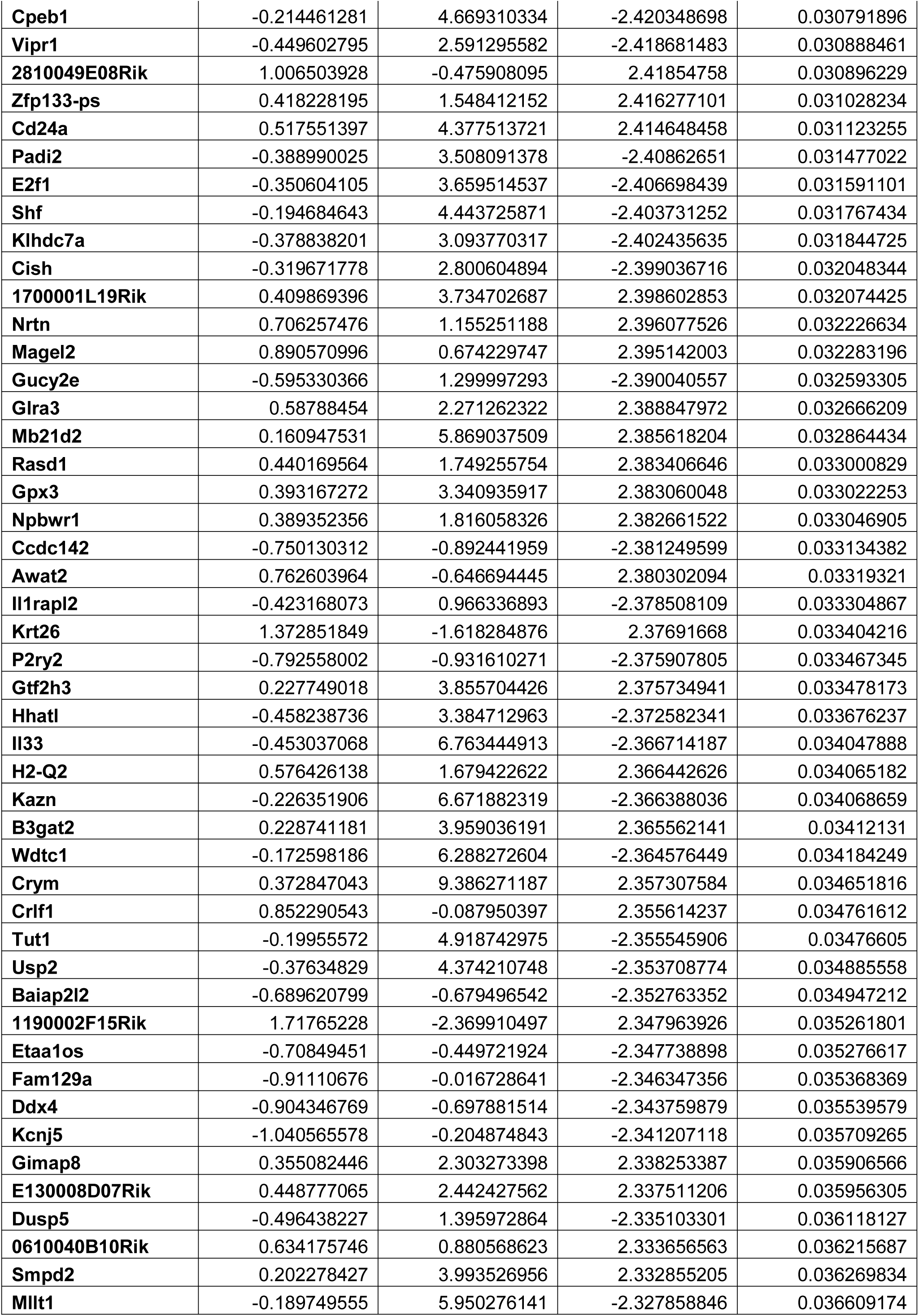
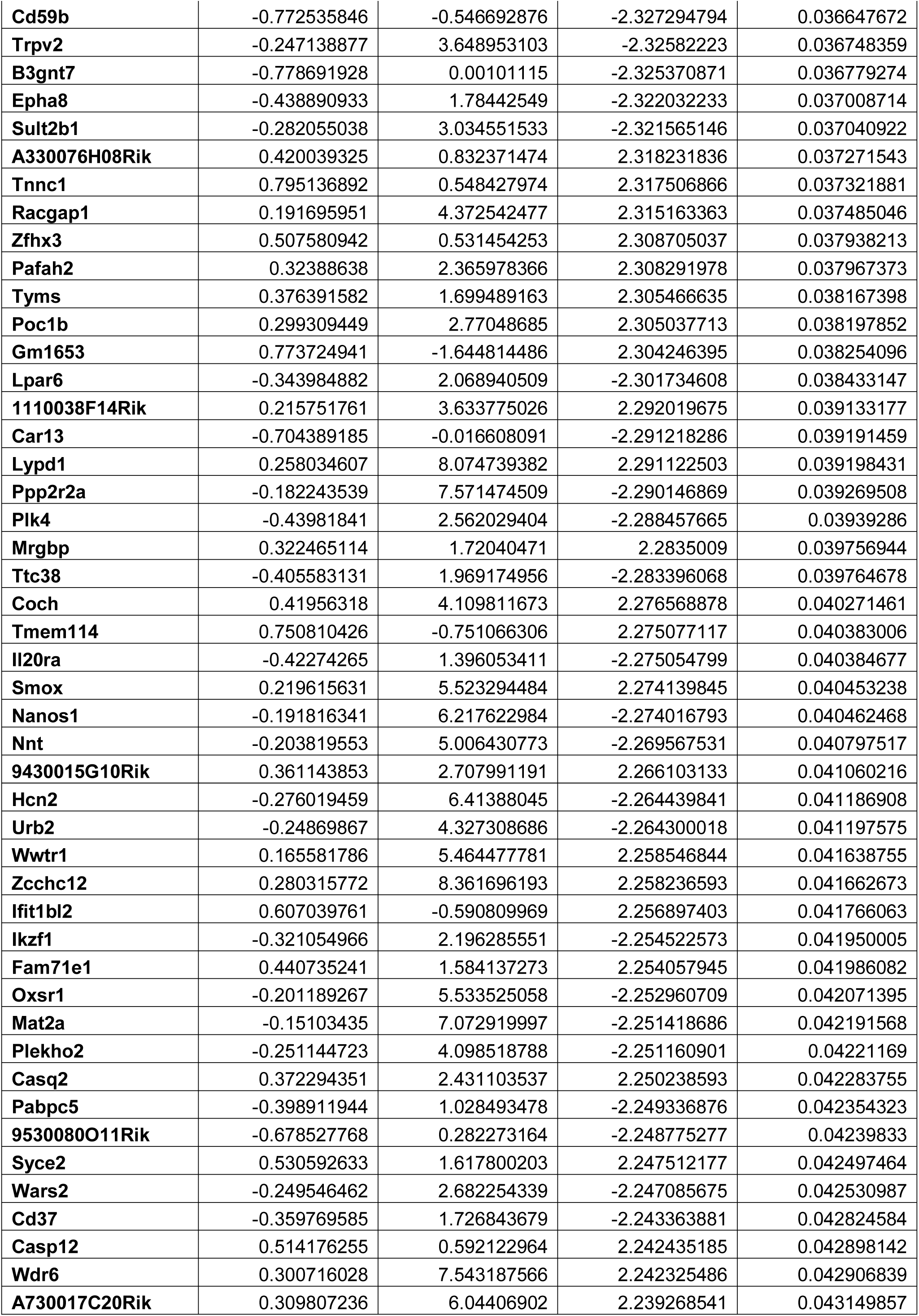
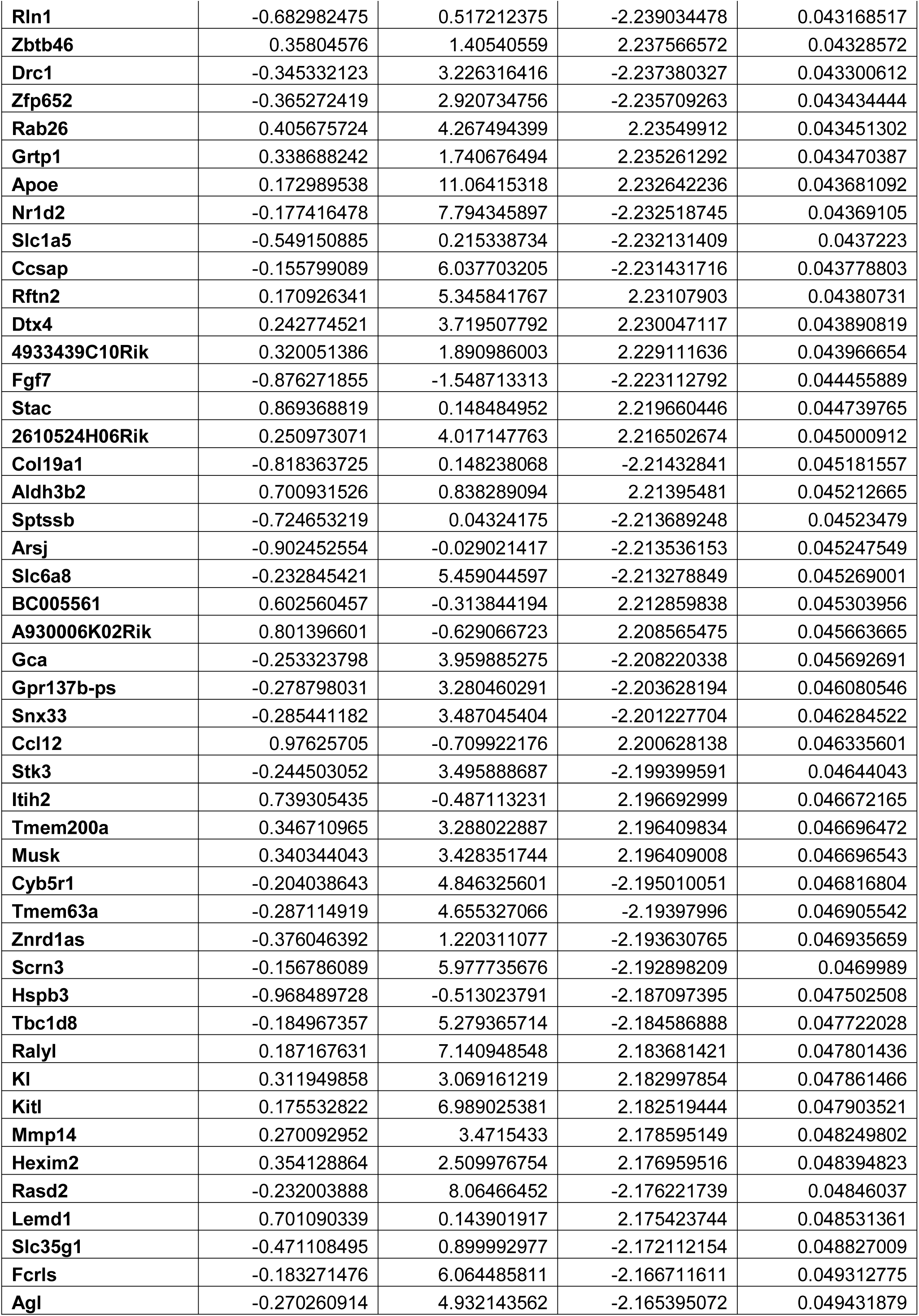
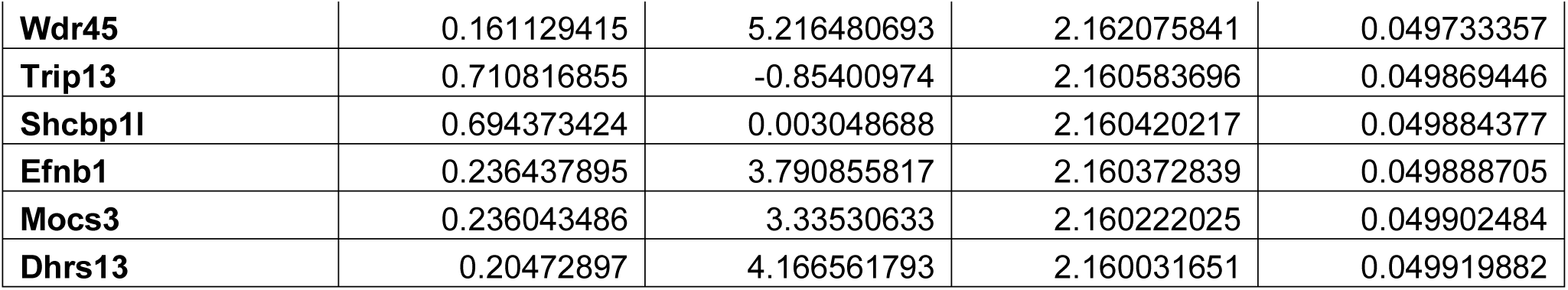
Differential expression analysis in the NAc for WT compared to Npas2 mutant females in the dark phase.

**Table 2.**
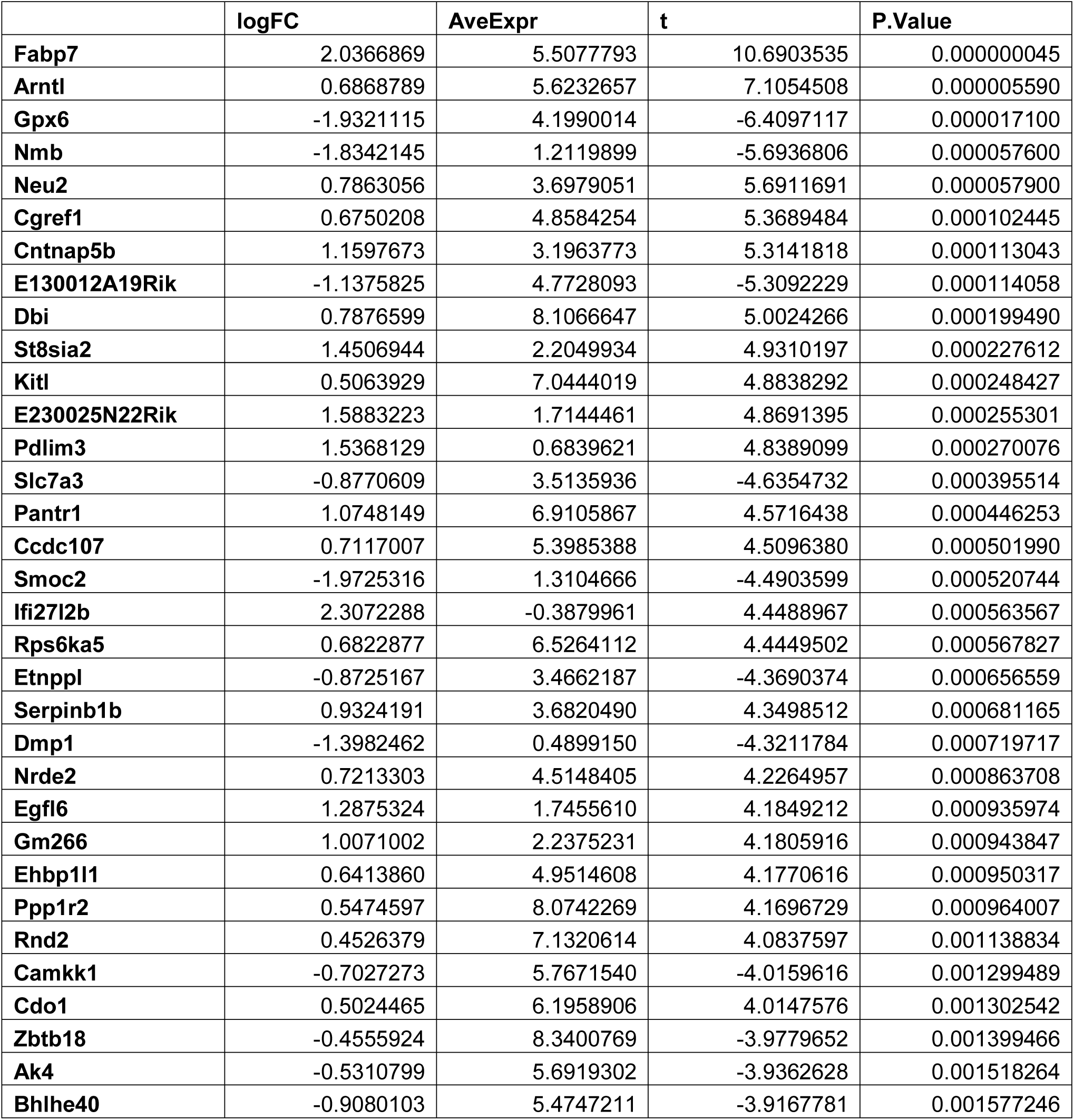
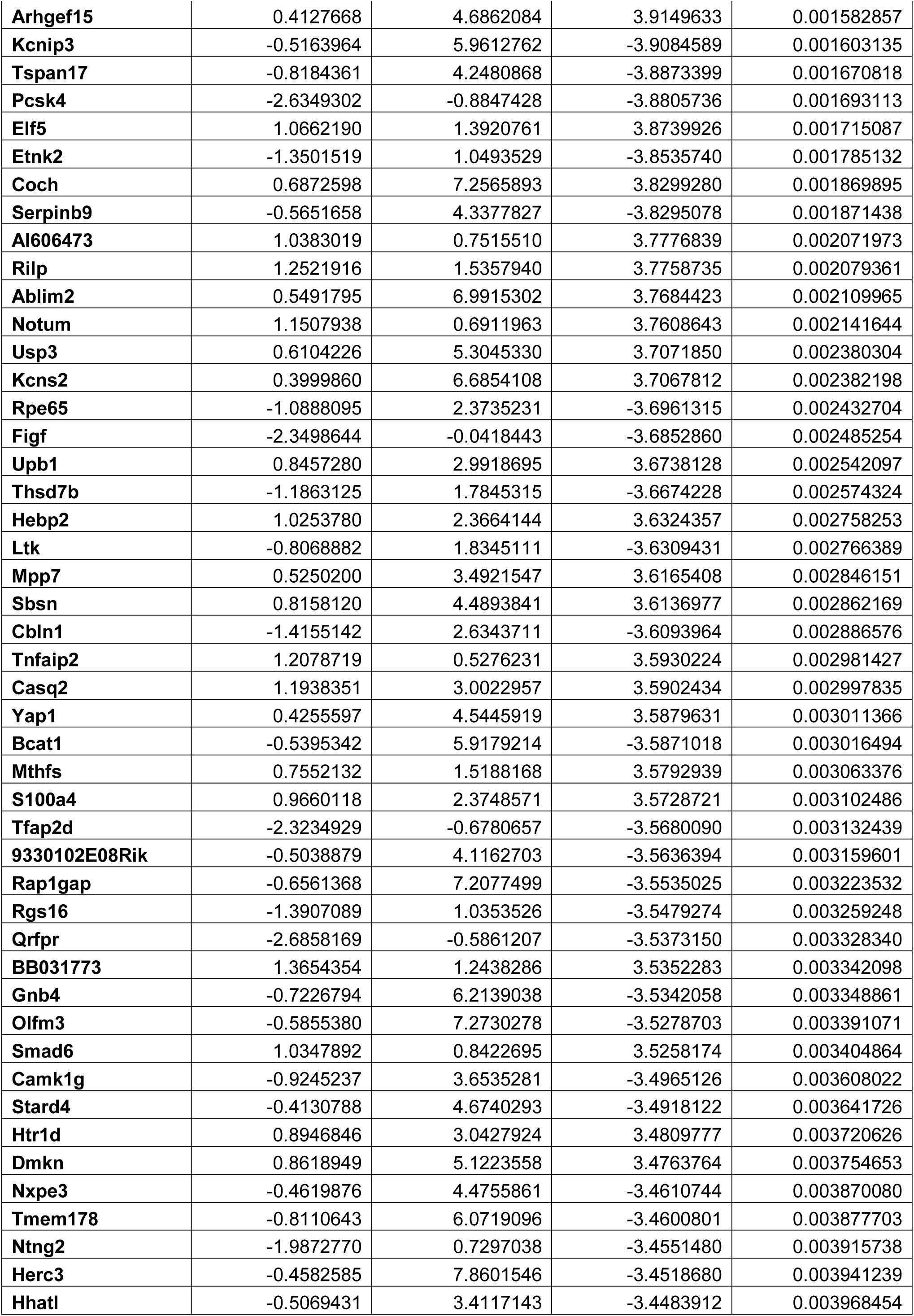
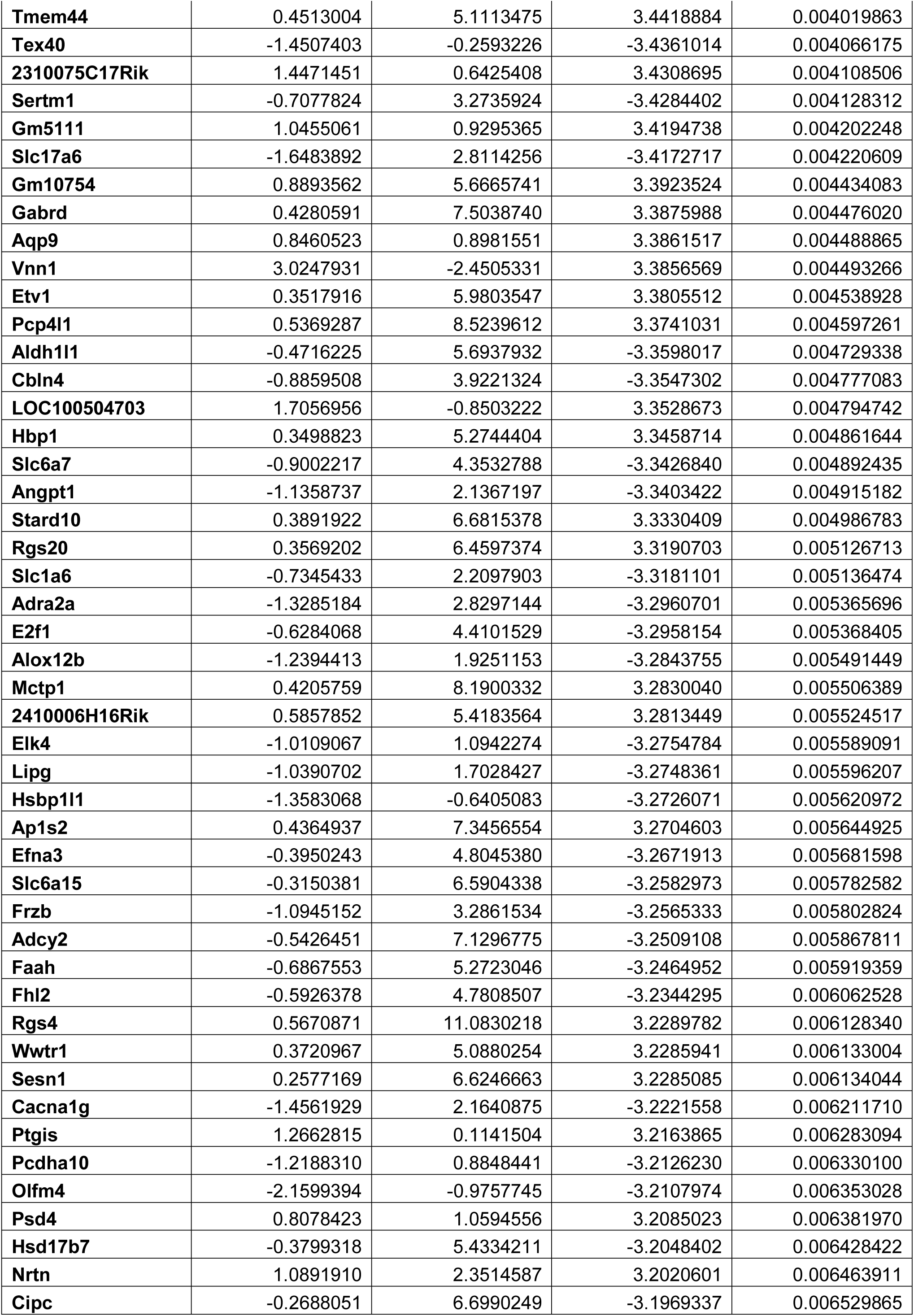
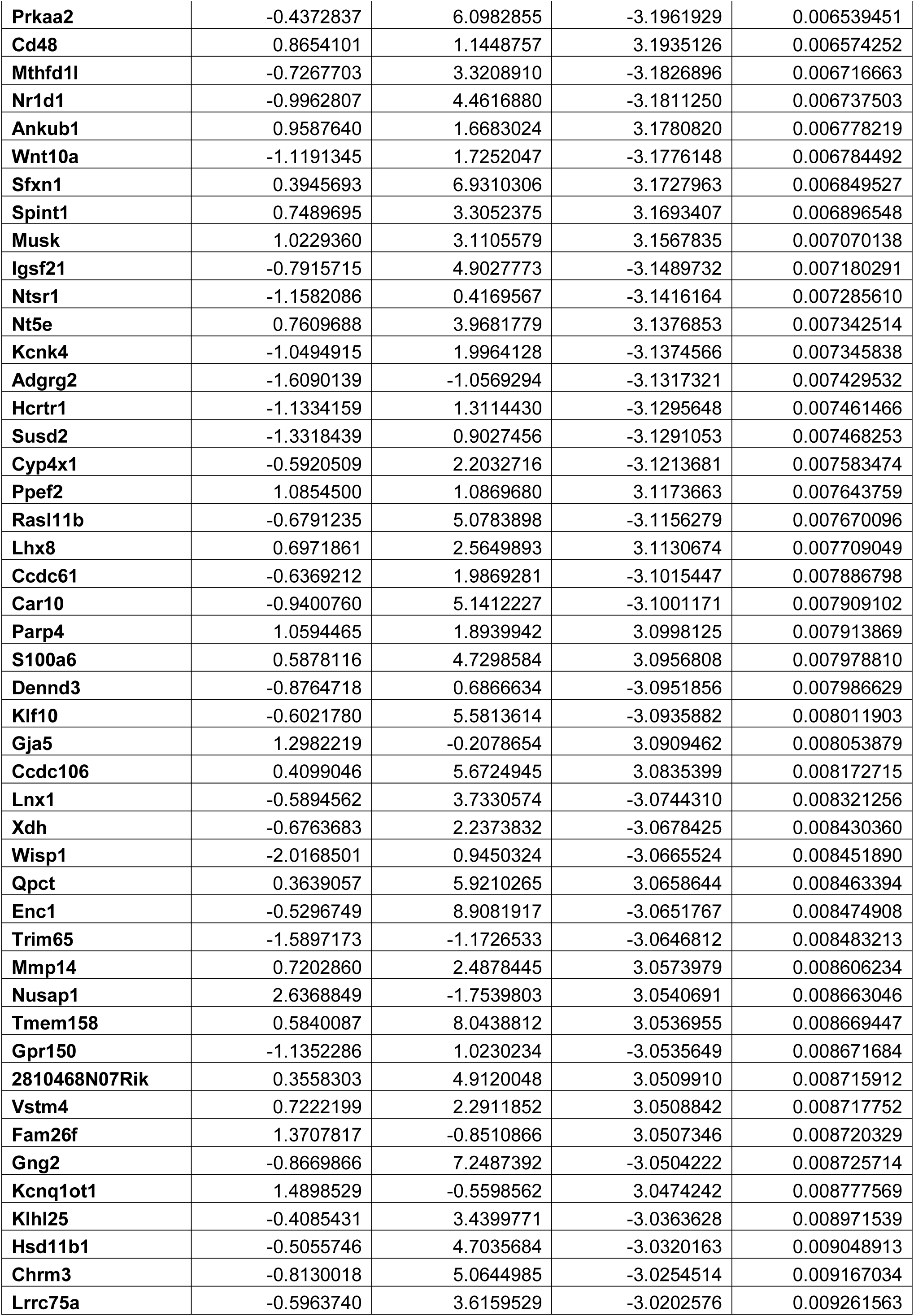
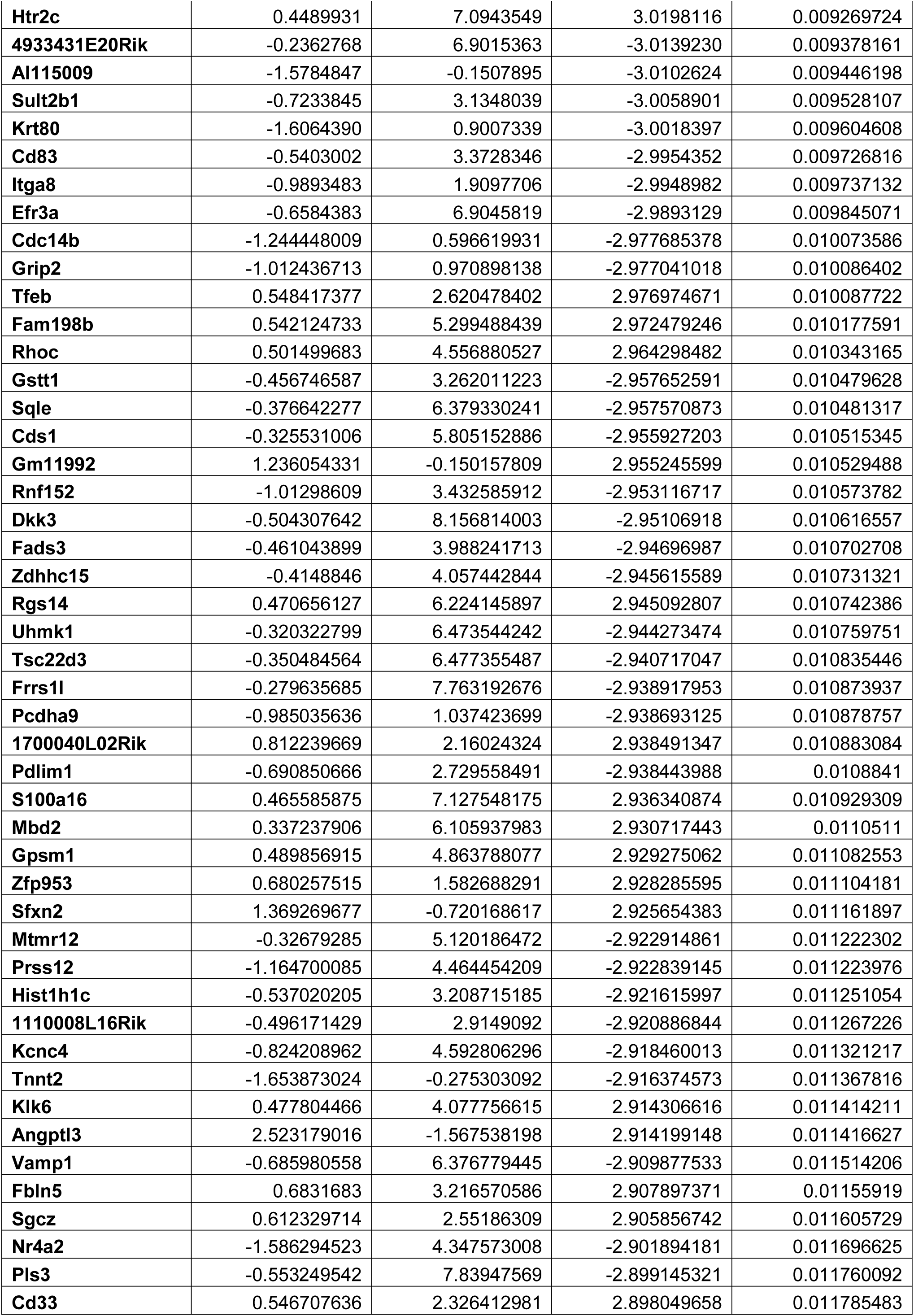
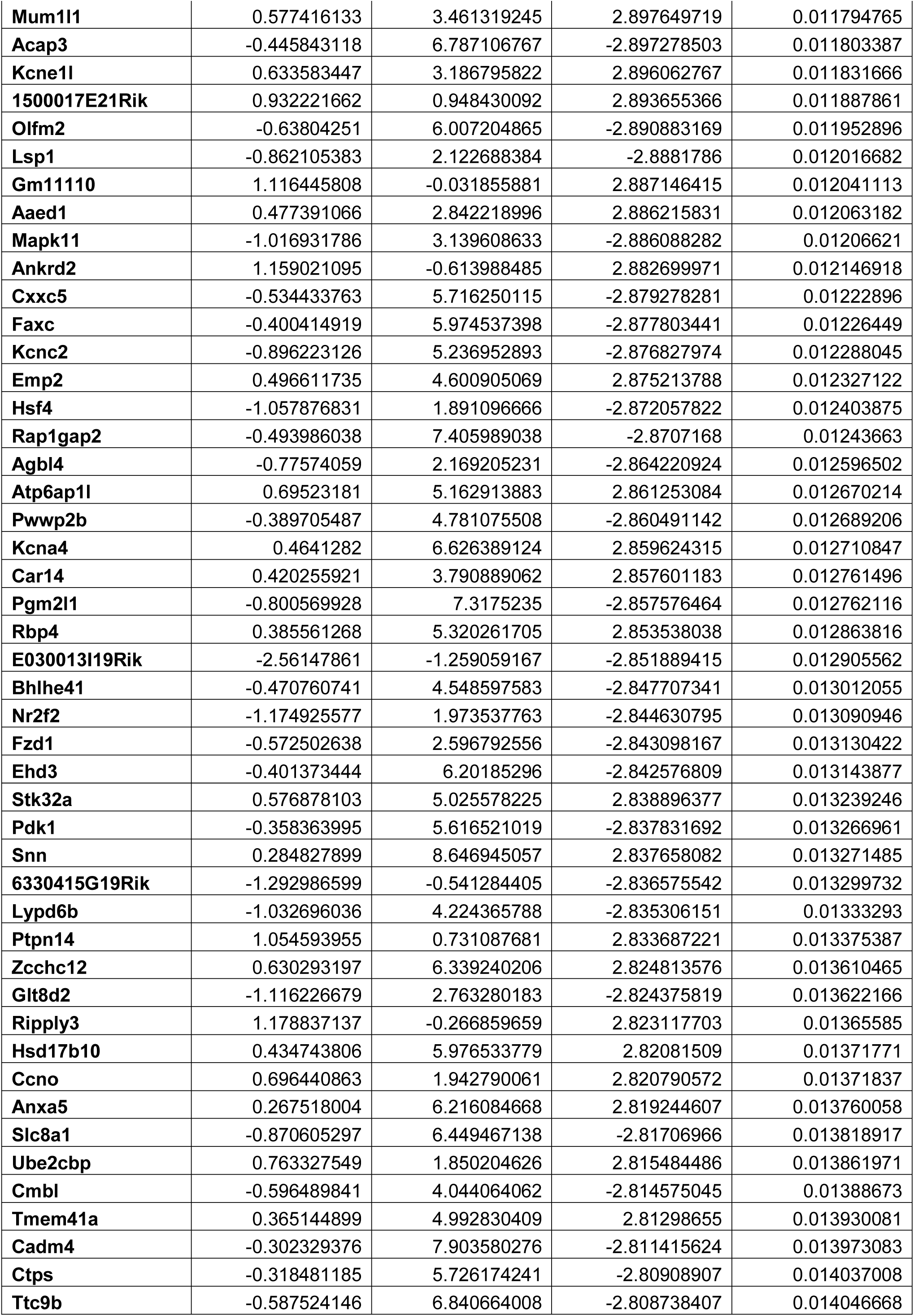
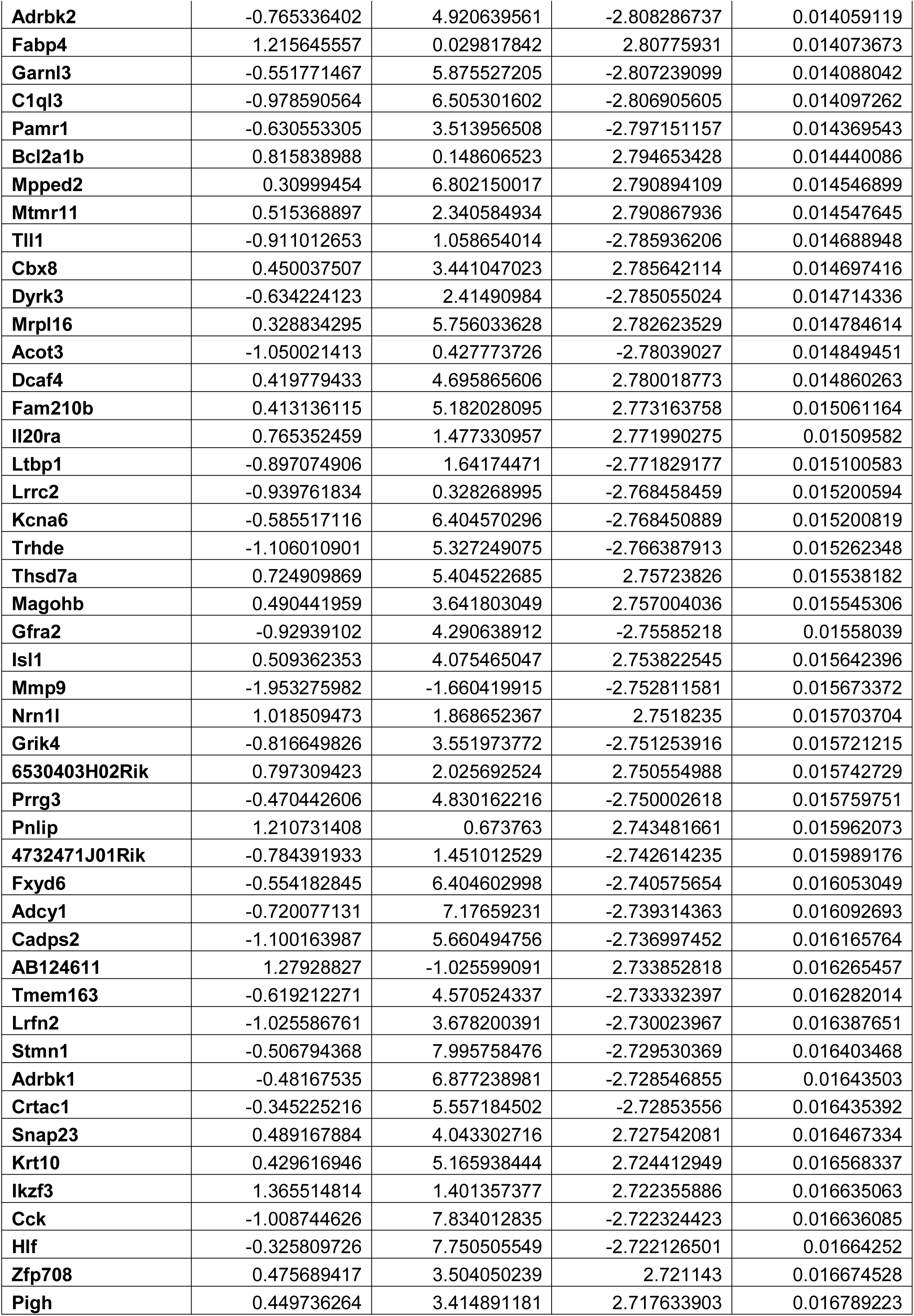
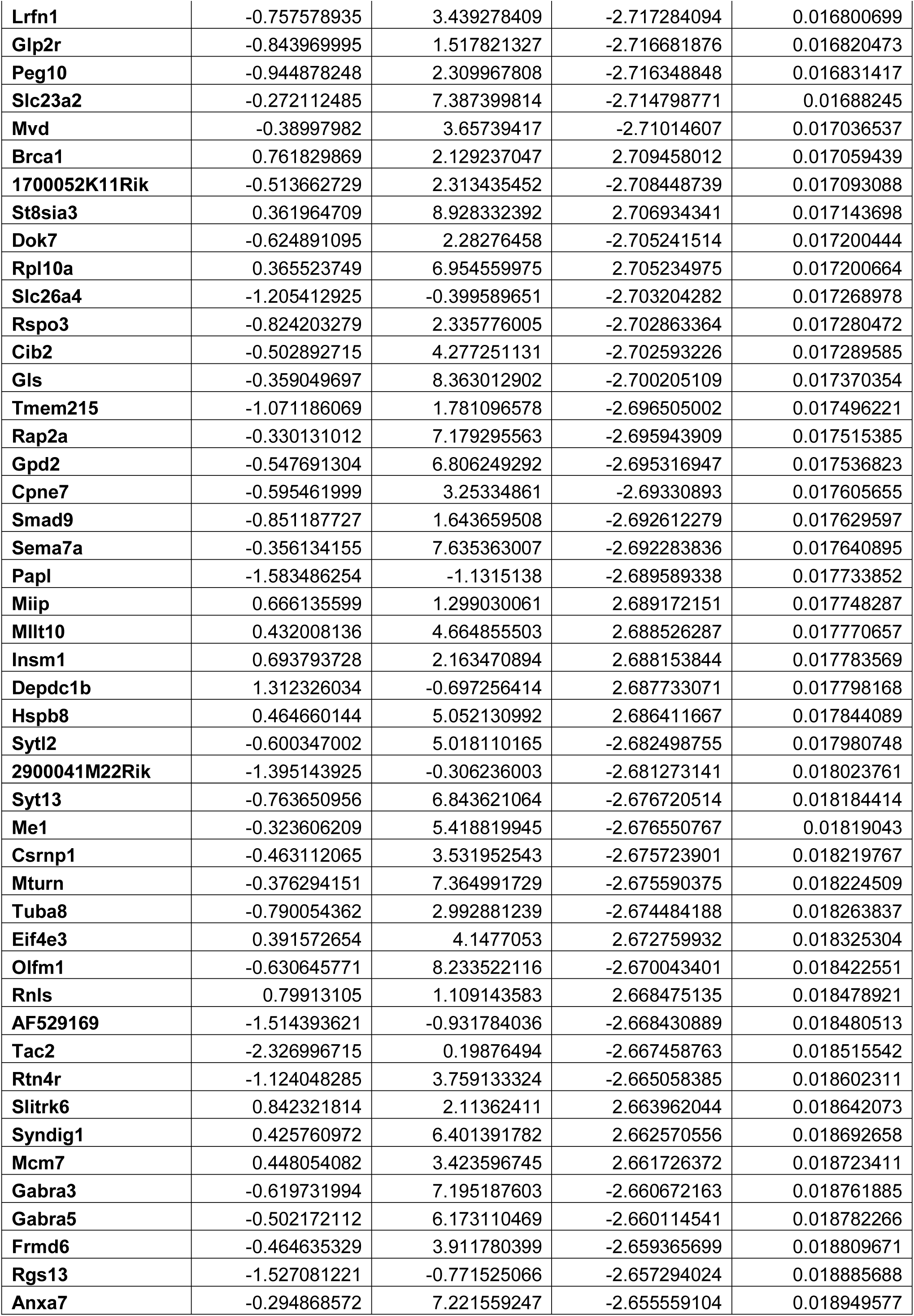
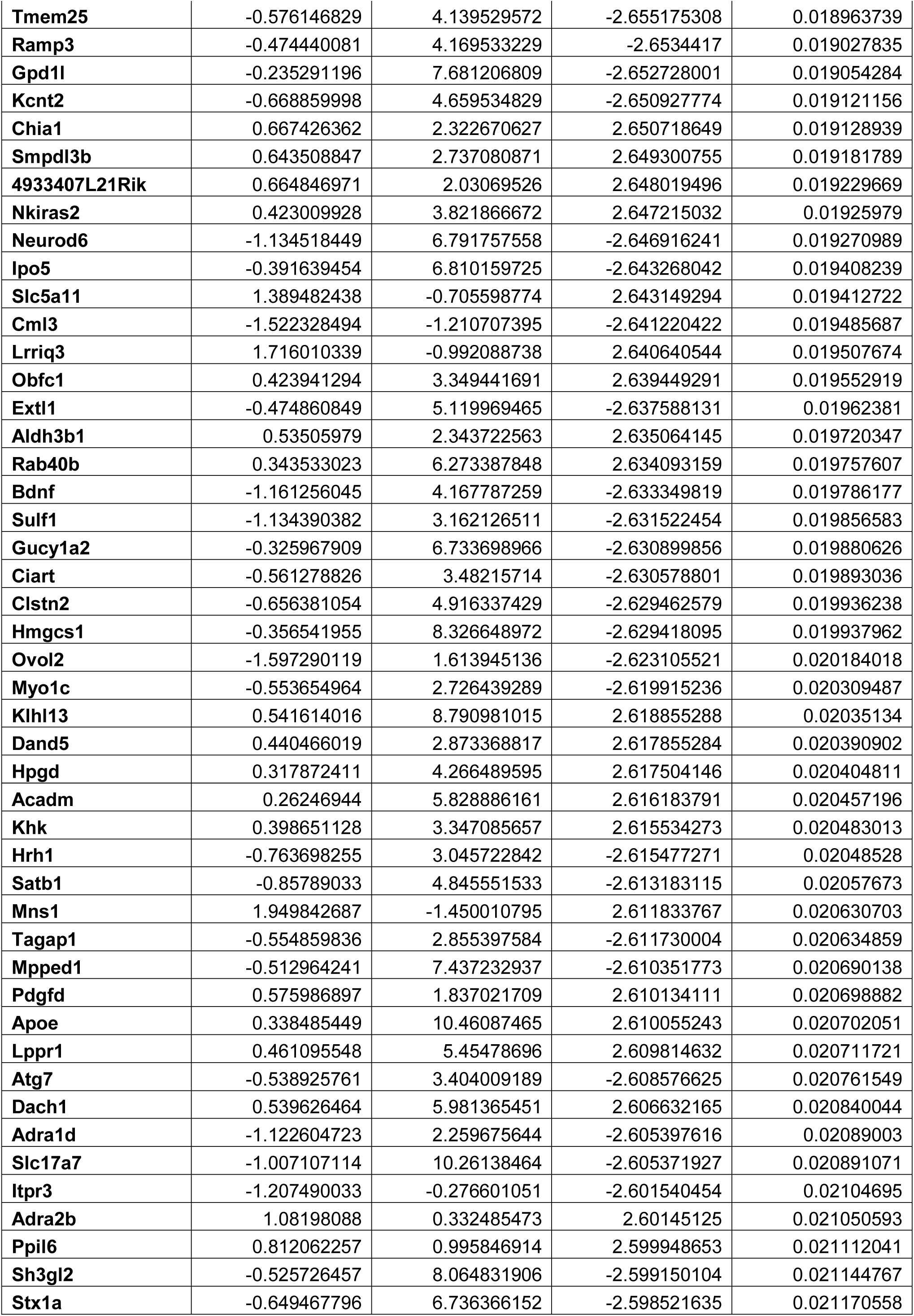
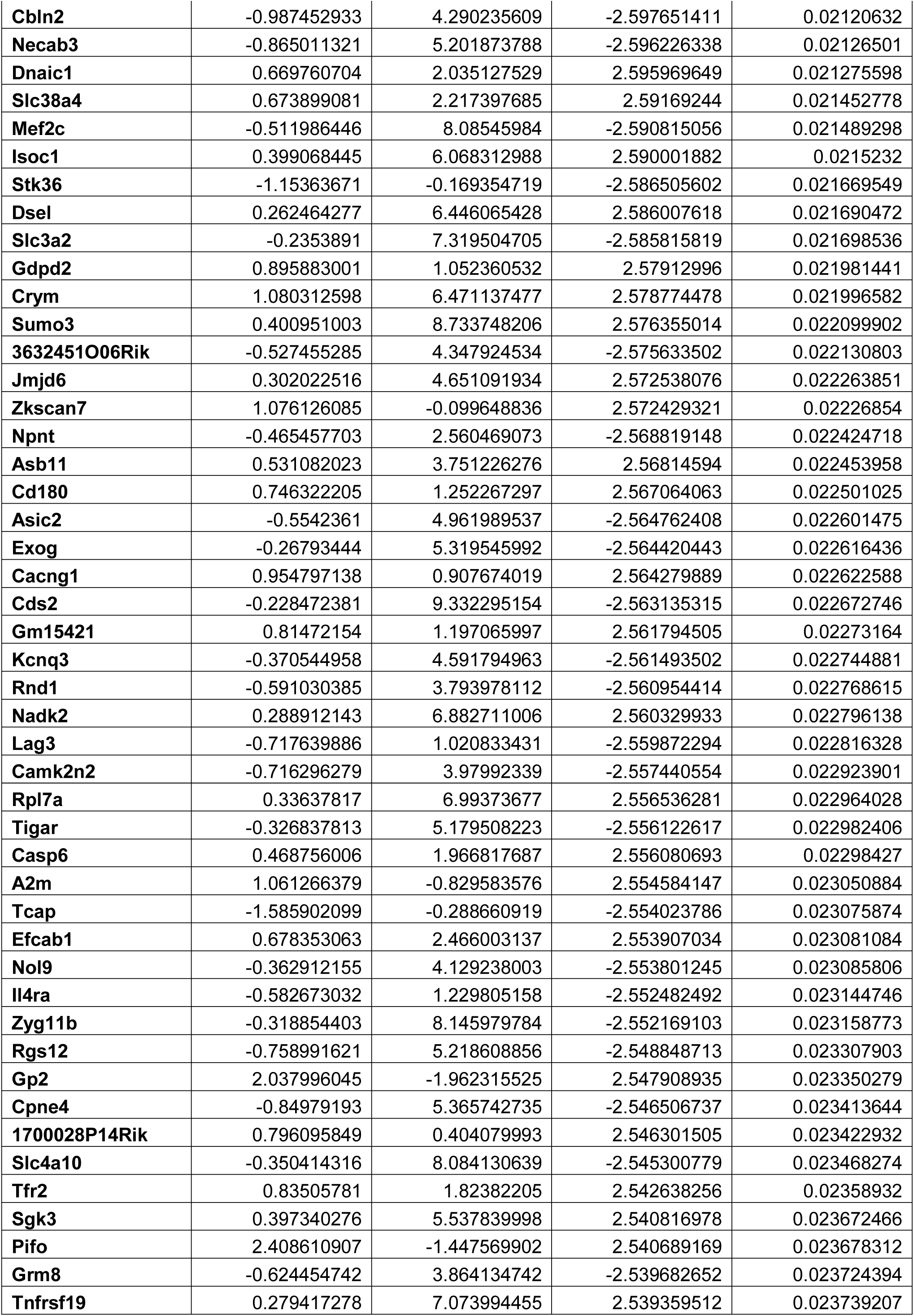
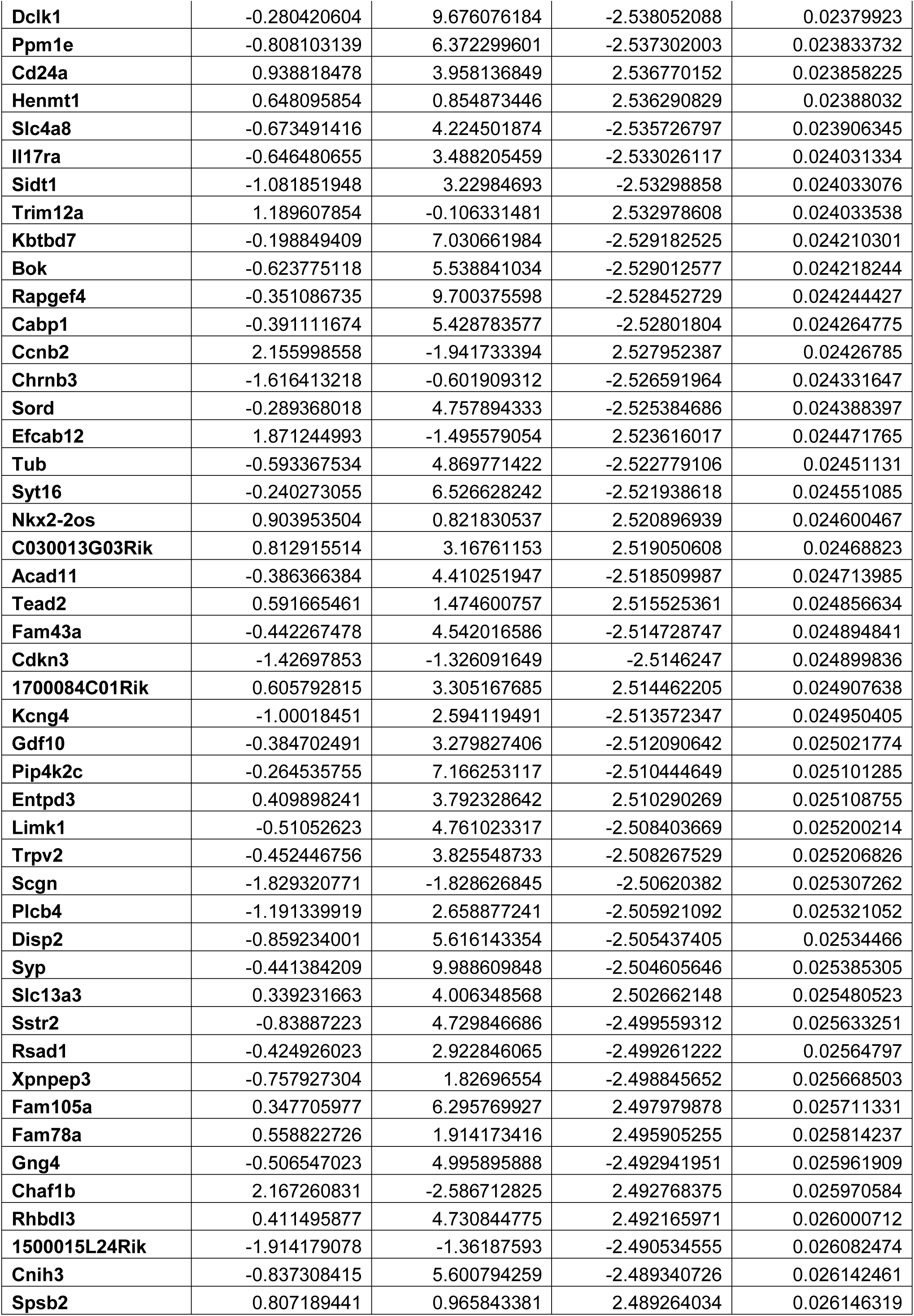
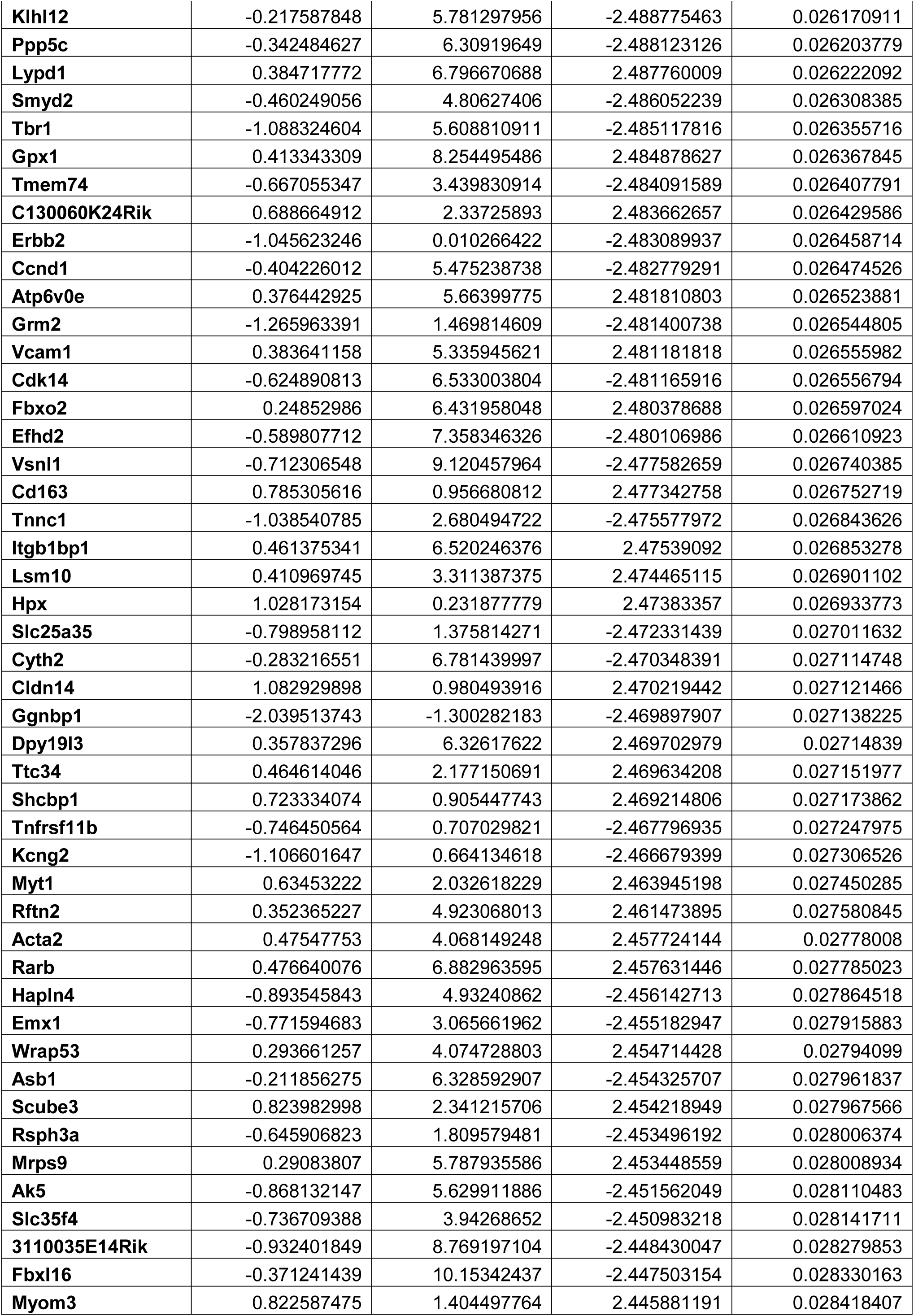
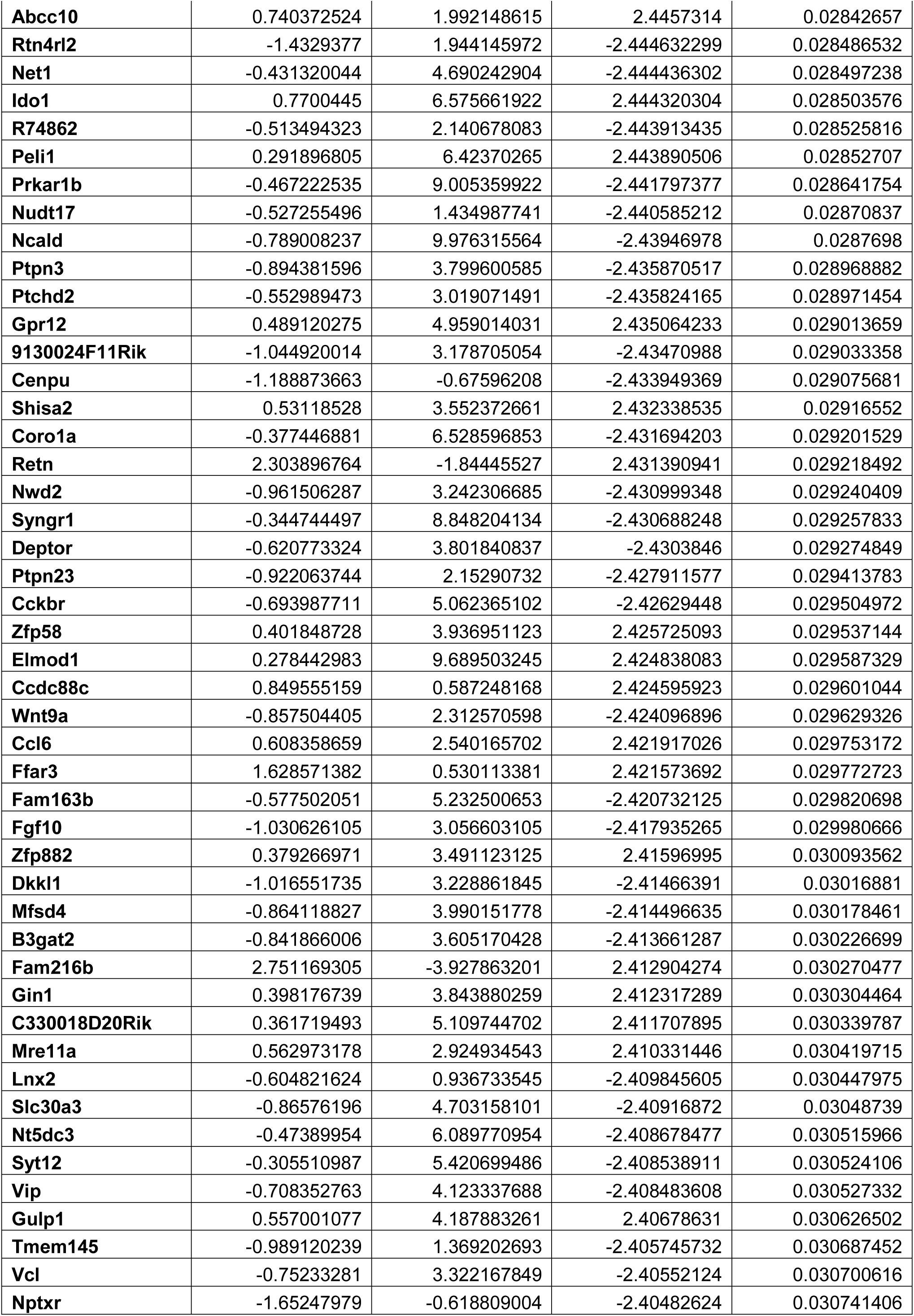
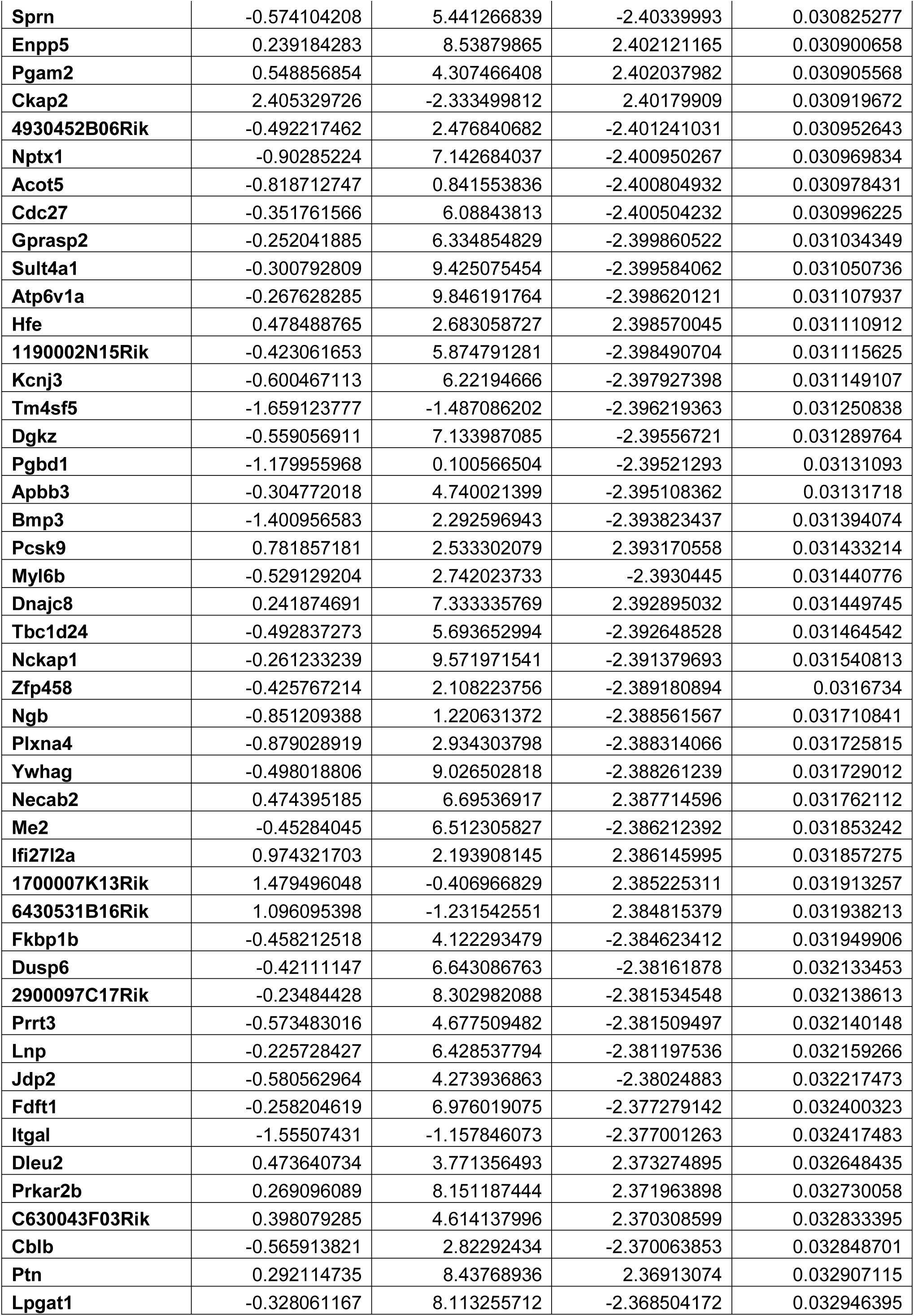
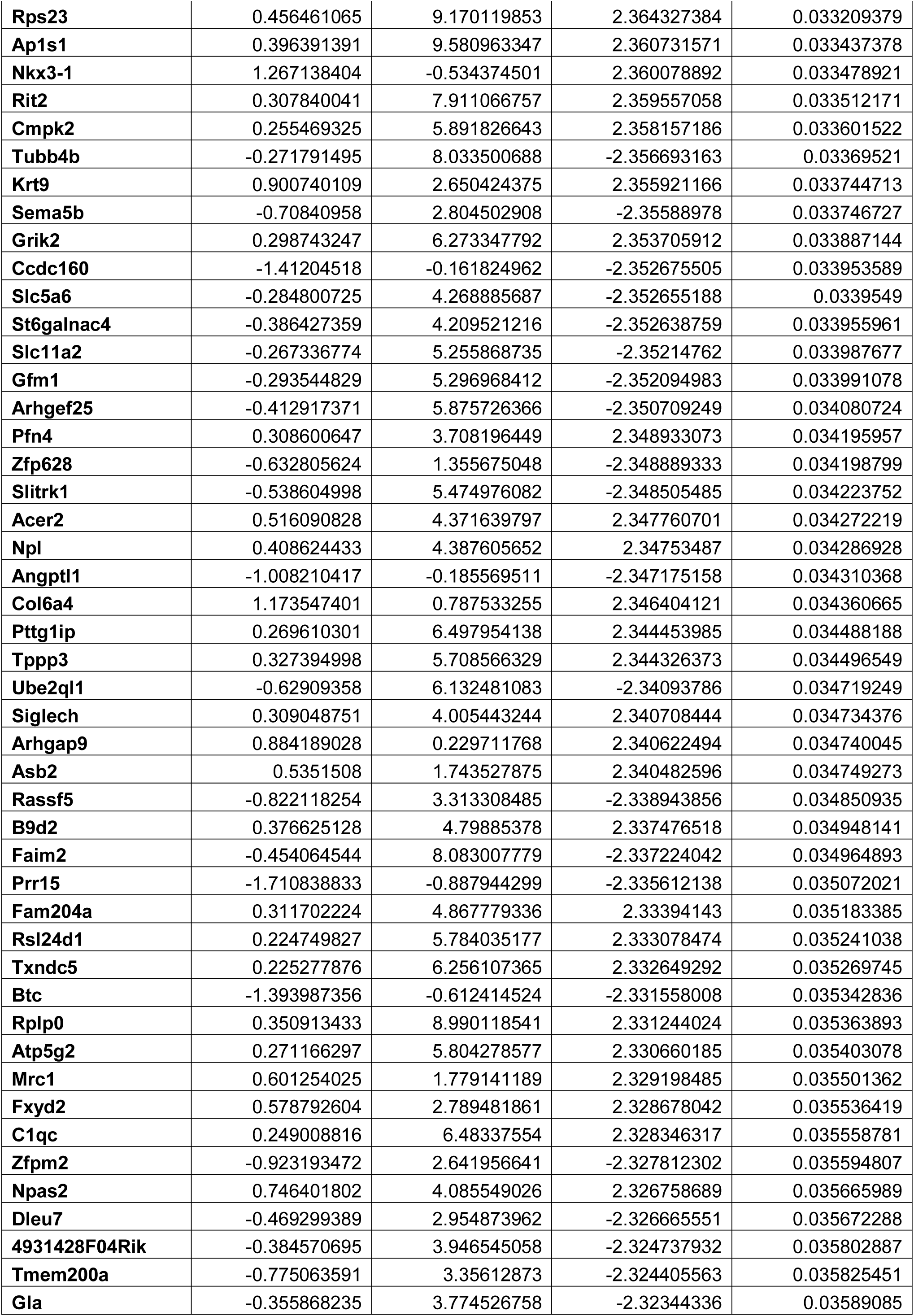
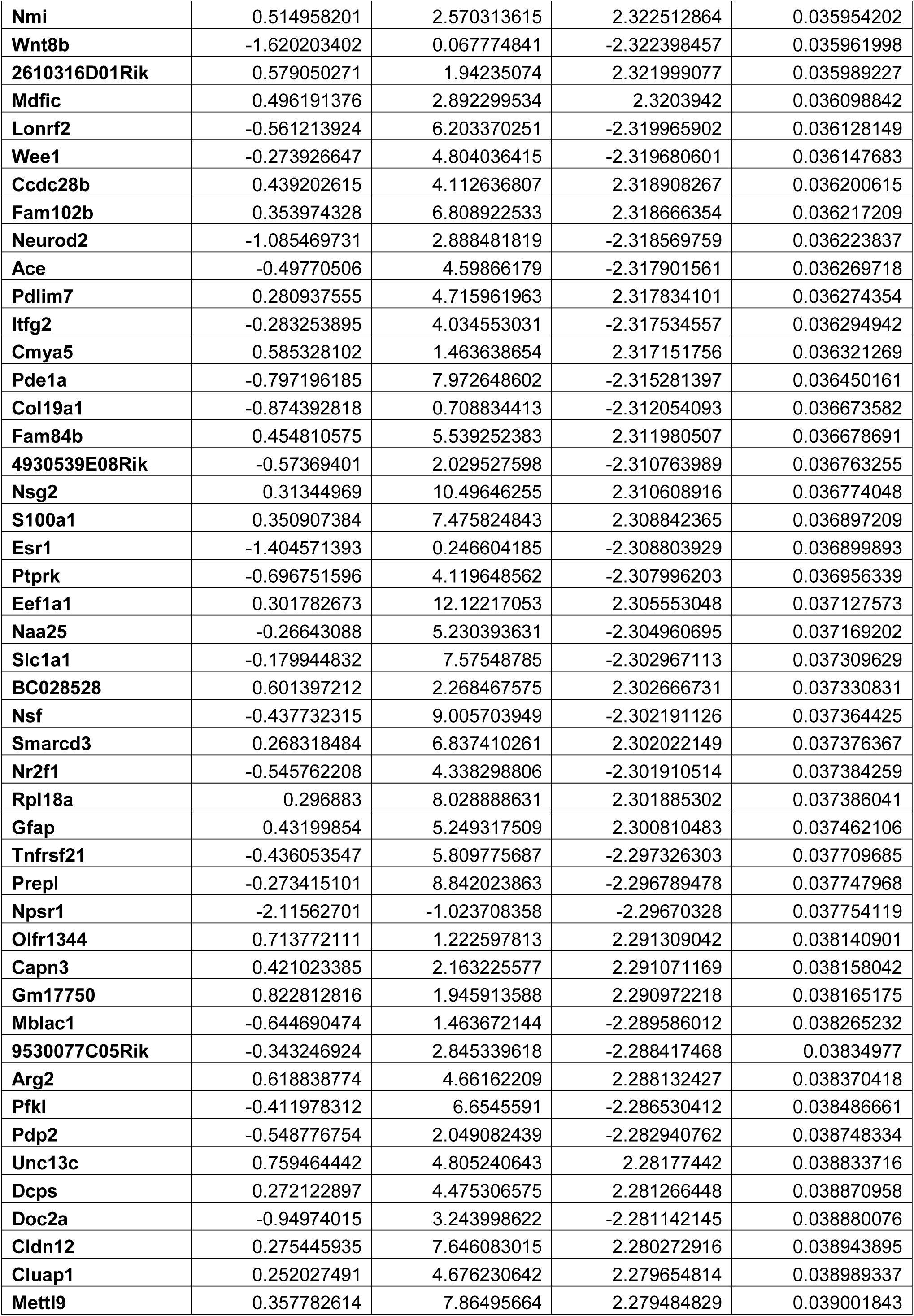
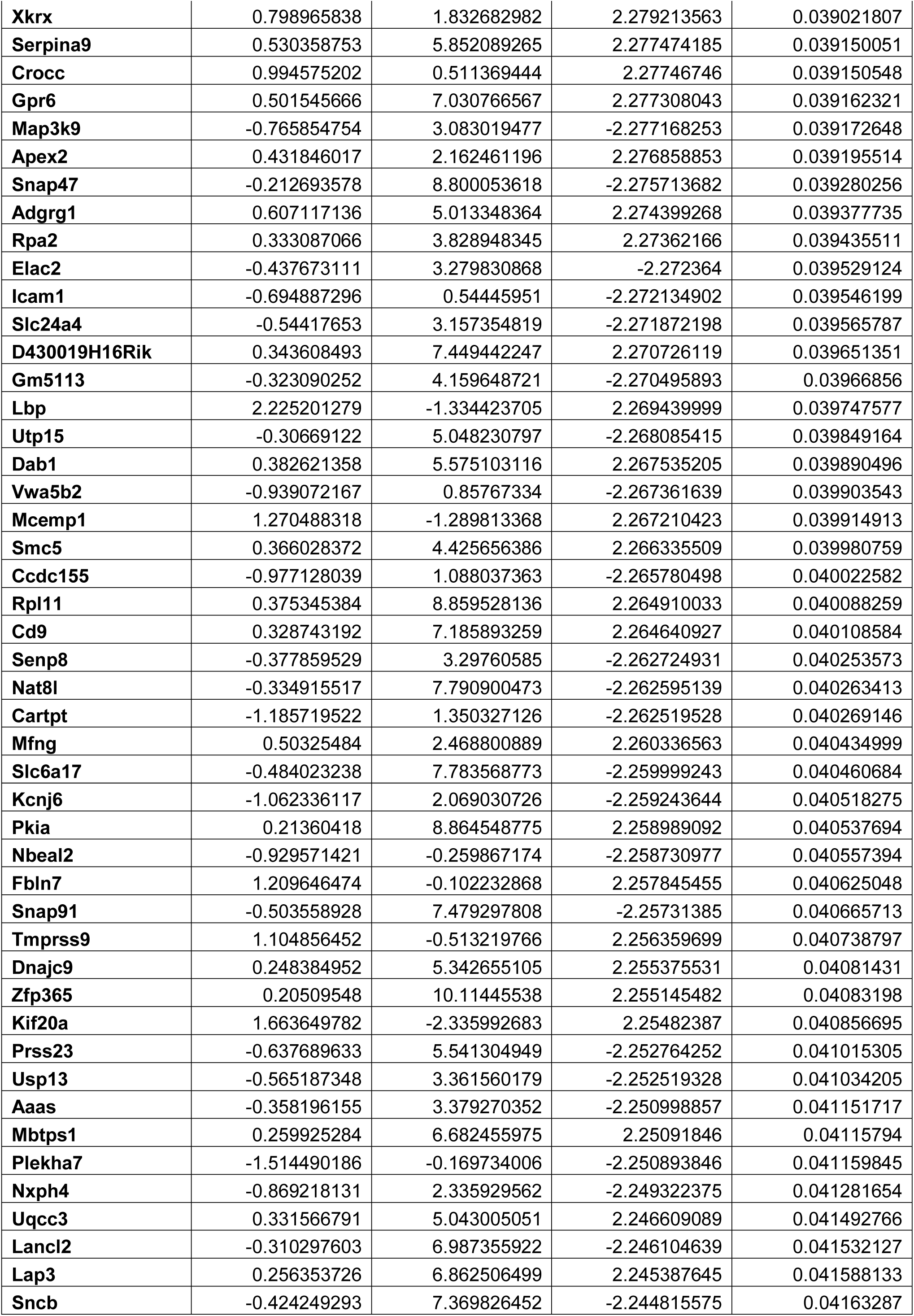
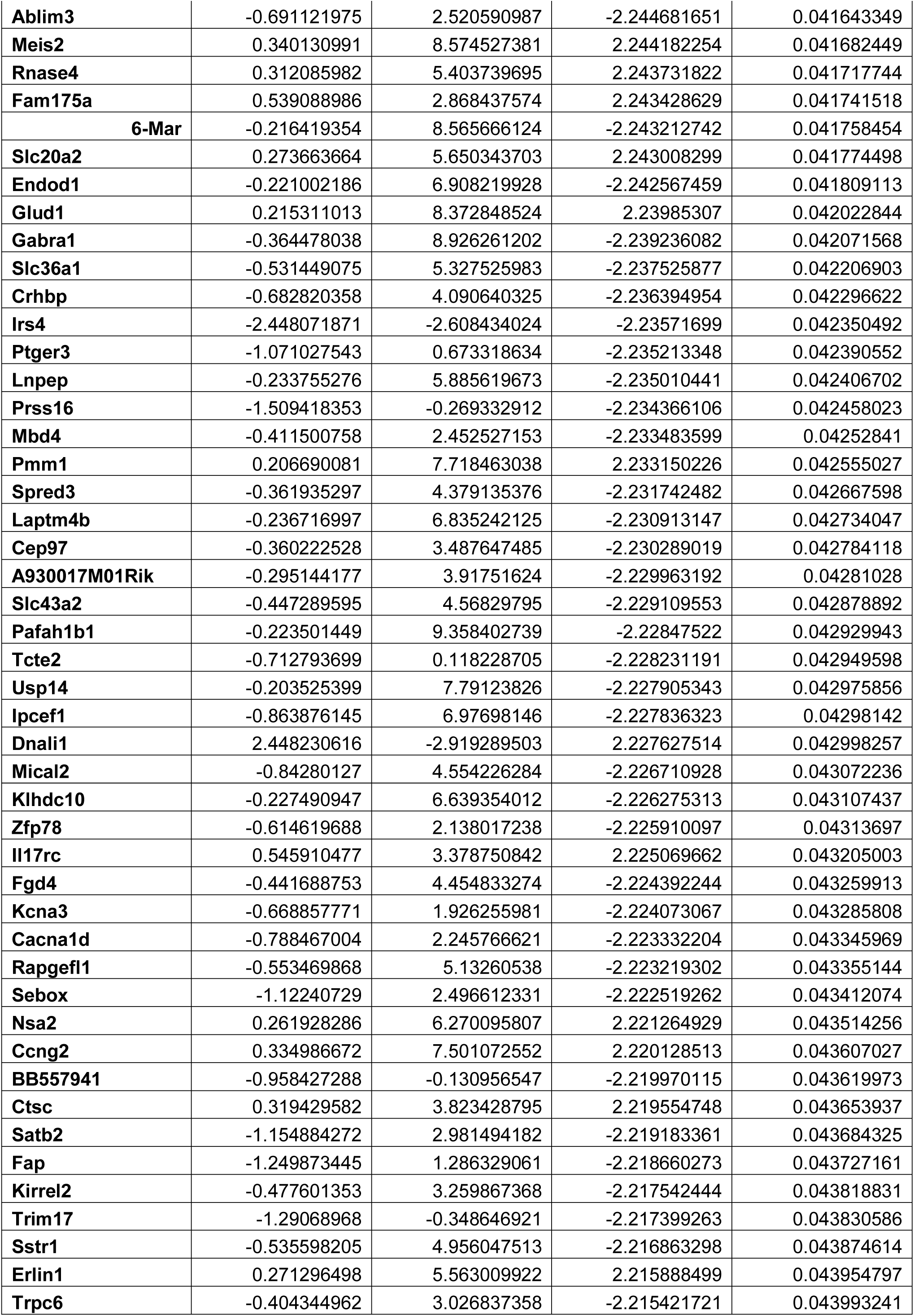
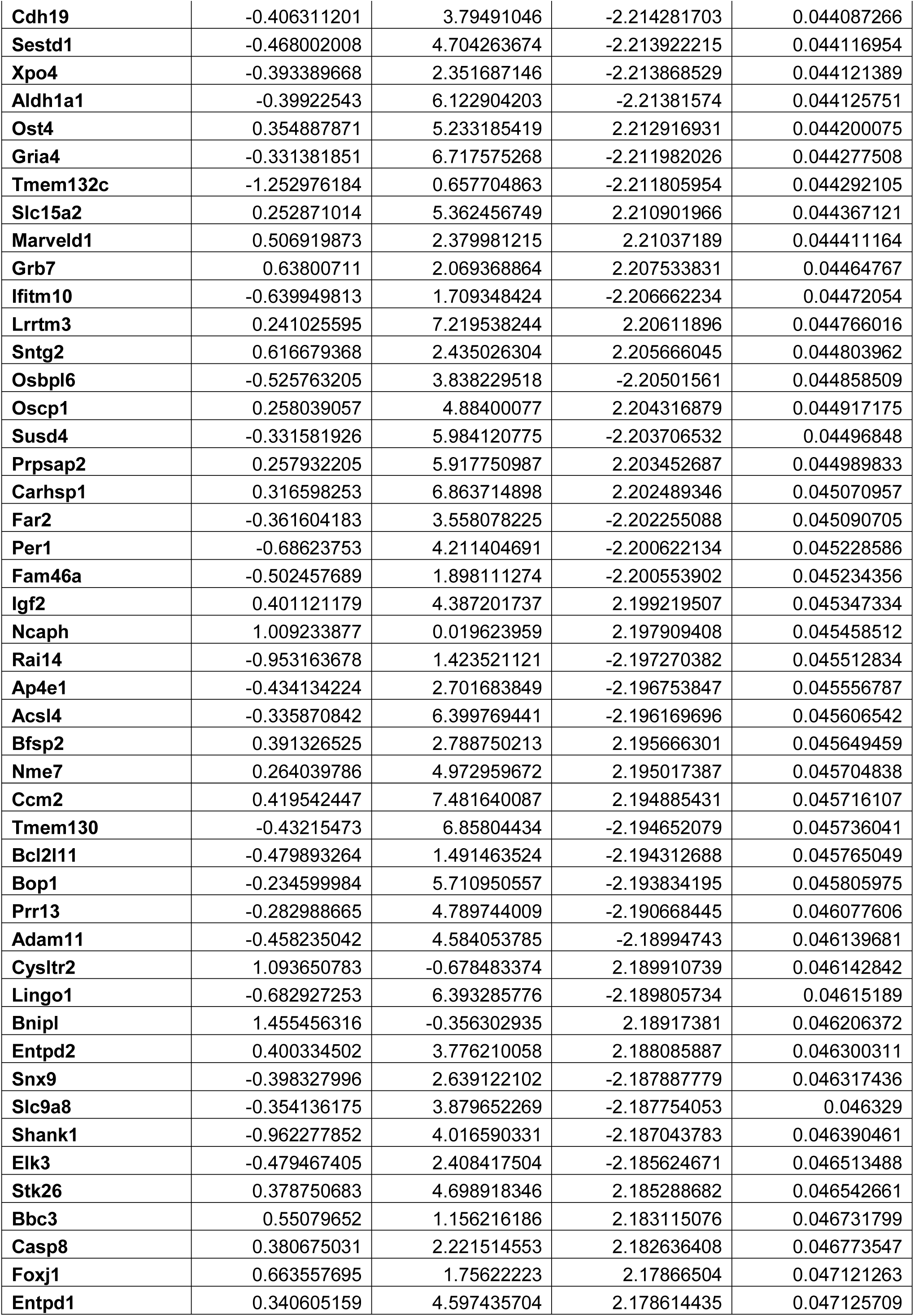
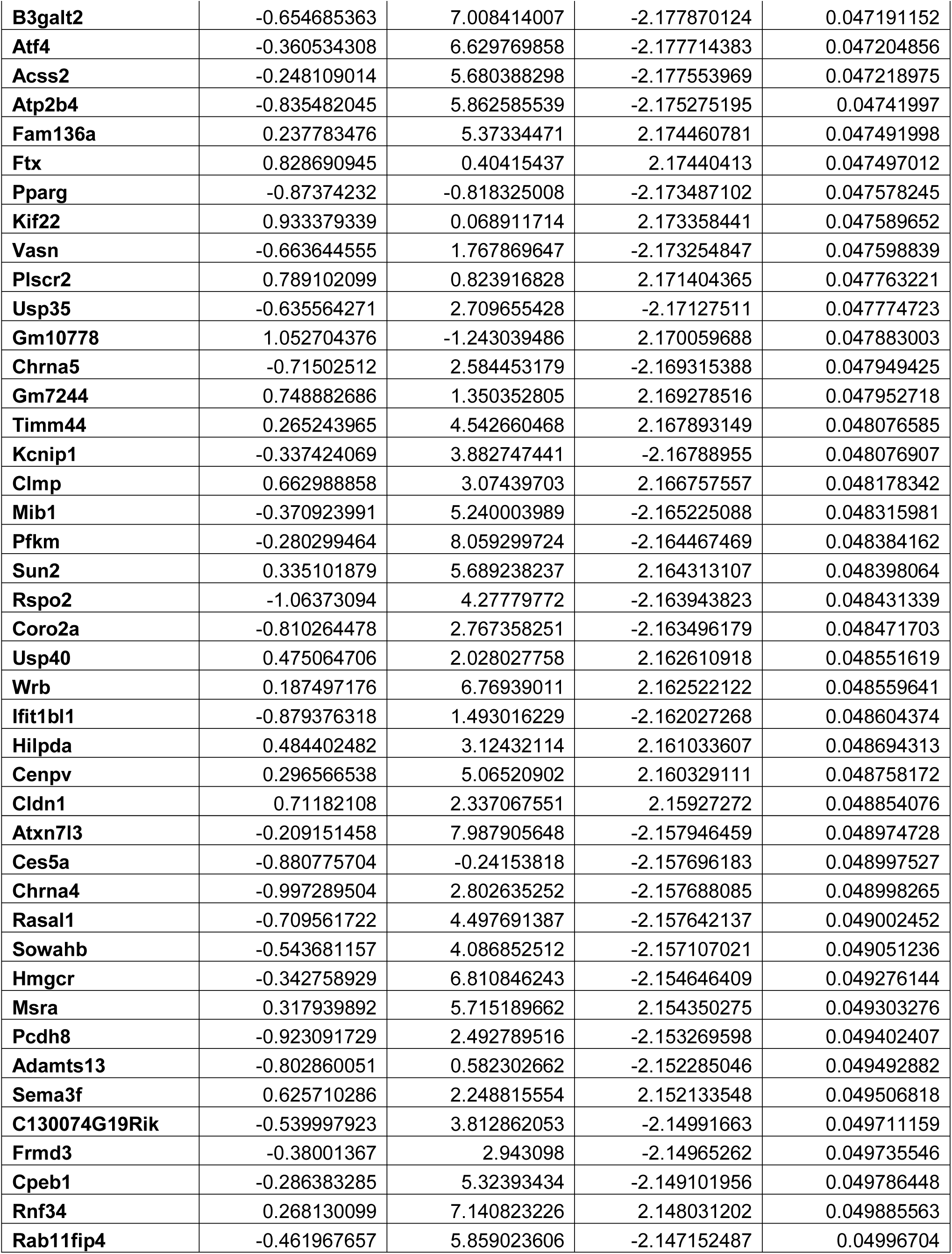
Differential expression analysis in the DLS for WT compared to Npas2 mutant females in the dark phase.

**Figure 9.**
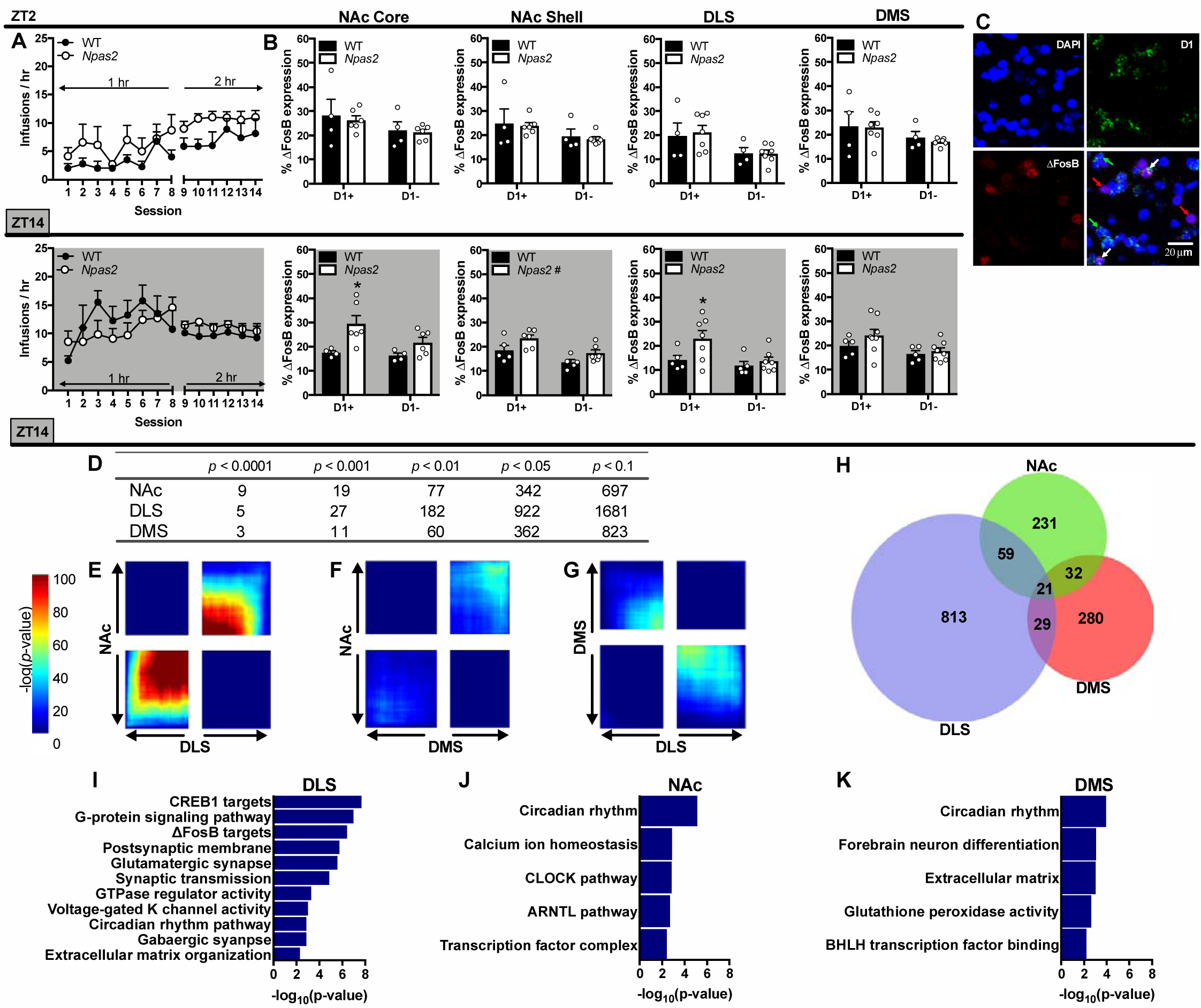
Striatal activation and differential gene expression in *Npas2* mutant females in the dark phase. (A) A cohort of female mice were trained to self-administer cocaine during either the light phase or dark phase. Infusions were limited to 25 in order to normalize cocaine intake between groups. (B) No differences were found between genotypes in.**Δ**FosB expression following light phase cocaine self-administration. However, following dark phase self-administration FosB expression was increased in *Npas2* mutant females in the NAc core, shell and DLS. This expression was specific to D1+ cells in the NAc core and DLS. Throughout, cocaine-induced.**Δ**FosB expression was greater in D1+ compared to D1-cells. (C) Representative staining is shown with the nucleus (DAPI, blue), D1 (green) and.**Δ**FosB (red) shown individually and as a composite image with 20 µm scale bar shown. (D) Table showing the cumulative number of DE genes between *Npas2* mutant and WT female mice in the dark phase with a fold change (FC) cut-off of 1.3. Various significance cutoffs are shown for the NAc, DLS and DMS. (E-G) RRHO plots showing various levels of overlap between genes that are changed with *Npas2* mutation in the NAc, DLS and DMS. (E) There is a high level of overlap between genes that are changed in the NAc and DLS, while (F) very few genes are changed in both the NAc and DMS of *Npas2* mutants. (G) Genes in the DMS and DLS are changed in opposite directions in *Npas2* mutant females. (H) Venn diagram showing unique or overlapping DEGs in the NAc, DLS and DMS with a cut-off of p<0.05 and FC=1.3. (I-K) Highly significant pathways altered in *Npas2* mutant females in the DLS, NAc and DMS are shown. Extended data list DEGs in the NAc (Figure 8-1), DLS (Figure 8-2) and DMS (Figure 8-3). Means+SEMs, ^#^p<0.1; *p<0.05, *n*=4-8.

We used a ranked p-value meta-analysis (Huo et al., 2020) to identify DEGs specific to the DLS and NAc. Of the 463 DEGs (meta-p<0.05), many that are similarly regulated appear to modulate neurotransmission, which might contribute to increased self-administration seen here. These DEGs encode potassium channels (*e*.*g. Kcn1l, Kcnc2, Kcna4, Kcna6*, etc), GABA receptor subunits (*e*.*g. Gabrd, Gabra3*, etc), matrix metallopeptidases (*Mmp14*), as well as the small rho GTPase *Rhoc* and *Snap23*, which encodes machinery necessary for vesicular fusion. Potassium channels were one of the most significant DEG categories and were primarily upregulated in both the DLS and NAc of *Npas2* mutant females.

Importantly, IPA revealed that highly significant DE pathways in *Npas2* mutant females in the DLS include CREB1 targets, G-protein signaling and ΔFosB targets (Fig.9i). This suggests that even at baseline, *Npas2* mutant females might have increased ΔFosB and CREB-mediated transcription in the DLS, which is further exacerbated in D1+ cells by cocaine self-administration (Fig.9b), but future experiments are needed to test this hypothesis directly.

As expected, the DEGs found across all 3 regions included mainly core circadian genes: *Ciart, Arntl* (*Bmal1*), *Nr1d1, Cipc, Bhlhe41* and *Dbp. Arntl* was the only upregulated circadian gene in *Npas2* mutant mice, likely to compensate for its inability to form a complex with the mutated *Npas2*. IPA also found significant enrichment in the pathway “circadian rhythms” across all regions, but especially the NAc (Fig.9ijk). Fifteen other DEGs were identified in all 3 regions suggesting they might be important target genes of *Npas2*. These include protein kinases (*Kitl, Camkk1*) and genes that are protective against oxidative stress (*Gpx6* and *Hebp2*). *Fabp7* was the most upregulated gene in *Npas2* mutants and plays a role in fatty acid uptake, transport and metabolism in astrocytes (Chmurzyńska, 2006).

## Discussion

Overall, *Npas2* mutation increases cocaine intake and the propensity to self-administer cocaine in a sex- and circadian-dependent manner. In the light phase, cocaine taking, reinforcement and motivation are increased across sex. During the dark phase, females are more affected; reinforcement and motivation are increased across sex, while females mutants have increased drug taking, extinction responding and reinstatement. Females also appear to be driving effects on motivation, since female mutants have the highest breakpoint ratios during both TOD, although a sex difference was not found. Interestingly, while males show contradicting effects of *Npas2* mutation on light phase cocaine reward and self-administration, self-administration is uniquely affected in females. While it isn’t uncommon to find opposing results for these drug-related behaviors (Larson et al., 2011), detecting a selective change in volitional drug self-administration, as with females here, is more typical.

It is well known that sex differences exist in circadian rhythms and that circulating estradiol contributes to these differences (Varnäs et al., 2003; Krizo & Mintz, 2015). Sex differences are also frequently found in the rhythmicity of peripheral circadian genes (Lu et al., 2013) and the master pacemaker, the suprachiasmatic nucleus (SCN)(Bailey & Silver, 2014). Although less is known about the extra-SCN brain, females show earlier peaks in circadian genes in the prefrontal cortex in humans (Lim et al., 2013) and the robustness of diurnal gene rhythms can vary by sex and brain region in rodents (Chun et al., 2015). These findings indicate that SCN-independent behaviors could also be affected by circulating hormones.

Despite an intersection between rhythms, hormones and extra-SCN behavior, research investigating how altered circadian rhythms, for example from circadian gene mutations, affect behavior sex-dependently is limited. One key paper found that female *Clock* mutant mice show more robust increases in exploratory and escape-seeking behavior (Easton et al., 2003). Sex differences have been examined in *Npas2* mutants, but only in the context of sleep and locomotor activity. Importantly, *Npas2* mutation doesn’t affect rhythms differently in males and females at baseline (Dudley et al., 2003), only under pathological conditions, such as sleep deprivation (Franken et al., 2006), suggesting our observed sex differences are not confounded by baseline differences. Since cocaine exposure disrupts circadian rhythms and sleep (Schierenbeck et al., 2008; Angarita et al., 2016), self-administration could be inducing sex-dependent sleep and rhythm disruptions, which might lead to sex differences in cocaine intake.

To determine whether circulating hormones might contribute to increased self-administration in *Npas2* mutants, we ovariectomized female mice before dark phase cocaine self-administration. Female OVX mice with non-functional NPAS2 showed no increase in self-administration, suggesting ovarian hormones could be driving enhanced effects in *Npas2* mutant females. It is critical to continue examining how circadian genes affect behavior sex-dependently, particularly in the context of SUD since its prevalence varies by sex (Kosten et al., 1993; Robbins et al., 1999; Kennedy et al., 2013). Interestingly, our sham controls only showed a trending increase (p=0.58) in self-administration, which is likely due to these mice being older or the stress of an additional surgery. These factors are crucial to consider when investigating sex differences in future studies.

In order to further identify how mutations in *Npas2* contribute to SU in females, we measured striatal activation using ΔFosB. While other Fos proteins are transiently induced by acute drug exposure, ΔFosB is a stable, long-lasting variant induced by long-term exposure (Robison et al., 2013). This allows activation from chronic drug self-administration to be measured. Here, female WT and *Npas2* mutant mice self-administered cocaine, but drug intake was limited in order to normalize increased cocaine exposure in mutant mice.

The striatum is primarily comprised of neuron populations expressing D1 or D2 dopamine receptors (Lu et al., 1998). Activation of D1-expressing neurons in the NAc promotes cocaine preference (Lobo et al., 2010) and cocaine activates D1 and D2 neurons differently (Bertran-Gonzalez et al., 2008), inducing ΔFosB in D1 neurons (Lobo et al., 2013). NPAS2 is highly enriched in the striatum (Garcia et al., 2000a), specifically in D1 neurons (Ozburn et al., 2015a) and we recently demonstrated that *Npas2* knockdown in the NAc increases glutamatergic excitability onto D1 cells (Parekh et al., 2019). Therefore, we measured ΔFosB expression in D1+ and D1-cells throughout the striatum and found that ΔFosB expression was increased in D1+ neurons in the NAc core and DLS of *Npas2* mutant females. The NAc shell was also moderately affected, in D1+ and D1-cells, but effects in the dorsal striatum were confined to the DLS. Since the DLS mediates habitual drug seeking, and the DMS goal-directed decision making (Yin et al., 2004), *Npas2* mutant females may rely more heavily upon habitual decision-making strategies during self-administration. Future studies could use action-outcome contingency degradation or outcome devaluation to measure response strategies in *Npas2* mutant females.

Increased induction of ΔFosB in *Npas2* mutants was also specific to the dark phase when the behavioral effects of *Npas2* mutation are greatest. *Npas2* expression peaks in the NAc around ZT16 (Falcon et al., 2013) when dark phase self-administration was conducted. It stands to reason that mutating *Npas2* would have a greater impact on behavior when mRNA expression is elevated. Higher expression suggests a functional necessity for NPAS2, however, a parallel protein rhythm has not been confirmed. Furthermore, drug intake in mice is typically higher during the dark phase, therefore, individuals could be more vulnerable to insults during periods predisposed to drug seeking. This TOD vulnerability emphasizes the importance of considering rhythms in gene expression in future studies.

In order to characterize the striatal mechanisms underlying increased dark phase self-administration in female *Npas2* mutant mice, we identified DEGs in the NAc, DLS and DMS in cocaine-naïve mice. The NAc core and shell were pooled since cocaine-induced activation was increased in the core and, to a lesser extent, shell of mutants. As expected, we found overall changes in circadian rhythm-related genes; for example, *Arntl* was upregulated, likely to compensate for its inability to form a transcriptional complex with the mutated *Npas2. Fabp7* was the most highly upregulated DEG in *Npas2* mutants across all regions. It regulates fatty acid uptake in astrocytes, sleep (Gerstner et al., 2017) and drug seeking under stressful conditions (Hamilton et al., 2018). *Fabp7* is rhythmically expressed throughout the brain (Schnell et al., 2014) and is similarly upregulated in *Arntl* and *Nr1d1* knockout mice (Schnell et al., 2014; Gerstner & Paschos, 2020). *Fabp7* could be upregulated in *Npas2* mutants due to the downregulation of *Nr1d1*, which represses *Fabp7* (Schnell et al., 2014).

Although some DEGs were changed throughout the striatum, patterns typically varied by subregion. The DLS and NAc had the most similar pattern of DE, while the DLS/DMS and DMS/NAc had less overlap. Less significant DE was also found in the DMS (Table.3).These findings parallel cocaine-induced activation patterns and emphasize how the DMS is differentially affected by *Npas2* mutation.

**Table 3.**
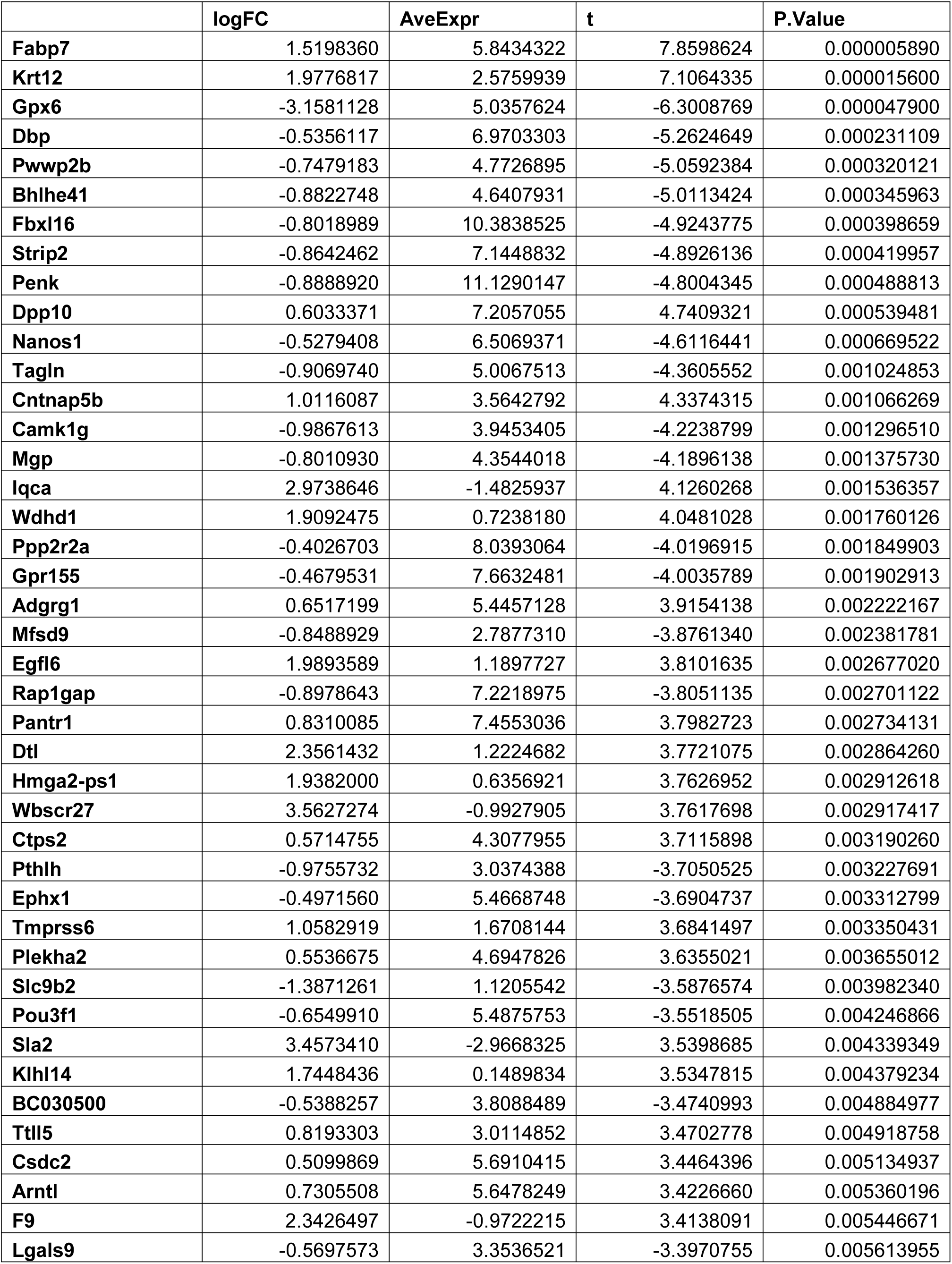
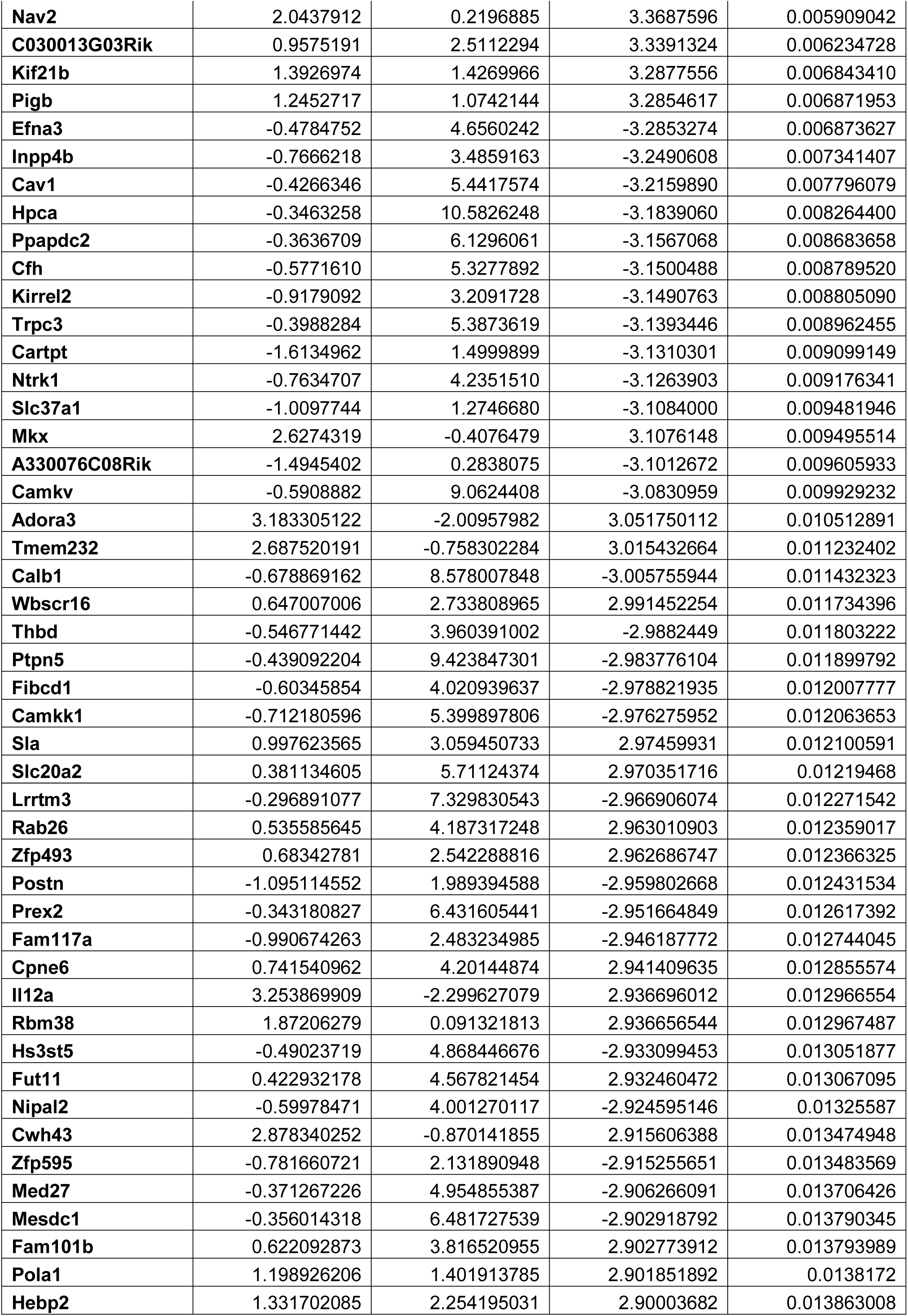
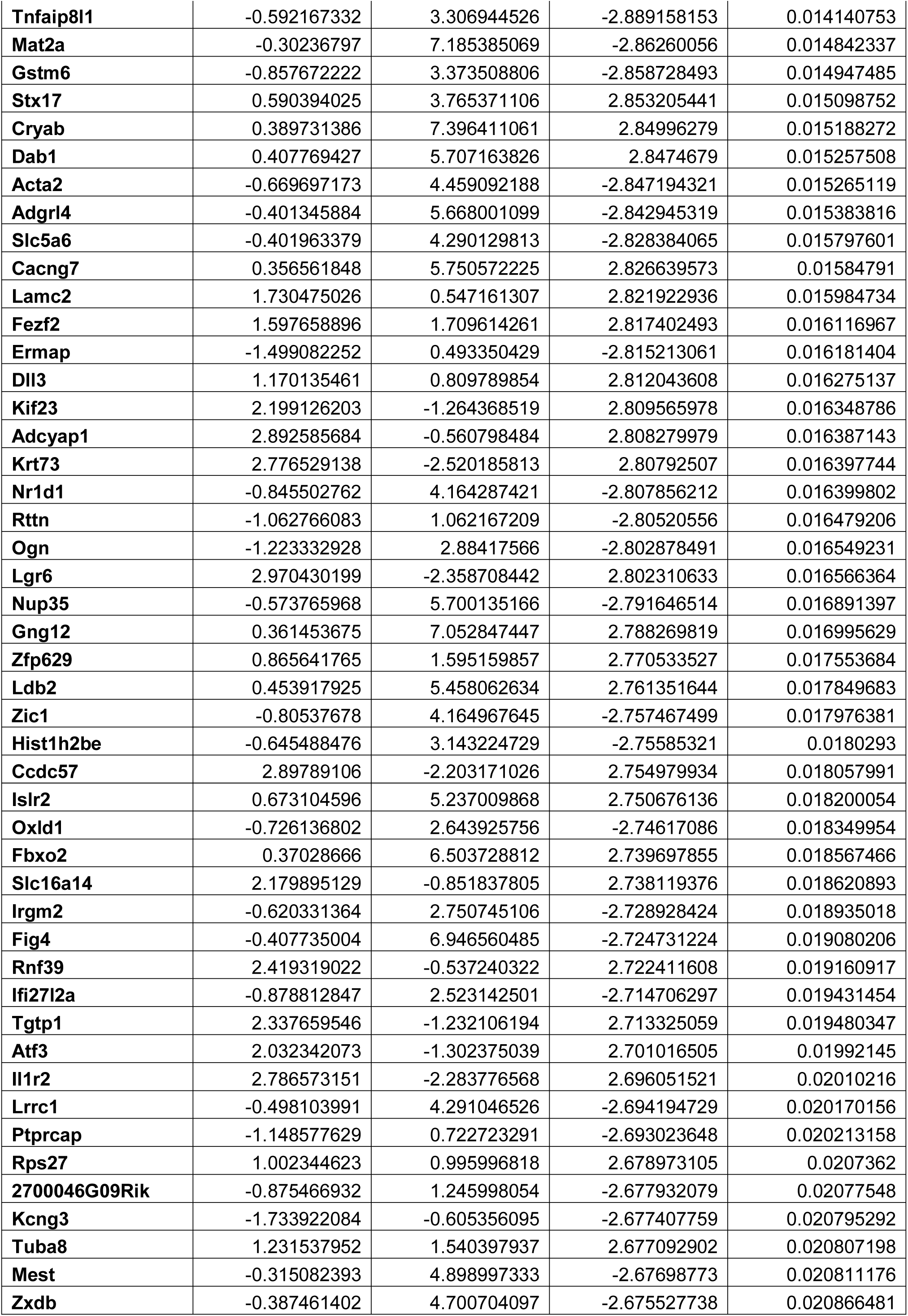
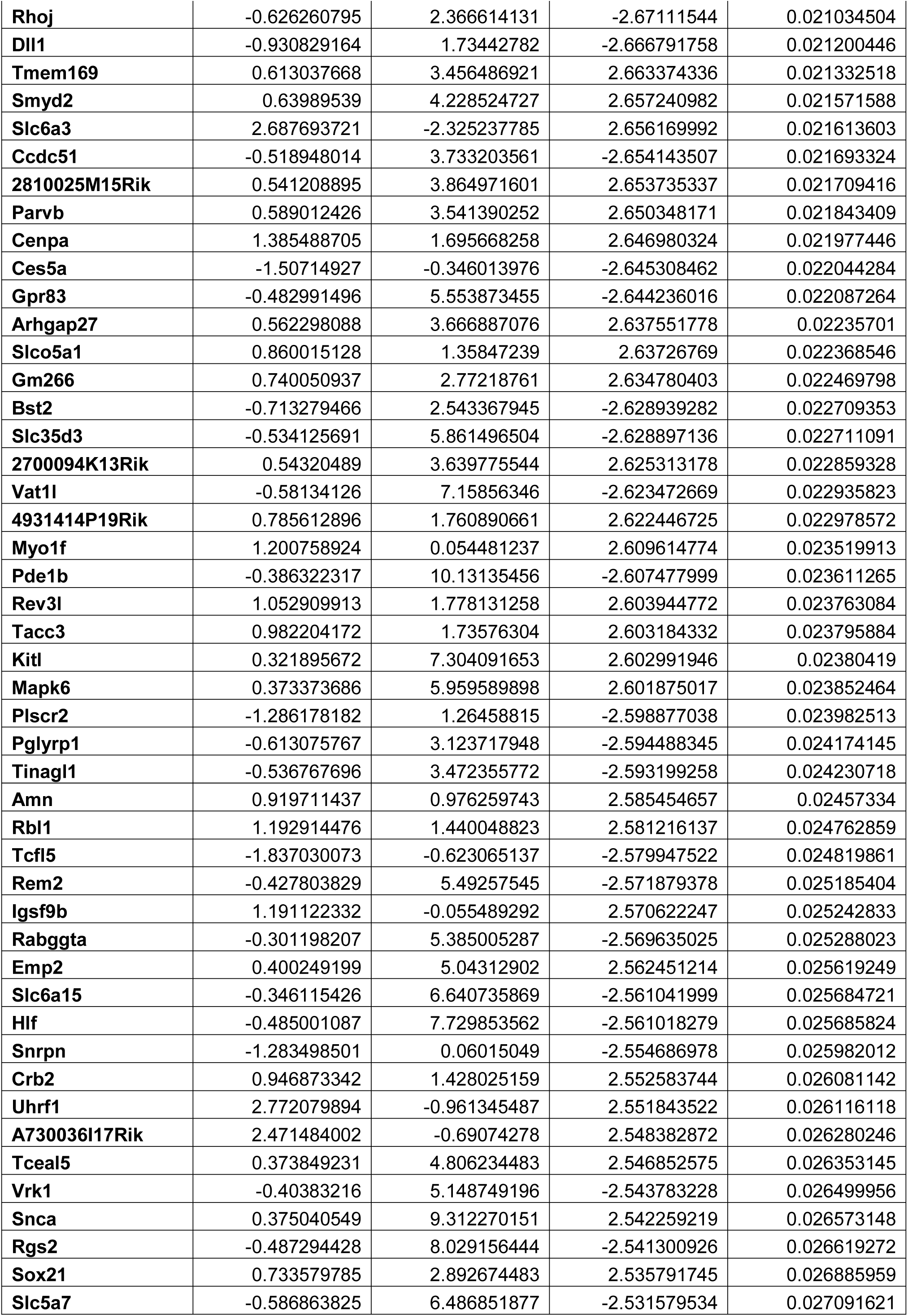
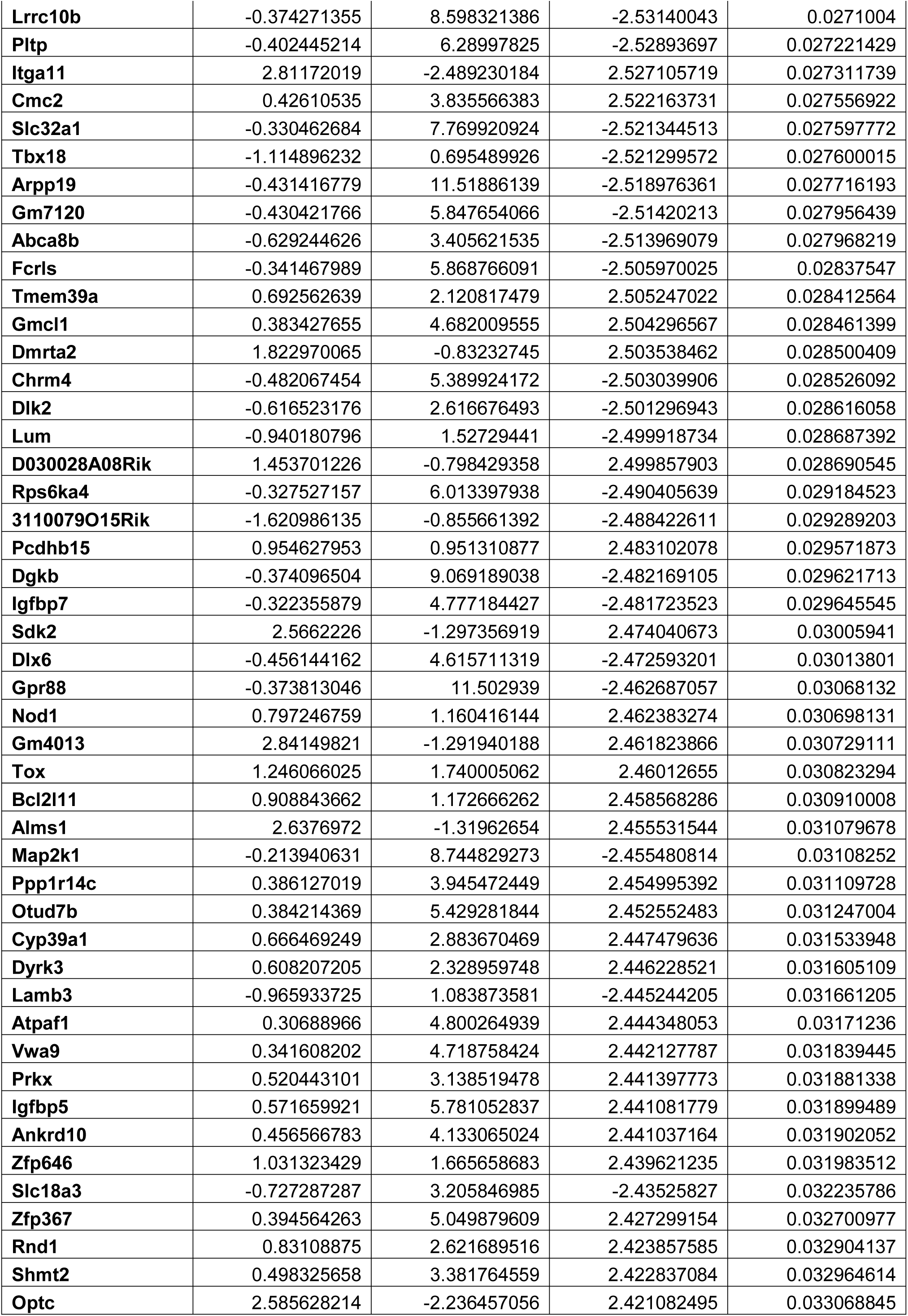
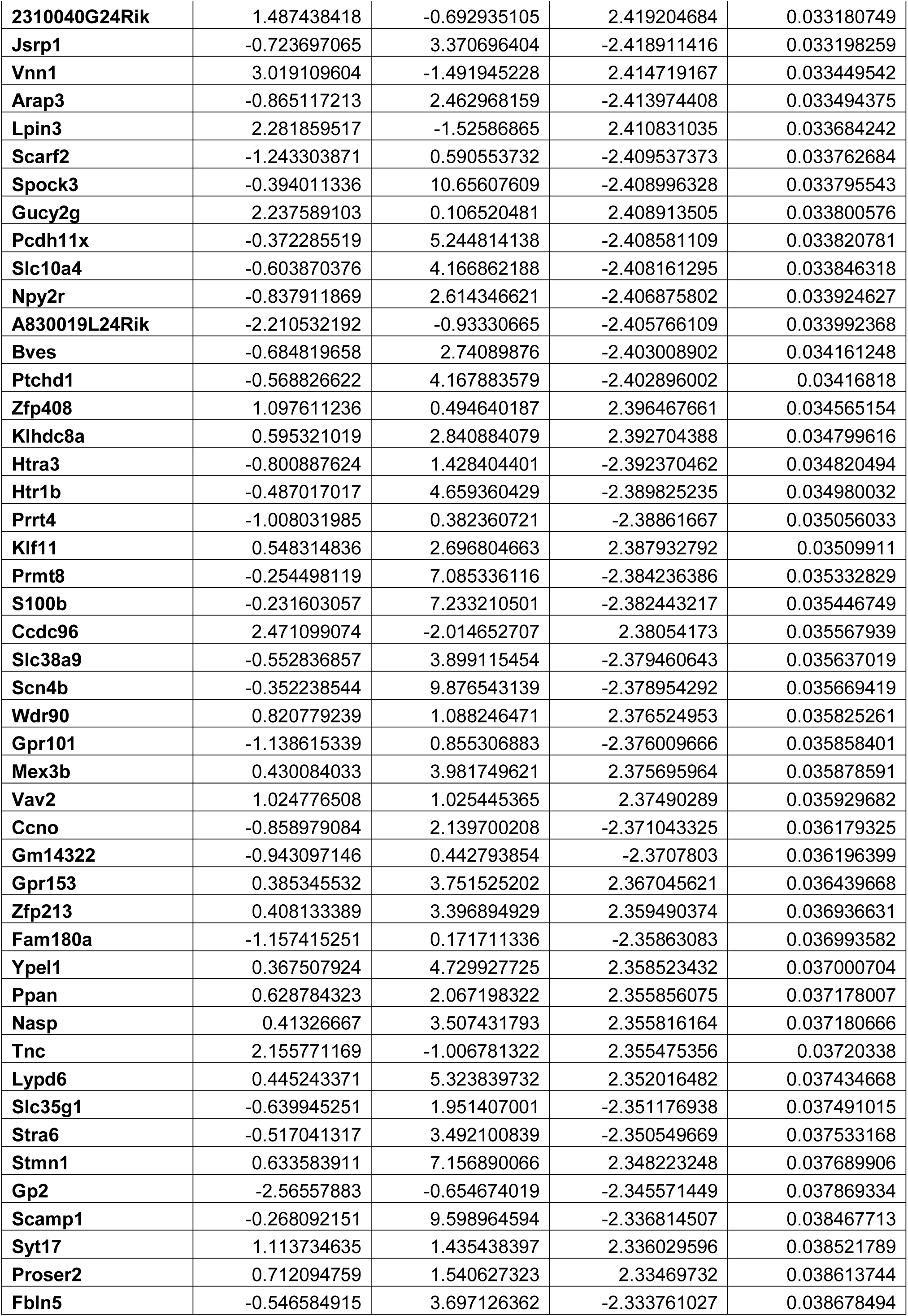
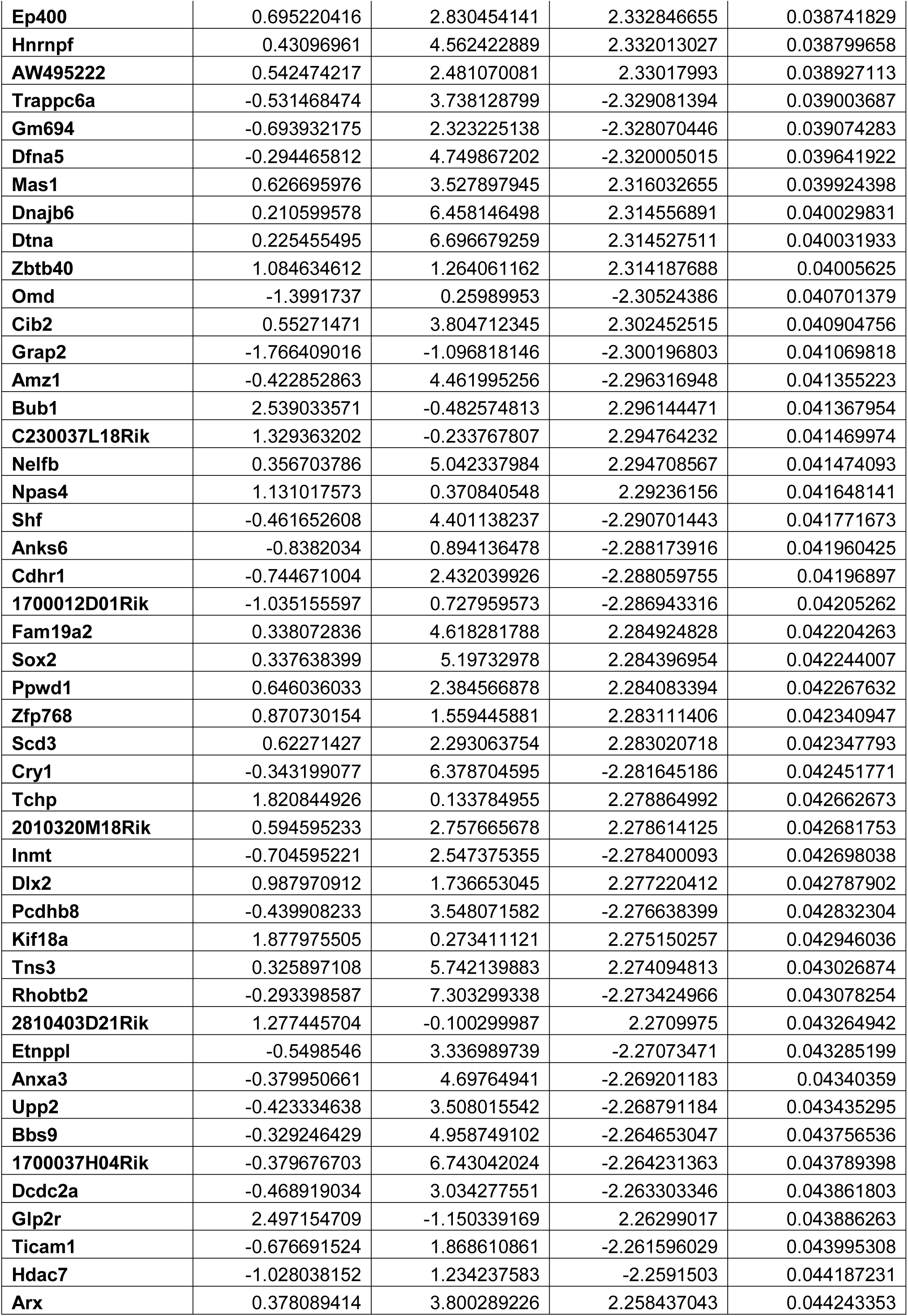
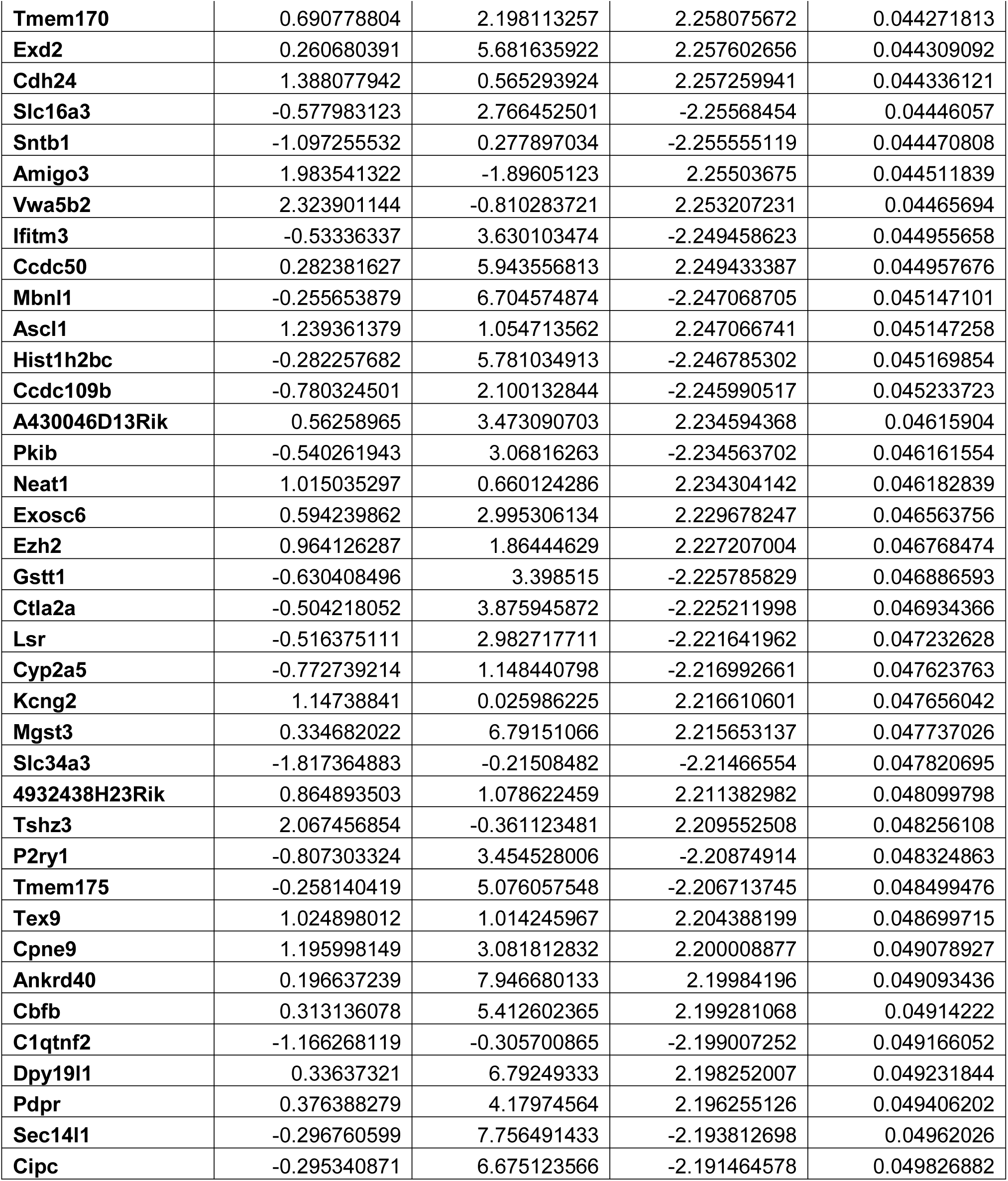
Differential expression analysis in the DMS for WT compared to Npas2 mutant females in the dark phase.

We next identified target genes in *Npas2* mutant females that might predispose increased cocaine self-administration. Using a meta-analysis (Huo et al., 2020), we identified 463 DEGs specific to the DLS and NAc where cocaine-induced activation is found in *Npas2* mutants. Many of these similarly regulated DEGs modulate neurotransmission and could therefore mediate increased glutamatergic transmission in the NAc of *Npas2* knockdown mice (Parekh et al., 2019), which could contribute to increased self-administration. These DEGs encode GABA receptor subunits, matrix metallopeptidases, small rho GTPases and synaptosomal proteins. GABAergic transmission is a promising target, since NPAS2 directly binds GABA receptor subunits (Ozburn et al., 2017). One of the most common DEG categories was potassium channels, which were frequently upregulated and could be modulating action potential generation and short-term plasticity in the DLS and NAc of mutant females. Importantly, DEGs from the NAc could have been diluted since the shell was activated to a lesser extent than the core. Overall, the DLS is most impacted by *Npas2* mutation with additional DEGs identified that regulate neurotransmission: other GTPase-related genes (*Arhgap9,Rasal1*), scaffolding genes (*Shank1*) and glutamate receptor subunits (*Gria4*).

Interestingly, enriched pathways in the NAc, DMS and DLS are also distinct. Circadian pathways were unsurprisingly identified in all regions (Dong et al., 2011; Baranger et al., 2016), since disruptions in circadian rhythms and transcription contribute to SU vulnerability. Importantly, circadian transcription factor (CLOCK,ARNTL) pathways were particularly significant in the NAc, suggesting a critical role in linking rhythm disruptions and drug use. Meanwhile, top pathways in the DLS suggest an increase in ΔFosB and CREB signaling at baseline, which could be exacerbated following self-administration. Other DLS pathways correspond to unique DEGs that mediate neurotransmission and synaptic plasticity. Future studies could examine the role of these genes in increased self-administration in *Npas2* mutants. For example, reducing ΔFosB-mediated transcription in *Npas2* mutants in D1+ cells in the DLS might normalize cocaine self-administration.

Ultimately, these results demonstrate that decreased function in NPAS2 drives vulnerability for SU in a circadian- and sex-dependent manner. Parallel changes in striatal activation and gene expression in *Npas2* mutant females in the dark phase suggest that gene expression changes, perhaps mediated by ΔFosB signaling, underlie behavioral vulnerabilities associated with mutating the key circadian transcription factor *Npas2*.

## Acknowledgements

We would like to thank Mariah Hildebrand and Laura Holesh for animal care and genotyping. We thank Drs. Steven McKnight and David Weaver for providing the *Npas2* mutant mice. This project used the University of Pittsburgh Health Sciences Sequencing Core at UPMC Children’s Hospital of Pittsburgh, (RNA sequencing). Cocaine was provided by NIDA via the NIH drug distribution center. This work was funded by the National Institutes of Health: DA039865, DA041872, DA039841, DA042886 (PI: Colleen McClung, PhD) and DA046117 (PI: Lauren DePoy).

